# A sex-specific evolutionary interaction between *ADCY9* and *CETP*

**DOI:** 10.1101/2021.05.12.443794

**Authors:** Isabel Gamache, Marc-André Legault, Jean-Christophe Grenier, Rocio Sanchez, Eric Rhéaume, Samira Asgari, Amina Barhdadi, Yassamin Feroz Zada, Holly Trochet, Yang Luo, Leonid Lecca, Megan Murray, Soumya Raychaudhuri, Jean-Claude Tardif, Marie-Pierre Dubé, Julie G. Hussin

**Affiliations:** Montreal Heart Institute, Montreal, Québec, Canada; Faculty of Medicine, Université de Montréal, Montreal, Quebec, Canada; Université de Montréal Beaulieu-Saucier Pharmacogenomics Centre, Montreal, Canada; Center for Data Sciences, Brigham and Women’s Hospital and Harvard Medical School, Boston, MA 02115, USA; Program in Medical and Population Genetics, Broad Institute of MIT and Harvard, Cambridge, MA 02115, USA; Socios En Salud, Lima, Peru; Department of Global Health and Social Medicine, Harvard Medical School; Centre for Genetics and Genomics Versus Arthritis, Manchester Academic Health Science Centre, University of Manchester, Manchester M13 9PL, UK; Department of Biomedical Informatics, Harvard Medical School, Boston, MA 02115, USA; Department of Medicine, Brigham and Women’s Hospital and Harvard Medical School, Boston, MA 02115, USA

## Abstract

Pharmacogenomic studies have revealed associations between rs1967309 in the adenylyl cyclase type 9 (*ADCY9*) gene and clinical responses to the cholesteryl ester transfer protein (CETP) modulator dalcetrapib, however, the mechanism behind this interaction is still unknown. Here, we characterized selective signals at the locus associated with the pharmacogenomic response in human populations and we show that rs1967309 region exhibits signatures of positive selection in several human populations. Furthermore, we identified a variant in *CETP*, rs158477, which is in long-range linkage disequilibrium with rs1967309 in the Peruvian population. The signal is mainly seen in males, a sex-specific result that is replicated in the LIMAA cohort of over 3,400 Peruvians. Analyses of RNA-seq data further suggest an epistatic interaction on *CETP* expression levels between the two SNPs in multiple tissues, which also differs between males and females. We also detected interaction effects of the two SNPs with sex on cardiovascular phenotypes in the UK Biobank, in line with the sex-specific genotype associations found in Peruvians at these loci. We propose that *ADCY9* and *CETP* coevolved during recent human evolution due to sex-specific selection, which points towards a biological link between dalcetrapib’s pharmacogene *ADCY9* and its therapeutic target *CETP*.

## Introduction

Coronary artery disease (CAD) is the leading cause of mortality worldwide. It is a complex disease caused by the accumulation of cholesterol-loaded plaques that block blood flow in the coronary arteries. The cholesteryl ester transfer protein (CETP) mediates the exchange of cholesterol esters and triglycerides between high-density lipoproteins (HDL) and lower density lipoproteins (1, 2). Dalcetrapib is a CETP modulator that did not reduce cardiovascular event rates in the overall dal-OUTCOMES trial of patients with recent acute coronary syndrome (3). However, pharmacogenomic analyses revealed that genotypes at rs1967309 in the *ADCY9* gene, coding for the ninth isoform of adenylate cyclase, modulated clinical responses to dalcetrapib (4). Individuals who carried the AA genotype at rs1967309 in *ADCY9* had less cardiovascular events, reduced atherosclerosis progression, and enhanced cholesterol efflux from macrophages when treated with dalcetrapib compared to placebo (4, 5). In contrast, those with the GG genotype had the opposite effects from dalcetrapib. Furthermore, a protective effect against the formation of atherosclerotic lesions was seen only in the absence of both *Adcy9* and *CETP* in mice (6), suggesting an interaction between the two genes. However, the underlying mechanisms linking *CETP* and *ADCY9,* located 50 Mb apart on chromosome 16, as well as the relevance of the rs1967309 non-coding genetic variant are still unclear.

Identification of selection pressure on a genetic variant can help shed light on its importance. Adaptation to different environments often leads to a rise in frequency of variants, by favoring survival and/or reproduction fitness. An example is the lactase gene (*LCT*) (7–11), where a positively selected intronic variant in *MCM6* leads to an escape from epigenetic inactivation of *LCT* and facilitates lactase persistence after weaning (12). Results of genomic studies for phenotypes such as adaptation to high-altitude hypoxia in Tibetans (13), fatty acid metabolism in Inuits (14) or response to pathogens across populations (15) have also been confirmed by functional studies (16–20). Thus, population and regulatory genomics can be leveraged to unveil the effect of genetic mutations at a single non-coding locus and reveal the biological mechanisms of adaptation.

When two or more loci interact during adaptation, a genomic scan will likely be underpowered to pinpoint the genetic determinants. In this study, we took a multi-step approach on the *ADCY9* and *CETP* candidate genes to specifically study their interaction (Figure 1). We used a joint evolutionary analysis to evaluate the potential signatures of selection in these genes (Step 1), which revealed positive selection pressures acting on *ADCY9.* Sex-specific genetic associations between the two genes are discovered in Peruvians (Step 2), a population in which natural selection for high-altitude was previously found on genes related to cardiovascular health (21). Furthermore, our know-down experiments and analyses of large-scale transcriptomics (Step 3) as well as available phenome-wide resources (Step 4) bring further evidence of a sex-specific epistatic interaction between *ADCY9* and *CETP*.

**Figure 1.**
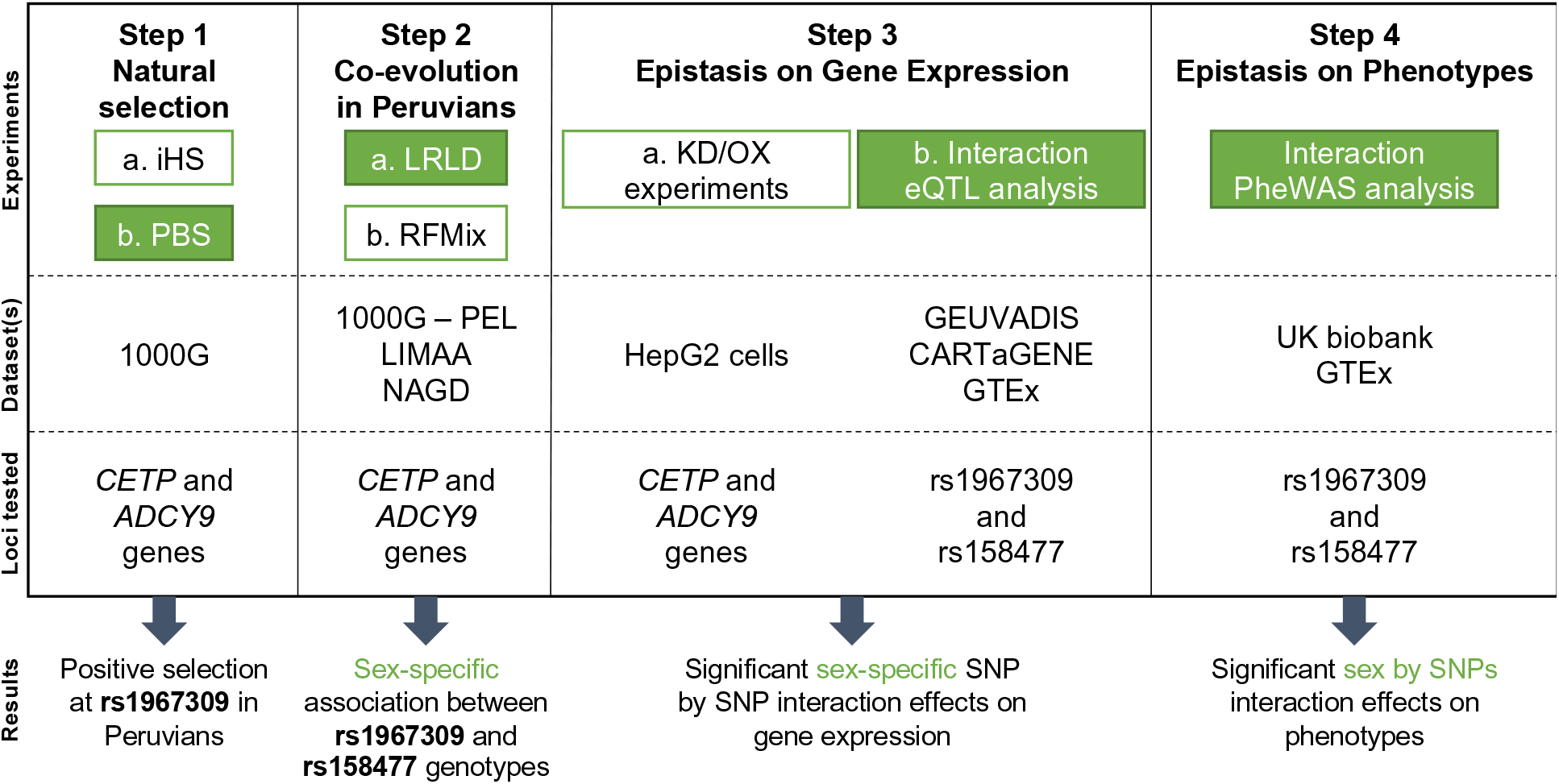
Flowchart of experimental design and main results. The four main steps of the analyses conducted in this study are reported along with the datasets used for each step and the genetic loci on which the analyses are performed. Green colored boxes represent analyses for which sex is considered. KD = Knock-down OX = Overexpression

## Results

### Signatures of selection at rs1967309 in *ADCY9* in human populations

The genetic variant rs1967309 is located in intron 2 of *ADCY9*, in a region of high linkage disequilibrium (LD), in all subpopulations in the 1000 Genomes Project (1000G), and harbors heterogeneous genotype frequencies across human populations (Figure 2a). Its intronic location makes it difficult to assess its functional relevance but exploring selective signals around intronic SNPs in human populations can shed light on their importance. In African populations (AFR), the major genotype is AA, which is the homozygous genotype for the ancestral allele, whereas in Europeans (EUR), AA is the minor genotype. The frequency of the AA genotype is slightly higher in Asia (EAS, SAS) and America (AMR) compared to that in Europe, becoming the most frequent genotype in the Peruvian population (PEL). Using the integrated haplotype score (iHS) (22) (Step 1a, Figure 1), a statistic that enables the detection of evidence for recent strong positive selection (typically when |iHS| > 2), we observed that several SNPs in the LD block around rs1967309 exhibit selective signatures in non-African populations (|iHS_SAS_|= 2.66, |iHS_EUR_|= 2.31), whereas no signal is seen in this LD block in African populations (Figure 2b, Supplementary Figure 1, Supplementary text). Our analyses suggest that this locus in *ADCY9* has been the target of recent positive selection in several human populations, with multiple, possibly independent, selective signals detectable around rs1967309. However, recent positive selection as measured by iHS does not seem to explain the notable increase in frequency for the A allele in the PEL population (f_A_=0.77), compared to the European (f_A_=0.41), Asian (f_A_=0.44), and other American populations (f_A_=0.54 in AMR without PEL).

**Figure 2.**
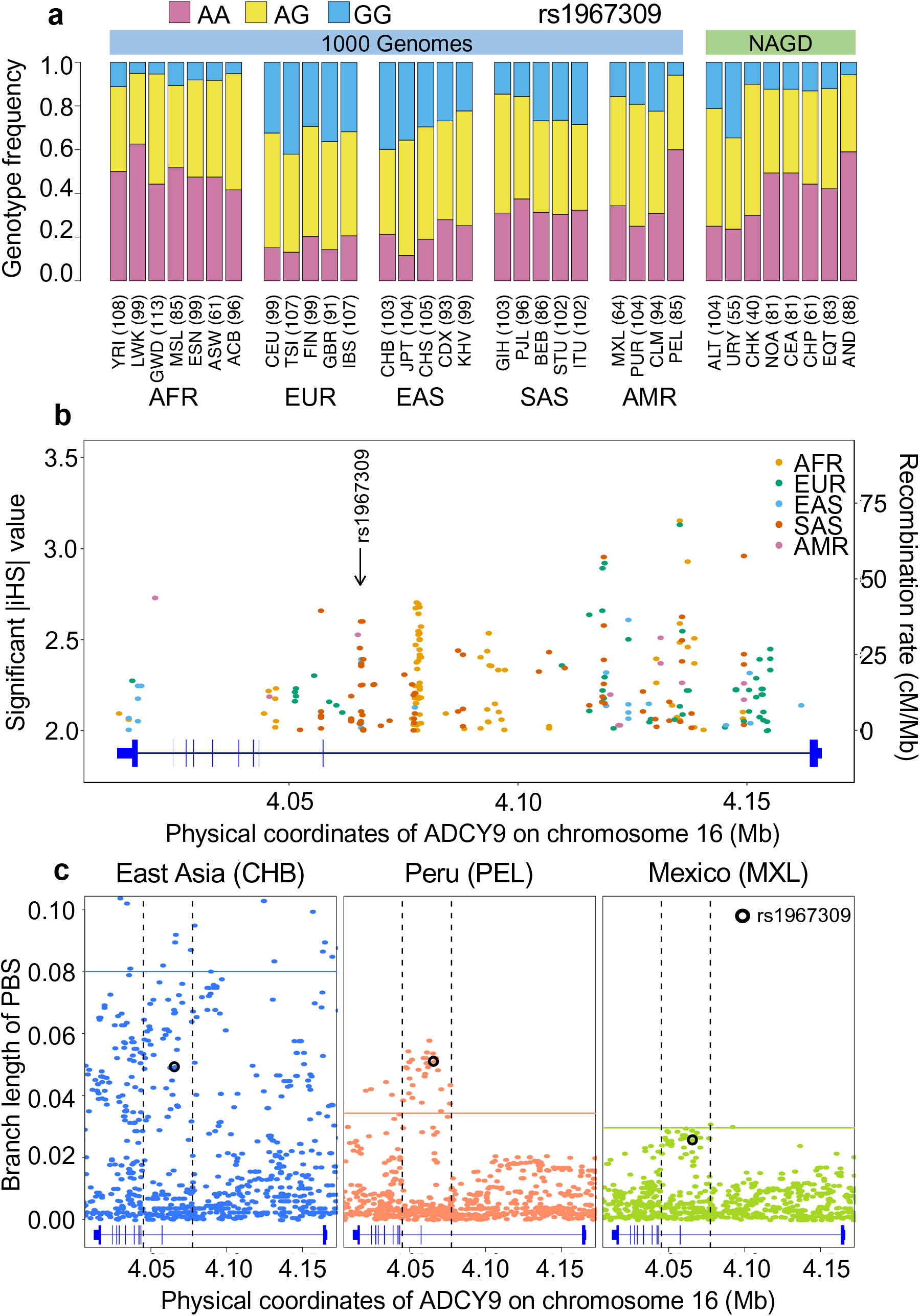
Natural selection signature at rs1967309 in *ADCY9*. (a) Genotype frequency distribution of rs1967309 in populations from the 1000 Genomes (1000G) Project and in Native Americans. (b) Significant iHS values (absolute values above 2) for 1000G continental populations and recombination rates from AMR-1000G population-specific genetic maps, in the *ADCY9* gene. (c) PBS values in the *ADCY9* gene, in CHB (outgroup, left panel), PEL (middle panel) and MXL (right panel). Horizontal lines represent the 95^th^ percentile PBS value genome-wide for each population. Vertical dotted black lines define the LD block around rs1967309 (black circle) from 1000G population-specific genetic maps. Gene plots for *ADCY9* showing location of its exons are presented in blue below each plot. Abbreviations: Altaic from Mongolia and Russia: ALT; Uralic Yukaghir from Russia: URY; Chukchi Kamchatkan from Russia: CHK; Northern American from Canada, Guatemala and Mexico: NOA; Central American from Costal Rica and Mexico: CEA; Chibchan Paezan from Argentina, Bolivia, Colombia, Costa Rica and Mexico: CHP; Equatorial Tucanoan from Argentina, Brazil, Colombia, Gualana and Paraguay: EQT; Andean from Bolivia, Chile, Colombia and Peru: AND. For 1000G populations, abbreviations can be found here https://www.internationalgenome.org/category/population/.

To test whether the difference between PEL and other AMR allele frequencies at rs1967309 is significant, we used the population branch statistic (PBS) (Step 1b, Figure 1). This statistic has been developed to locate selection signals by summarizing differentiation between populations using a three-way comparison of allele frequencies between a specific group, a closely related population, and an outgroup (13). It has been shown to increase power to detect incomplete selective sweeps on standing variation. Applying this statistic to investigate rs1967309 allele frequency in PEL, we used Mexicans (MXL) as a closely related group and a Chinese population (CHB) as the outgroup (Methods). Over the entire genome, the CHB branches are greater than PEL and MXL branches (mean_CHB_=0.020, mean_MXL_=0.008, mean_PEL_=0.009), which reflects the expectation under genetic drift. However, the estimated PEL branch length at rs1967309 (Figure 2c), which reflects differentiation since the split from the MXL population (PBS_PEL,rs1967309_=0.051, empirical p-value = 0.014), surpasses the CHB branch length (PBS_CHB,rs1967309_=0.049, empirical p-value > 0.05), which reflects differentiation since the split between Asian and American populations, whereas no such effect is seen in MXL (PBS_MXL,rs1967309_=0.026, empirical p-value > 0.05), or for any other AMR populations. Furthermore, the PEL branch lengths at several SNPs in this LD block (Figure 2c) are in the top 5% of all PEL branch lengths across the whole genome (PBS_PEL,95th_ = 0.031), whereas these increased branch lengths are not observed outside of the LD block (Figure 2c). These results are robust to the choice of the outgroup and the closely related AMR population (Methods).

The increase in frequency of the A allele at rs1967309 is also seen in genotype data from Native American populations (23), with Andeans showing genotype frequencies highly similar to PEL (f_A_=0.77, Figure 2a). The PEL population has a large Andean ancestry (Methods, Supplementary Figure 2a,b) and almost no African ancestry, strongly suggesting that the increase in AA genotype arose in the Andean population and not from admixture with Africans. The PEL individuals that harbor the AA genotype for rs1967309 do not exhibit a larger genome-wide Andean ancestry than non-AA individuals (p-value=0.30, Mann-Whitney U test). Overall, these results suggest that the ancestral allele A at rs1967309, after dropping in frequency following the out-of-Africa event, has increased in frequency in the Andean population and has been preferentially retained in the Peruvian population’s genetic makeup, potentially because of natural selection.

### Evidence for co-evolution between *ADCY9* and *CETP* in Peru

The pharmacogenetic link between *ADCY9* and the CETP modulator dalcetrapib raises the question of whether there is a genetic relationship between rs1967309 in *ADCY9* and *CETP*, both located on chromosome 16. Such a relationship can be revealed by analyzing patterns of long-range linkage disequilibrium (LRLD) (24, 25), in order to detect whether specific combinations of alleles (or genotypes) at two loci are particularly overrepresented. To do so, we calculated the genotyped-based linkage disequilibrium (r^2^) (Step 2a, Figure 1) between rs1967309 and each SNP in *CETP* with minor allele frequency (MAF) above 0.05. In the Peruvian population, there are four SNPs, (including 2 in perfect LD in PEL) that exhibit r^2^ values with rs1967309 that are in the top 1% of r^2^ values (Figure 3a) computed for all 37,802 pairs of SNPs in *ADCY9* and *CETP* genes with MAF>0.05 (Methods). Despite the r^2^ values themselves being low (r^2^_s158477_=0.080, r^2^_rs158480;rs158617_=0.089, r^2^_rs12447620_=0.090), these values are highly unexpected for these two genes situated 50 Mb apart (*ADCY9/CETP* empirical p-value<0.006, Supplementary Table 1) and thus correspond to a significant LRLD signal. This signal is not seen in other 1000G populations (Supplementary Table 1). We also computed r^2^ between the four identified SNPs’ genotypes and all *ADCY9* SNPs with MAF above 0.05 (Figure 3b). The distribution of r^2^ values for the rs158477 *CETP* SNP shows a clear bell-shaped pattern around rs1967309 in *ADCY9*, which strongly suggests the rs1967309-rs158477 genetic association detected is not simply a statistical fluke, while the signal in the region for the other SNPs is less conclusive. The SNP rs158477 in *CETP* is also the only one that has a PEL branch length value higher than the 95^th^ percentile, also higher than the CHB branch length value (PBS_PEL,rs158477_= 0.062, Supplementary Figure 3a), in line with the observation at rs1967309. Strikingly, this *CETP* SNP’s genotype frequency distribution across the 1000G and Native American populations resembles that of rs1967309 in *ADCY9* (Figure 3c). Given that the Peruvian population is admixed (26), particular enrichment of genome segments for a specific ancestry, if present, would lead to inflated LRLD between these segments (27–30), we thus performed several admixture-related analyses (Step 2b, Figure 1). No significant enrichment is seen at either locus and significant LRLD is also seen in the Andean source population (Figure 3-figure supplement 1a,b, Supplementary text). Furthermore, we see no enrichment of Andean ancestry in individuals harboring the overrepresented combination of genotypes, AA at rs1967309 + GG at rs158477, compared to other combinations (p-value=0.18, Mann-Whitney U test). These results show that admixture patterns in PEL cannot be solely responsible for the association found between rs1967309 and rs158477. Finally, using a genome-wide null distribution which allows to capture the LRLD distribution expected under the admixture levels present in this sample (Supplementary text), we show that the r^2^ value between the two SNPs is higher than expected given their allele frequencies and the physical distance between them (genome-wide empirical p-value=0.01, Figure 3d). Taken together, these findings strongly suggest that the AA/GG combination is being transmitted to the next generation more often (ie. is likely selectively favored) which reveals a signature of co-evolution between *ADCY9* and *CETP* at these loci.

**Figure 3.**
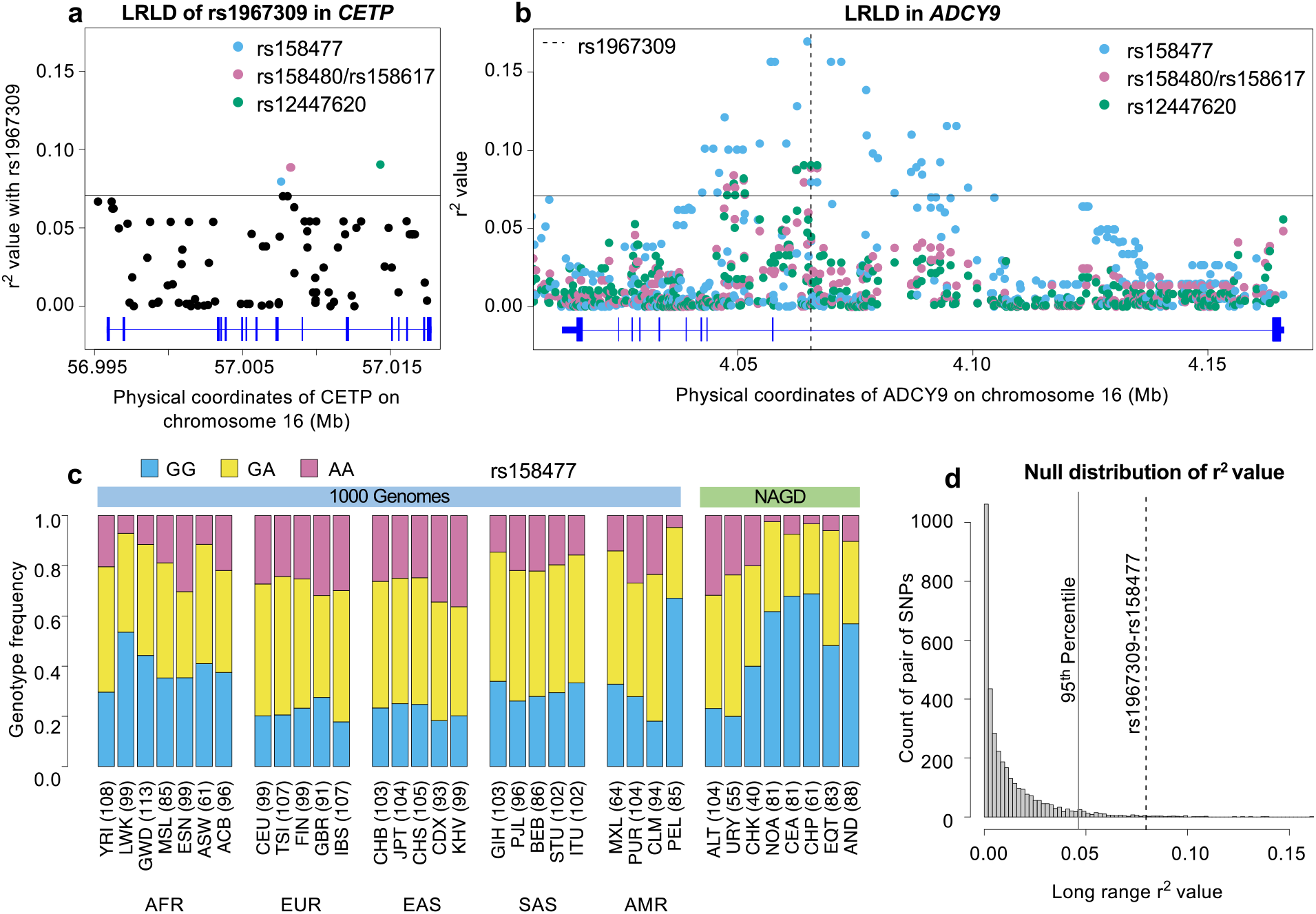
Long-range linkage disequilibrium between rs1967309 and rs158477 in Peruvians from Lima, Peru. (a) Genotype correlation (r^2^) between rs1967309 and all SNPs with MAF>5% in *CETP*, for the PEL population. (b) Genotype correlation between the 3 loci identified in (a) to be in the 99^th^ percentile and all SNPs with MAF>5% in *ADCY9*. The dotted line indicates the position of rs1967309. The horizontal lines in (a,b) represent the threshold for the 99^th^ percentile of all comparisons of SNPs (MAF>5%) between *ADCY9* and *CETP*. Figure 3-figure supplement 1 presents the same plots for Andeans and in the replication cohort (LIMAA) and Figure 3-figure supplement 2 compares the r^2^ values between PEL and LIMAA (c) Genotype frequency distribution of rs158477 in 1000G and Native American populations. (d) Genomic distribution of r^2^ values from 3,513 pairs of SNPs separated by between 50-60 Mb and 61±10 cM away across all Peruvian chromosomes from the PEL sample, compared to the rs1967309-rs158477 r^2^ value (dotted grey line) (genome-wide empirical p-value=0.01). The vertical black line shows the threshold for the 95^th^ percentile threshold of all pairs. Gene plots showing location of exons for *CETP* (a) and *ADCY9* (b) are presented in blue below each plot. Abbreviations: Altaic from Mongolia and Russia: ALT; Uralic Yukaghir from Russia: URY; Chukchi Kamchatkan from Russia: CHK; Northern American from Canada, Guatemala and Mexico: NOA; Central American from Costal Rica and Mexico: CEA; Chibchan Paezan from Argentina, Bolivia, Colombia, Costa Rica and Mexico: CHP; Equatorial Tucanoan from Argentina, Brazil, Colombia, Gualana and Paraguay: EQT; Andean from Bolivia, Chile, Colombia and Peru: AND. For 1000G populations, abbreviations can be found here https://www.internationalgenome.org/category/population/

Still, such a LRLD signal can be due to a small sample size (29). To confirm independently the association between genotypes at rs1967309 of *ADCY9* and rs158477 of *CETP*, we used the LIMAA cohort (31, 32), a large cohort of 3,509 Peruvian individuals with genotype information, to replicate our finding. The ancestry distribution, as measured by RFMix (Methods) is similar between the two cohorts (Supplementary Figure 2a,b), however, the LIMAA cohort population structure shows additional subgroups compared to the 1000G PEL population sample (Supplementary Figure 2c-e): to limit confounders, we excluded individuals coming from these subgroups (Supplementary text). In this dataset (N=3,243), the pair of SNPs rs1967309-rs158477 is the only pairs identified in PEL who shows evidence for LRLD, with an r^2^ value in the top 1% of all pairs of SNPs in *ADCY9* and *CETP* (*ADCY9/CETP* empirical p-value=0.003, Figure 3-figure supplement 1c,d, Figure 3-figure supplement 2, Supplementary Table 1). The r^2^ test used above is powerful to detect allelic associations, but the net association measured will be very small if selection acts on a specific genotype combination rather than on alleles. In that scenario, and when power allows it, the genotypic association is better assessed by with a *χ*^2^ distributed test statistic (with four degrees of freedom, 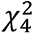) comparing the observed and expected genotype combination counts (25). The test confirmed the association in LIMAA (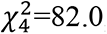, permutation p-value <0.001, genome-wide empirical p-value=0.0003, Supplementary text). The association discovered between rs1967309 and rs158477 is thus generalizable to the Peruvian population and not limited to the 1000G PEL sample.

### Sex-specific long-range linkage disequilibrium signal

Because the allele frequencies at rs1967309 were suggestively different between males and females (Figure 4-figure supplement 1), we performed sex-stratified PBS analyses, which suggested that the LD block around rs1967309 is differentiated between sexes in the Peruvians (Figure 4-figure supplement 2, Supplementary text). We therefore explored further the effect of sex on the LRLD association found between rs1967309 and rs158477 and performed sex-stratified LRLD analyses. These analyses revealed that the correlation between rs1967309 and rs158477 is only seen in males in PEL (Figure 4a,b, Supplementary Figure 4a,b, Supplementary Table 1): the r^2^ value rose to 0.348 in males (*ADCY9/CETP* empirical p-value=8.23×10^-5^, genome-wide empirical p-value<2.85 x 10^-4^, N=41) and became non-significant in females (*ADCY9*/*CETP* empirical p-value=0.78, genome-wide empirical p-value=0.80, N=44). In the Andean population, the association between rs1967309 and rs158477 is not significant when we stratified by sex (Supplementary Table 1), but we still see significant association signals with rs158477 at other SNPs in *ADCY9* LD block in both sexes (Figure 4-figure supplement 3). The LRLD result in PEL cannot be explained by differences of Andean ancestry proportion between males and females (p-value=0.27, Mann-Whitney U test). A permutation analysis that shuffled the sex labels of samples established that the observed difference between the sexes is larger than what we expect by chance (p-value=0.002, Supplementary Figure 4c, Supplementary text). In the LIMAA cohort, we replicate this sex-specific result (Figure 4c,d, Supplementary Table 1) where the r^2^ test is significant in males (*ADCY9*/*CETP* empirical p-value=0.003, N=1,941) but not in females (*ADCY9/CETP* empirical p-value=0.52, N=1,302). The genotypic *χ*^2^ test confirms the association between *ADCY9* and *CETP* is present in males (*χ*^2^ = 56.6, permutation p-value=0.001, genome-wide empirical p-value=0.002, Supplementary text), revealing an excess of rs1967309-AA + rs158477-GG. This is also the genotype combination driving the LRLD in PEL. In females, the test also shows a weaker but significant effect (*χ*^2^ = 37.0, permutation p-value = 0.017, genome-wide empirical p-value = 0.001) driven by an excess of a different genotype combination, rs1967309-AA + rs158477-AA, which is, however, not replicated in PEL possibly because of lack of power (Supplementary text).

**Figure 4.**
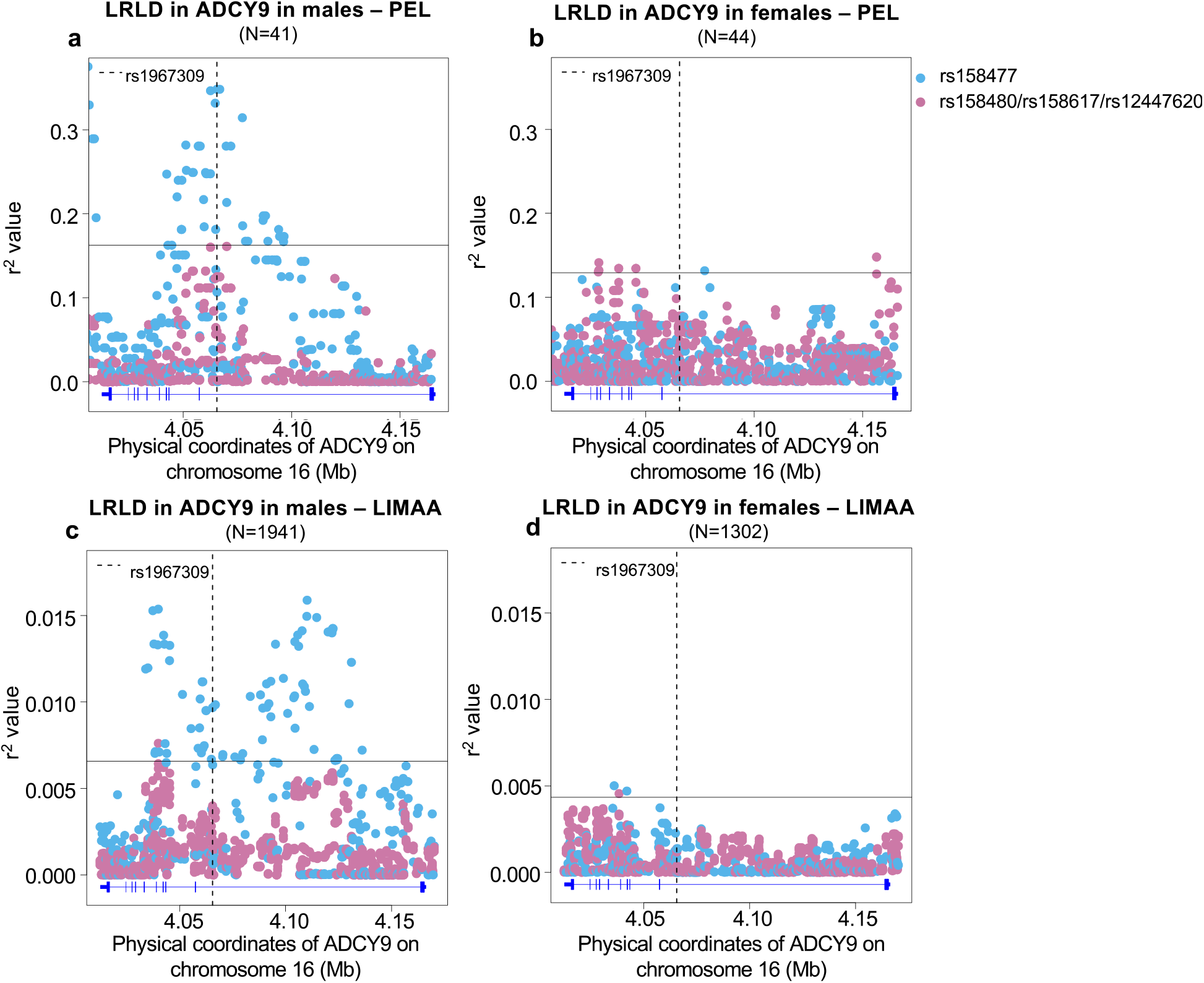
Sex-specific long-range linkage disequilibrium. Genotype correlation between the loci identified in *CETP* in Figure 3a and all SNPs with MAF>5% in *ADCY9* for (a,b) the PEL population and (c,d) LIMAA cohort in males (a,c) and in females (b,d). Genotype frequencies per sex are shown in Figure 4-figure supplement 1 and sex-specific PBS values in Figure 4-figure supplement 2. The horizontal line shows the threshold for the 99^th^ percentile of all comparisons of SNPs (MAF>5%) between *ADCY9* and *CETP*. The vertical dotted line represents the position of rs1967309. Blue dots represent the rs158477 SNPs and pink represents the other three SNPs identified in Figure 3a (rs158480, rs158617 and rs12447620), which are in near-perfect LD. Figure 4-figure supplement 3 shows the same analysis in Andeans from NAGD. Gene plots for *ADCY9* showing location of its exons are presented in blue below each plot.

### Epistatic effects on *CETP* gene expression

LRLD between variants can suggest the existence of gene-gene interactions, especially if they are functional variants (29). In order to be under selection, mutations typically need to modulate a phenotype or an endophenotype, such as gene expression. We have shown previously (6) that CETP and *Adcy9* interact in mice to modulate several phenotypes, including atherosclerotic lesion development. To test whether these genes interact in humans, we knocked down (KD) *ADCY9* in hepatocyte HepG2 cells (Step 3a, Figure 1) and performed RNA sequencing on five KD biological replicates and five control replicates, to evaluate the impact of decreased *ADCY9* expression on the transcriptome. We confirmed the KD was successful as *ADCY9* expression is reduced in the KD replicates (Figure 5a), which represents a drastic drop in expression compared to the whole transcriptome changes (False Discovery Rate [FDR] = 4.07 x 10^-14^, Methods). We also observed that *CETP* expression was increased in *ADCY9-KD* samples compared to controls (Figure 5a), an increase that is also transcriptome-wide significant (FDR=1.97 x 10^-7^, ß = 1.257). This increased expression was validated by qPCR, and western blot also showed increased CETP protein product (Methods, Figure 5-figure supplement 1a,b, Supplementary text), but its overexpression did not significatively modulate *CETP* expression (Figure 5-figure supplement 1c). Knocking down or overexpressing *CETP* did not impact *ADCY9* expression on qPCR (Figure 5-figure supplement 1 d,e). These experiments demonstrate an interaction between *ADCY9* and *CETP* at the gene expression level and raised the hypothesis that *ADCY9* potentially modulates the expression of *CETP* through a genetic effect mediated by rs1967309.

**Figure 5.**
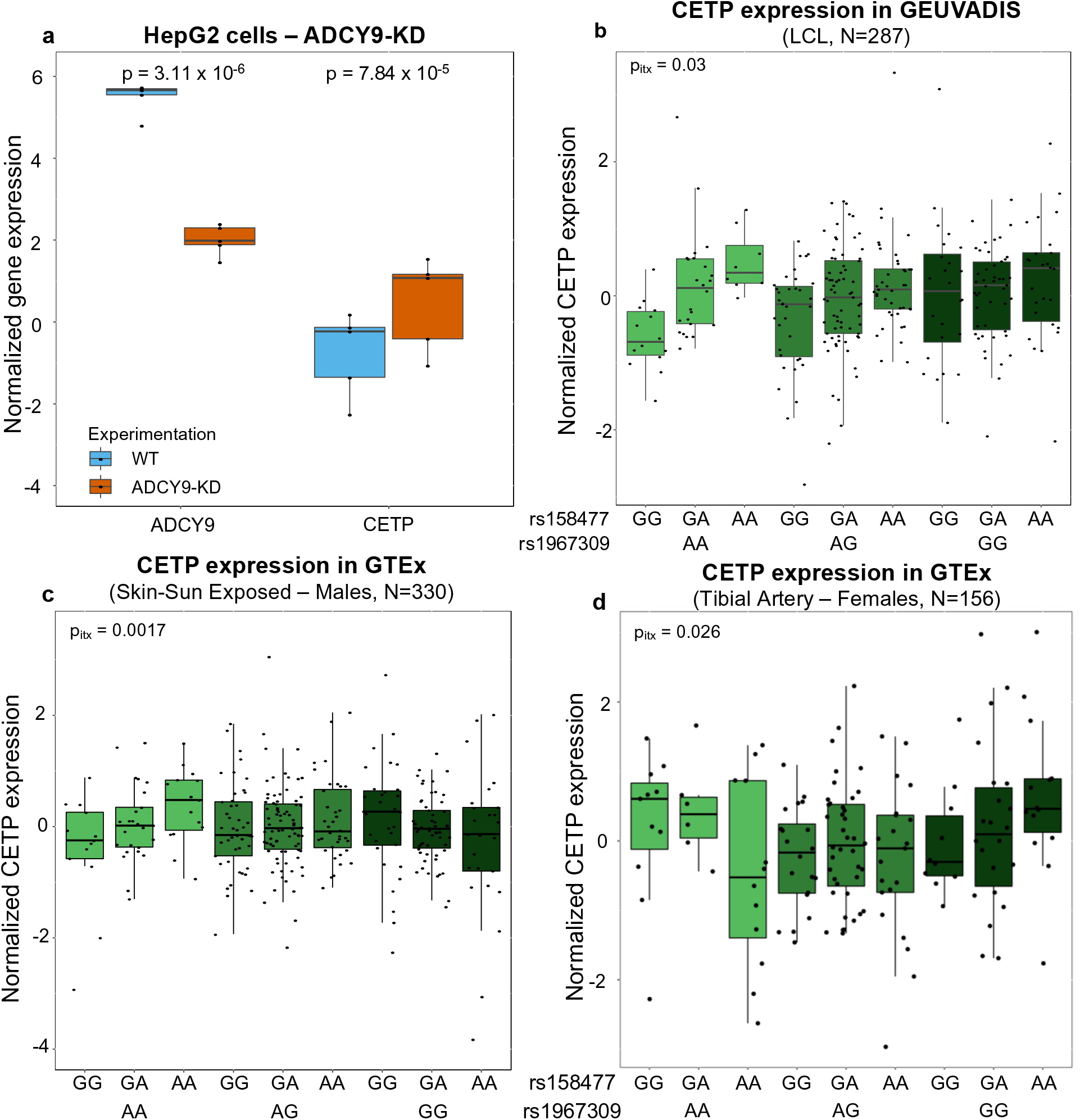
Effect of *ADCY9* on *CETP* expression. (a) Normalized expression of *ADCY9* or *CETP* genes depending on wild type (WT) and *ADCY9-KD* in HepG2 cells from RNA sequencing on five biological replicates in each group. P-values were obtained from a two-sided Wilcoxon paired test. qPCR and western blot results in HepG2 are presented in Figure 5-figure supplement 1. (b,c,d) *CETP* expression depending on the combination of rs1967309 and rs158477 genotypes in (b) GEUVADIS (p-value=0.03, ß= -0.22, N=287), (c) GTEx-Skin Sun Exposed in males (p-value=0.0017, ß= -0.32, N=330) and in (d) GTEx-Tibial artery in females (p-value=0.026, ß= 0.38, N=156), for individuals of European descent according to principal component analysis. P-values reported were obtained from a two-way interaction of a linear regression model for the maximum number of PEER/sPEER factors considered. Figure 5-figure supplement 2 show the interaction p-values depending on number of PEER/sPEER factors included in the linear models.

To test for potential interaction effects between rs1967309 and *CETP*, we used RNA-seq data from diverse projects in humans: the GEUVADIS project (33), the Genotype-Tissue Expression (GTEx v8) project (34) and CARTaGENE (CaG) (35) (Step 3b, Figure 1). When looking across tissues in GTEx, *ADCY9* and *CETP* expressions negatively correlate in almost all the tissues (Supplementary Figure 6, Supplementary text), which is consistent with the effect observed during the *ADCY9-KD* experiment, showing increased expression of *CETP* expression when *ADCY9* is lowly expressed (Figure 5a, Figure 5-figure supplement 1a,b). We evaluated the effects of the SNPs on expression levels of *ADCY9* and *CETP* by modelling both SNPs as continuous variables (additive model) (Methods). The *CETP* SNP rs158477 was reported as an expression quantitative trait locus (eQTL) in GTEx v7 and, in our models, shows evidence of being a *cis* eQTL of *CETP* in several other tissues (Supplementary text), although not reaching genome-wide significance. To test specifically for an epistatic effect between rs1967309 and rs158477 on *CETP* expression, we included an interaction term in eQTL models (Methods). We note here that we are testing for association for this specific pair of SNPs only, and that effects across tissues are not independent, such that we set our significance threshold at p-value=0.05. This analysis revealed a significant interaction effect (p-value=0.03, ß= -0.22) between the two SNPs on *CETP* expression in GEUVADIS lymphoblastoid cell lines (Figure 5b, Supplementary Figure 7a). In rs1967309 AA individuals, copies of the rs158477 A allele increased *CETP* expression by 0.46 (95% CI 0.26-0.86) on average. In rs1967309 AG individuals, copies of the rs158477 A allele increased CETP expression by 0.24 (95% CI 0.06-0.43) on average and the effect was null in rs1967309 GG individuals (p-value_GG_=0.58). This suggests that the effect of rs158477 on *CETP* expression changes depending on genotypes of rs1967309. The interaction is also significant in several GTEx tissues, most of which are brain tissues, like hippocampus, hypothalamus and substantia nigra, but also in skin, although we note that the significance of the interaction depends on the number of PEER factors included in the model (Supplementary Figure 8). These factors are needed to correct for unknown biases in the data, but also potentially lead to decreased power to detect interaction effects (36). In CaG whole blood samples, the interaction effect using additive genetic effect at rs1967309 was not significant, similarly to results from GTEx in whole blood samples. However, given the larger size of the dataset, we evaluated a genotypic encoding for the rs1967309 SNP in which the interaction effect is significant (p-value=0.008, Supplementary Figure 7b) in whole blood, suggesting that rs1967309 could be modulating rs158477 eQTL effect, in this tissue at least, with a genotype-specific effect. We highlight that the sample sizes of current transcriptomic resources do not allow to detect interaction effects at genome-wide significance, however the likelihood of finding interaction effects between our two SNPs on *CETP* expression in three independent datasets is unlikely to happen by chance alone, providing evidence for a functional genetic interaction.

Given the sex-specific results reported above, we stratified our interaction eQTL analyses by sex. We observed that the interaction effect on *CETP* expression in CaG whole blood samples (N_male_=359) is restricted to male individuals, and, despite low power due to smaller sample size in GEUVADIS, the interaction is also only suggestive in males (Supplementary Figure 7c,d). In GTEx, most well-powered tissues that showed a significant effect in the sex-combined analyses also harbor male-specific interactions (Supplementary Figure 9). For instance, GTEx skin male samples (N_male_=330) show the most significant male-specific interaction effects, with the directions of effects replicating the sex-combined result in GEUVADIS (an increase of *CETP* expression for each rs158477 A allele in rs1967309 AA individuals) albeit with an observable reversal of the direction in rs1967309 GG individuals (decrease of *CETP* expression with additional rs158477 A alleles) (Figure 5c, Figure 5-figure supplement 2a). However, significant effects in females are detected in tissues not previously seen as significant for the interaction in the sex-combined analysis, in the tibial artery (Figure 5d, Figure 5-figure supplement 2) and the heart atrial appendage (Supplementary Figure 9). For tissues with evidence of sex-specific effects in stratified analyses, we also tested the effect of an interaction between sex, rs158477 and rs1967309 (Methods) on *CETP* expression: the three-way interaction is only significant for tibial artery (Figure 5-figure supplement 2).

### Epistatic effects on phenotypes

The interaction effect of rs1967309 and rs158477 on *CETP* expression in several tissues, found in multiple independent RNA-seq datasets, coupled with the detection of LRLD between these SNPs in the Peruvian population suggest that selection may act jointly on these loci, specifically in Peruvians or Andeans. These populations are well known for their adaptation to life in high altitude, where the oxygen pressure is lower and where the human body is subjected to hypoxia (37–40). High altitude hypoxia impacts individuals’ health in many ways, such as increased ventilation, decreased arterial pressure, and alterations of the energy metabolism in cardiac and skeletal muscle (41, 42). To test which phenotype(s) may explain the putative coevolution signal discovered (Step 4, Figure 1), we investigated the impact of the interaction between rs1967309 and rs158477 on several physiological traits, energy metabolism and cardiovascular outcomes using the UK Biobank and GTEx cohort (Figure 6-figure supplement 1, Supplementary Table 2). The UK Biobank has electronic medical records and GTEx has cause of death and variables from medical questionnaires (34). The interaction term was found to be nominally significant (p-value<0.05) for forced vital capacity (FVC), forced expiratory volume in 1-second (FEV1) and whole-body water mass, and suggestive (p-value<0.10) for the basal metabolic rate, all driven by the effects in females (Figure 6a). For CAD, the interaction is suggestive (p-value<0.10) and, in this case, driven by males (Figure 6a).

**Figure 6.**
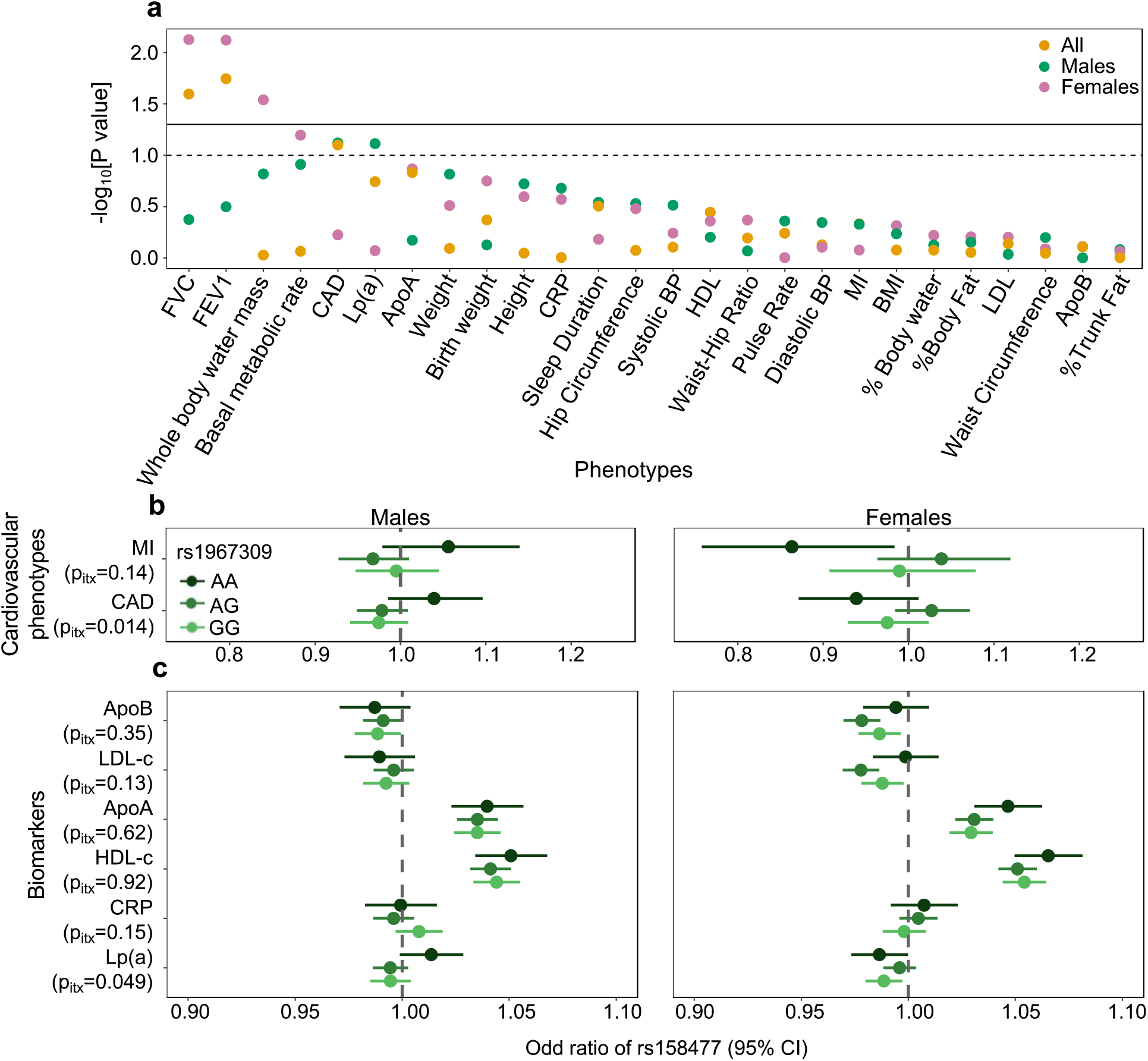
Epistatic association of rs1967309 and rs158477 on phenotypes in the UK biobank. (a) Significance of the interaction effect between rs1967309 and rs158477 on several physiological traits, energy metabolism and cardiovascular outcomes overall and stratified by sex in the UK biobank. Horizontal lines represent the p-value thresholds at 0.05 (plain) and 0.10 (dotted). Single-SNP p-values are shown in Figure 6-figure supplement 1. (b,c) Sex-stratified effects of rs158477 on (b) cardiovascular phenotypes and (c) biomarkers depending on the genotype of rs1967309 (genotypic encoding). The p-values p_itx_ reported come from a likelihood ratio test comparing models with and without the three-way interaction term between the two SNPs and sex. Sex-combined results using GTEx cardiovascular phenotype data are shown in Figure 6-figure supplement 2. See Supplementary Table 2 for the list of abbreviations.

Given this sex-specific result on CAD, the condition targeted by dalcetrapib, we tested the effect of an interaction between sex, rs158477 and rs1967309 (genotypic encoding, see Methods) on binary cardiovascular outcomes including myocardial infarction (MI) and CAD. For CAD, we see a significant three-way interaction effect, meaning that for individuals carrying the AA genotype at rs1967309, the association between rs158477 and CAD is in the opposite direction in males and females. In other words, in rs1967309-AA females, having an extra A allele at rs158477, which is associated with higher *CETP* expression (Figure 5b), has a protective effect on CAD. Conversely, in rs1967309-AA males, each A allele at rs158477 increases the probability of having an event (Figure 6a). Little effect is seen in either sex for AG or GG at rs1967309, although the heterozygotes AG behave differently in females (which further justifies the genotypic encoding of rs1967309). The beneficial effect of the interaction on CAD thus favors the rs1967309-AA + rs153477-GG males and the rs1967309-AA + rs153477-AA females, two genotype combinations which are respectively enriched in a sex-specific manner in the LIMAA cohort (Supplementary text). Again, observing such a result that concords with the direction of effects in the LRLD sex-specific finding is noteworthy. A significant interaction between the SNPs is also seen in the GTEx cohort (p-value=0.004, Figure 6-figure supplement 2, Supplementary text), using questionnaire phenotypes reporting on MI, but the small number of individuals precludes formally investigating sex effects.

Among the biomarkers studied (Supplementary Table 2), only lipoprotein(a) [Lp(a)] is suggestive in males (p-value=0.08) for an interaction between rs1967309 and rs158477, with the same direction of effect as that for CAD (Figure 6). Again, given the differences observed between the sexes, we tested the effect of an interaction between sex, rs158477 and rs1967309 (genotypic coding, Methods) on biomarkers, and only Lp(a) was nominally significant in a three-way interaction (p-value=0.049). The pattern is similar to the results for CAD, ie. a change in the effect of rs158477 depending on the genotype of rs1967309 in males, with the effect for AA females in the opposite direction compared to males (Figure 6b). These concordant results between CAD and Lp(a) support that the putative interaction effect between the loci under study on phenotypes involves sex as a modifier.

## Discussion

In this study, we used population genetics, transcriptomics and interaction analyses in biobanks to study the link between *ADCY9* and *CETP*. Our study revealed selective signatures in *ADCY9* and a significant genotypic association between *ADCY9* and *CETP* in two Peruvian cohorts, specifically between rs1967309 and rs158477, which was also seen in the Native population of the Andes. The interaction between the two SNPs was found to be nominally significant for respiratory and cardiovascular disease outcomes (Figure 6, Figure 6-figure supplement 2). Additionally, a nominally significant epistatic interaction was seen on *CETP* expression in many tissues, including the hippocampus and hypothalamus in the brain. Despite brain tissues not displaying the highest *CETP* expression levels, CETP that is synthesized and secreted in the brain could play an important role in the transport and the redistribution of lipids within the central nervous system (43, 44) and has been associated with Alzheimer’s disease risk (45, 46). These findings reinforce the fact that the SNPs are likely functionally interacting, but extrapolating on the specific phenotypes under selection from these results is not straight forward. Identifying the phenotype and environmental pressures that may have caused the selection signal is complicated by the fact that the UK Biobank participants, on which the marginally significant associations have been found, do not live in the same environment as Peruvians. In Andeans from Peru, selection in response to hypoxia at high altitude was proposed to have effects on the cardiovascular system (21). The hippocampus functions are perturbed at high altitude (eg. deterioration of memory (47, 48)), whereas the hypothalamus regulates the autonomic nervous system (ANS) and controls the heart and respiratory rates (49, 50), phenotypes which are affected by hypoxia at high altitude (51, 52). Furthermore, high altitude-induced hypoxia (53, 54) and cardiovascular system disturbances (55, 56) have been shown to be associated in several studies (57–61), thus potentially sharing common biological pathways. Therefore, our working hypothesis is that selective pressures on our genes of interest in Peru are linked to the physiological response to high-altitude, which might be the environmental driver of coevolution.

The significant interaction effects on *CETP* expression vary between sexes in amplitude and direction, with most signals driven by male samples, but significant interaction effects observed in females only, despite sample sizes being consistently lower than for males. Notably, in the tibial artery and heart atrial appendage, two tissues directly relevant to the cardiovascular system, the female-specific interaction effect on *CETP* expression is reversed between rs1967309 genotypes AA and GG, compared to the effects seen in males in skin and brain tissues. Given our *ADCY9-KD* were done in liver cell lines from male donors, future work to fully understand how rs1967309 and rs158477 interact will focus on additional experiments in cells from both male and female donors in these relevant tissues. In a previous study, we showed that inhibition of both *Adcy9* and *CETP* impacted many phenotypes linked to the ANS in male mice (6), but in the light of our results, these experiments should be repeated in female mice. The function of ANS is important in a number of pathophysiological states involving the cardiovascular system, like myocardial ischemia and cardiac arrhythmias, with significant sex differences reported (62–64).

The interaction effect between the *ADCY9* and *CETP* SNPs on both respiratory and cardiovascular phenotypes differs between the sexes, with effects on respiratory phenotypes limited to females (Figure 6a) and cardiovascular disease phenotype associations showing significant three-way sex-by-SNPs effects (Figure 6). Furthermore, the LRLD signal is present mainly in males (Figure 4), although the genotype association is also seen in female for a different genotype combination, suggesting the presence of sex-specific selection. This type of selection is very difficult to detect, especially on autosomes, with very few empirical examples found to date in the human genome despite strong theoretical support of their occurrence (65). However, sexual dimorphism in gene expression between males and females on autosomal genes has been linked to evolutionary pressures (66–68), possibly with a contribution of epistasis. As the source of selection, we favor the hypothesis of differential survival over differential ability to reproduce, because the genetic combination between *ADCY9* and *CETP* has high chances to be broken up by recombination at each generation. Even in the case where recombination is suppressed in males between these loci, they would still have equal chances to pass the favored combination to both male and female offspring, which would not explain the sex-specific LRLD signal. We see an enrichment for the rs1967309-AA + rs158477-GG in males and rs1967309-AA + rs158477-AA in females, which are the beneficial combination for CAD in the corresponding sex, possibly pointing to a sexually antagonistic selection pressure, where the fittest genotype combination depends on the sex.

Such two-gene selection signature, where only males show strong LRLD, can happen if a specific genotype combination is beneficial in creating males (through differential gamete fitness or in utero survival, for example) or if survival during adulthood is favored with a specific genotype combination compared to other genotypes. In the case of age-dependent differential survival, the genotypic association is expected to be weaker at younger ages, however the LRLD signal between rs1967309 and rs158477 in the LIMAA cohort did not depend on age neither in males nor in females (Supplementary text). Since very few individuals were younger than 20 years old, it is likely that the age range in this cohort is not appropriate to distinguish between the two possibilities. This age-dependent survival therefore remains to be tested in comparison with pediatric cohorts of Peruvians: if the LRLD signal is absent in newborns for example, it will suggest a strong selective pressure acts early in life on boys. To specifically test the in-utero hypothesis, a cohort of stillborn babies with genetic information could allow to evaluate if the genotype combination is more frequent in these. Lastly, it may be that the evolutionary pressure is linked to the sex chromosomes (69, 70), and a three-way interaction between *ADCY9*, *CETP* and Y chromosome haplotypes or mitochondrial haplogroups remains to be explored.

Even though we observed the LRLD signal between rs1967309 and rs158477 in two independent Peruvian cohorts, reducing the likelihood that our result is a false positive, one limitation is that the individuals were recruited in the same city (Lima) in both cohorts. However, we show that both populations are heterogeneous with respect to ancestry (Supplementary Figure 2), suggesting that they likely represent accurately the Peruvian population. As recent admixture and population structure can strongly influence LRLD, we performed several analyses to consider these confounders, in the full cohorts and in the sex-stratified analyses. All analyses were robust to genome-wide and local ancestry patterns, such that our results are unlikely to be explained by these effects alone (Supplementary text). Unfortunately, we could not use expression and phenotypic data from Peruvian individuals, which makes all the links between the selection pressures and the phenotype associations somewhat indirect. Future studies should focus on evaluating the phenotypic impact of the interaction specifically in Peruvians individuals, in cohorts such as the Population Architecture using Genomics and Epidemiology (PAGE) (71), in order to confirm the marginally significant associations found in European cohorts. Indeed, the Peruvian/Andean genomic background could be of importance for the interaction effect observed in this population, which reduces the power of discovery in individuals of unmatched ancestry. Furthermore, not much is known about the strength of this type of selection, and simulations would help evaluate how strong selection would need to be in a single generation to produce this level of LRLD. Another limitation is the low number of samples per tissue in GTEx and the cell composition heterogeneity per tissue and per sample (72), which can be partially captured by PEER factors and can modulate the eQTL effects. Therefore, our power to detect tissue-specific interaction effects is reduced in this dataset, making it quite remarkable that we were able to observe multiple nominally significant interaction effects between the loci.

Despite these limitations, our results support a functional role for the *ADCY9* intronic SNP rs1967309, likely involved in a molecular mechanism related to *CETP* expression, but this mechanism seems to implicate sex as a modulator in a tissue-specific way, which complicates greatly its understanding. In the dal-OUTCOMES clinical trial, the partial inhibitor of CETP, dalcetrapib, did not decrease the risk of cardiovascular outcomes in the overall population, but rs1967309 in the *ADCY9* gene was associated to the response to the drug, which benefitted AA individuals (4). Interestingly, rs1967309 AA is found in both the male and female beneficial combinations of genotypes for CAD, the same that are enriched in Peruvians, but without taking rs158477 and sex into account, this association was masked. The modulation of *CETP* expression by rs1967309 could impact CETP’s functions that are essential for successfully reducing cardiovascular events. The rs158477 locus could be a key player for these functions, and dalcetrapib may be mimicking its impact, hence explaining the pharmacogenomics association. Furthermore, in the light of our results, some of these effects could differ between men and women (73), which may need to be taken into consideration in the future precision medicine interventions potentially implemented for dalcetrapib.

In conclusion, we discovered a putative epistatic interaction between the pharmacogene *ADCY9* and the drug target gene *CETP*, that appears to be under selection in the Peruvian population. Our approach exemplifies the potential of using evolutionary analyses to help find relationships between pharmacogenes and their drug targets. We characterized the impact of the *ADCY9/CETP* interaction on a range of phenotypes and tissues. Our gene expression results in brain tissues suggest that the interaction could play a role in protection against challenges to the nervous system caused by stress such as hypoxia. The female-specific eQTL interaction results in arteries and heart tissues further suggest a link with the cardiovascular system, and the phenotype association results support further this hypothesis. In light of the associations between high altitude-induced hypoxia and cardiovascular system changes, the interaction identified in this study could be involved in both systems: for example, ADCY9 and CETP could act in pathways involved in adaptation to high altitude, which could influence cardiovascular risk via their interaction in a sex-specific manner. Finally, our findings of an evolutionary relationship between *ADCY9* and *CETP* during recent human evolution points towards a biological link between dalcetrapib’s pharmacogene *ADCY9* and its therapeutic target *CETP*.

## Material and Methods

### Key Resources Table

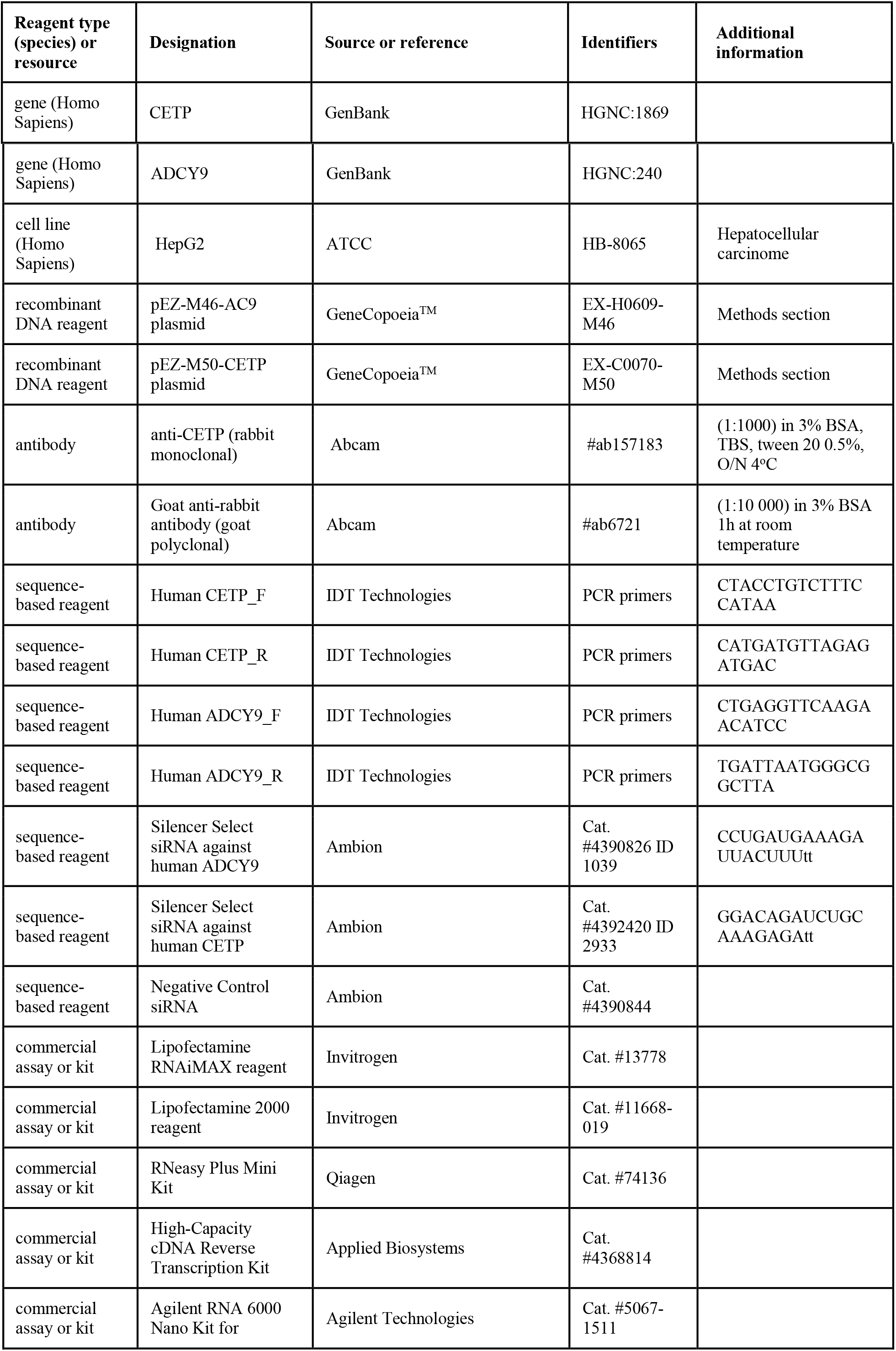
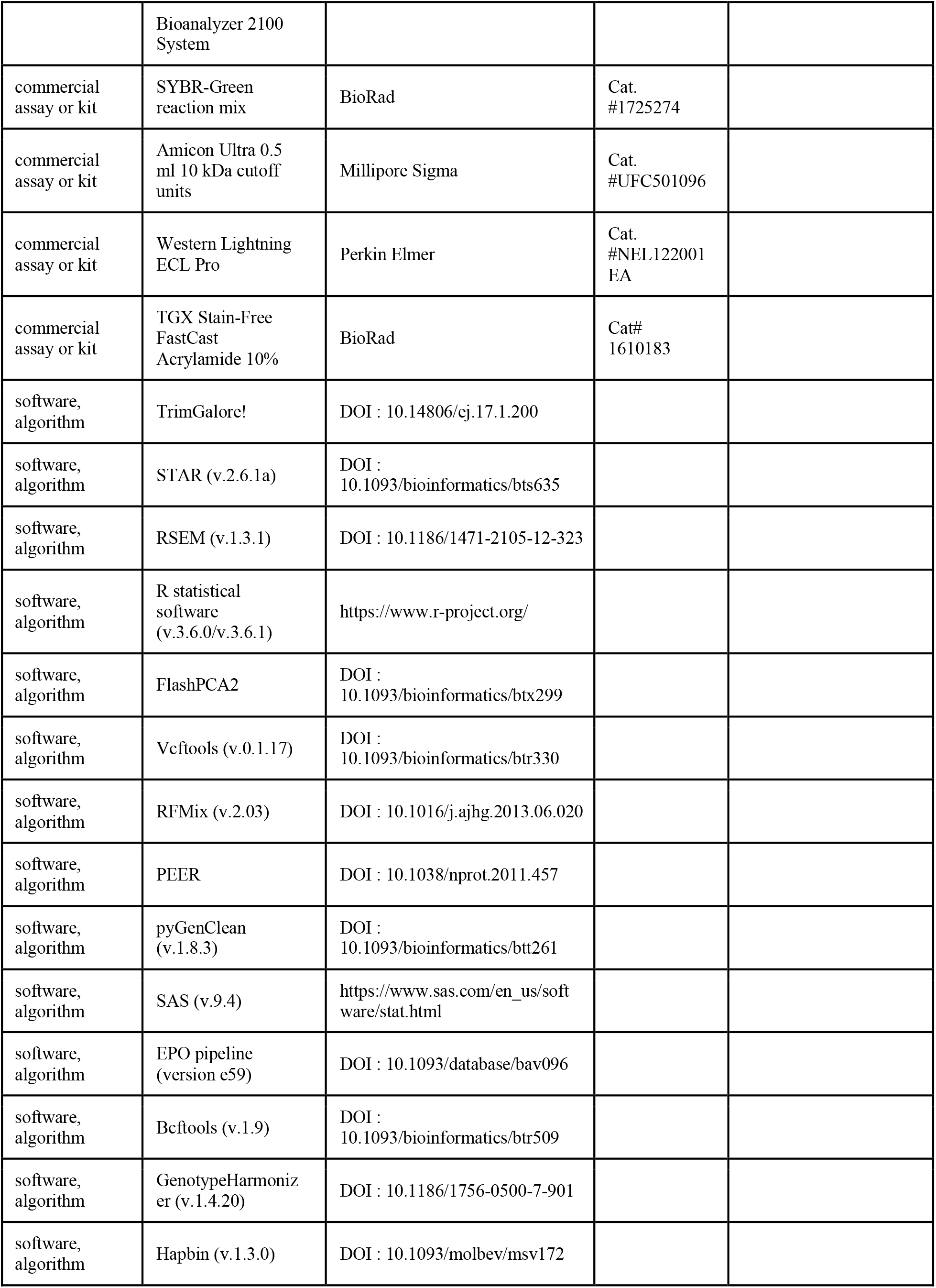
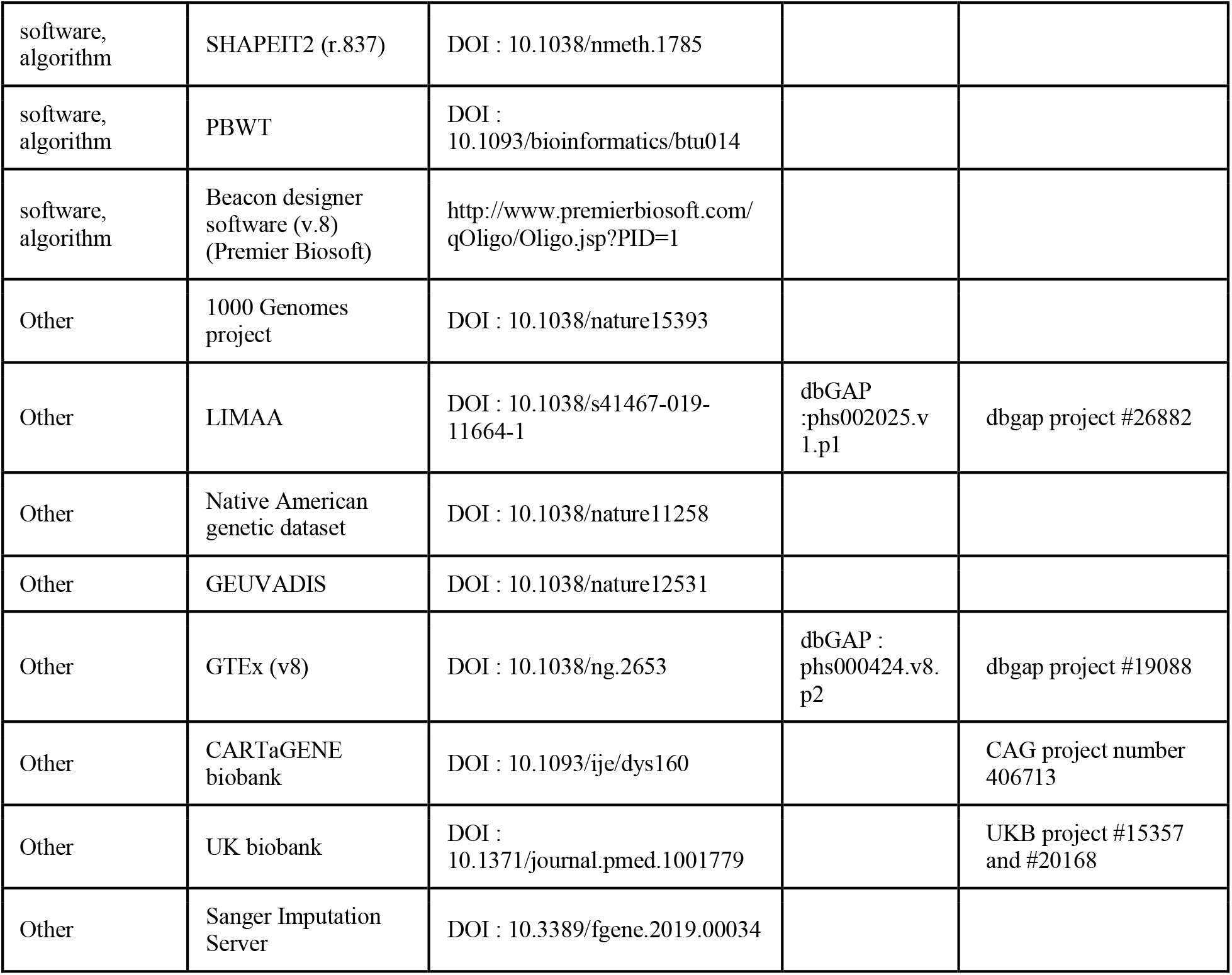

### Population Genetics Datasets

The whole-genome sequencing data from the 1000 Genomes project (1000G) Phase III dataset (ftp://ftp.1000genomes.ebi.ac.uk/vol1/ftp/release/20130502/) was filtered to exclude INDELs and CNVs so that we kept only biallelic SNPs. This database has genomic variants of 2,504 individuals across five ancestral populations: Africans (AFR, *n* = 661), Europeans (EUR, *n* = 503), East Asians (EAS, *n* = 504), South Asians (SAS, *n* = 489) and Americans (AMR, *n* = 347) (74). The replication dataset, LIMAA, has been previously published (31, 32) and was accessed through dbGaP [phs002025.v1.p1, dbgap project #26882]. This cohort was genotyped with a customized Affymetric LIMAAray containing markers optimized for Peruvian-specific rare and coding variants. We excluded related individuals as reported previously (31), resulting in a final dataset of 3,509 Peruvians. We also identified fine-scale population structure in this cohort and a more homogeneous subsample of 3,243 individuals (1,302 females and 1,941 males) in this cohort was kept for analysis (Table 1, Supplementary text). The Native American genetic dataset (NAGD) contains 2,351 individuals from Native descendants from the data from a previously published study (23). Individuals were separated by their linguistic families identified by Reich and colleagues (23). NAGD came under the Hg18 coordinates, so a lift over was performed to transfer to the Hg19 genome coordinates. Pre-processing details for these datasets are described in Supplementary text.

**Table 1.**
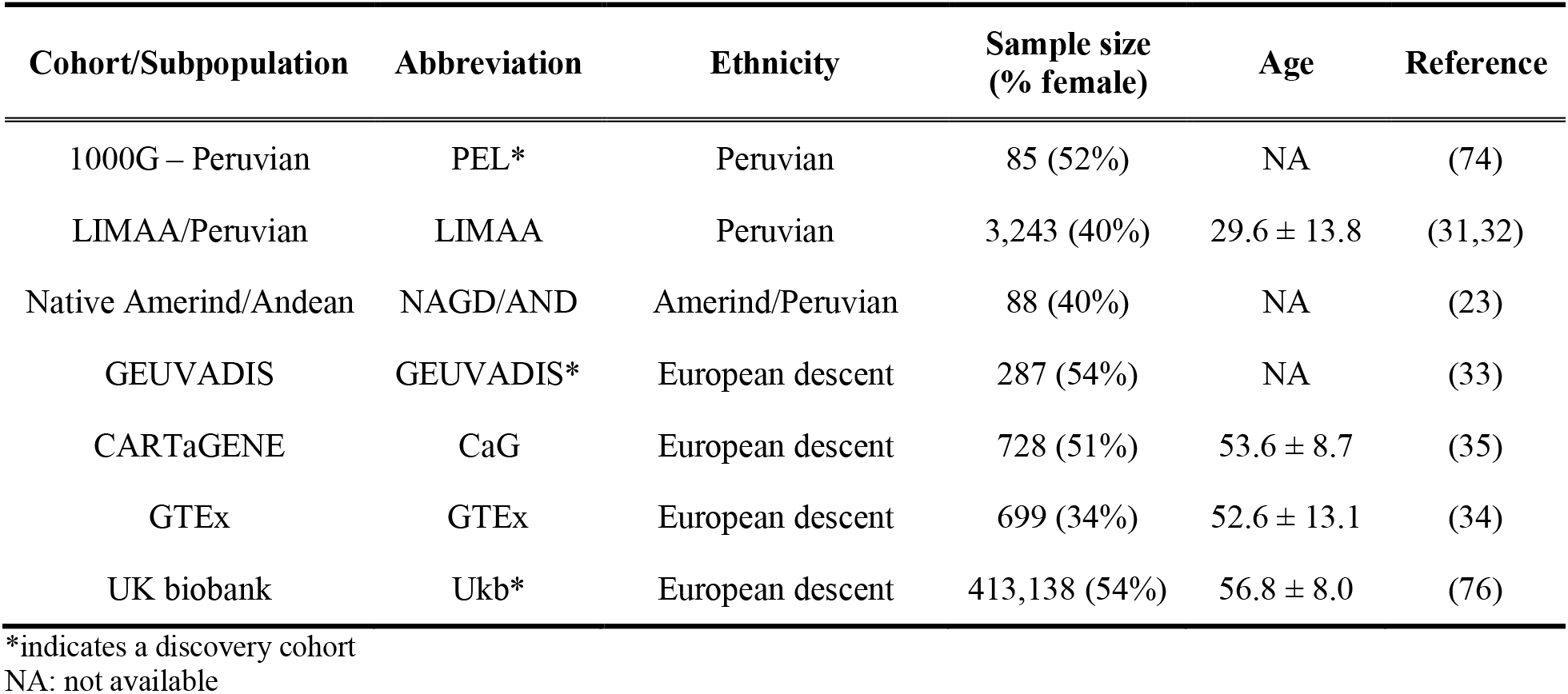
Cohort information. Sample sizes are reported after quality control steps.

### eQTL Datasets

We used several datasets (Table 1) for which we had both RNA-seq data and genotyping. First, the GEUVADIS dataset (33) for 1000G individuals was used (available at https://www.internationalgenome.org/data-portal/data-collection/geuvadis). A total of 287 non-duplicated European samples (CEU, GBR, FIN, TSI) were kept for analysis. Second, the Genotype-Tissue Expression v8 (GTEx) (34) was accessed through dbGaP (phs000424.v8.p2, dbgap project #19088) and contains gene expression across 54 tissues and 948 donors, genetic and phenotypic information. Phenotype analyses are described in Supplementary text. The cohort contains mainly of European descent (84.6%), aged between 20 and 79 years old. Analyses were done on 699 individuals, 66% of males and 34% of females (Supplementary Figure 10a). Third, we used the data from the CARTaGENE biobank (35) (CAG project number 406713) which includes 728 RNA-seq whole-blood samples with genotype data, from individuals from Quebec (Canada) aged between 36 and 72 years old (Supplementary Figure 10b). Genotyping and RNA-seq data processing pipelines for these datasets are detailed in Supplementary text. To quantify *ADCY9* gene expression, we removed the isoform transcript ENST00000574721.1 (ADCY9-205 from the Hg38) from the Gene Transfer Format (GTF) file because it is a “retained intron” and accumulates genomic noise (Supplementary text), masking true signals for *ADCY9*. To take into account hidden factors, we calculated PEER factors (75) on the normalized expressions, on all samples and stratified by sex (sPEER factors). To detect eQTL effects, we performed a two-sided linear regression on *ADCY9* and *CETP* expressions using R (v.3.6.0) (https://www.r-project.org/) with the formula *lm*(*p ∼ rs*1967309 ∗ *rs*158477 + *Covariates*) for evaluating the interaction effect, *lm*(*p ∼ rs*1967309 + *rs*158477 + *Covariates*) for the main effect of the SNPs and *lm*(*p ∼ rs*1967309 ∗ *rs*158477 ∗ *sex* + *Covariates*) for evaluating the three-way interaction effect. Under the additive model, each SNP is coded by the number of non-reference alleles (G for rs1967309 and A for rs158477), under the genotypic model, dummy coding is used with homozygous reference genotype set as reference. The covariates include the first 5 Principal Components (PCs), age (except for GEUVADIS, information not available), sex, as well as PEER factors. We tested the robustness of our results to the inclusion of different numbers of PEER factors in the models and we report them all for GEUVADIS, CARTaGENE and GTEx (Supplementary Figure 7-9). Reported values in the text are for five PEER factors in GEUVADIS, ten PEER factors in CARTaGENE, 25 sPEER for skin sun exposed in male and 10 sPEER for artery tibial in female in GTEx. Covariates specific to each cohort are reported in Supplementary text.

### UK biobank processing and selected phenotypes

The UK biobank (76) contains 487,392 genotyped individuals from the UK still enrolled as of August 20^th^ 2020, imputed using the Haplotype Reference Consortium as the main reference panel, and accessed through project #15357 and UKB project #20168. Additional genetic quality control was done using pyGenClean (v.1.8.3) (77). Variants or individuals with more than 2% missing genotypes were filtered out. Individuals with discrepancies between the self-reported and genetic sex or with aneuploidies were removed from the analysis. We considered only individuals of European ancestry based on PCs, as it is the largest population in the UK Biobank, and because ancestry can be a confounder of the genetic effect on phenotypes. We used the PCs from UK Biobank to define a region in PC space using individuals identified as “white British ancestry” as a reference population. Using the kinship estimates from the UK Biobank, we randomly removed individuals from kinship pairs where the coefficient was higher than 0.0884 (corresponding to a 3^rd^ degree relationship). The resulting post QC dataset included 413,138 individuals. For the reported phenotypes, the date of baseline visit was between 2006 and 2010. The latest available hospitalization records discharge date was June 30^th^ 2020 and the latest date in the death registries was February 14^th^ 2018. We used algorithmically-defined cardiovascular outcomes based on combinations of operation procedure codes (OPCS) and hospitalization or death record codes (ICD9/ICD10). A description of the tested continuous variables can be found in Supplementary Table 2. We used age at recruitment defined in variable #21022 and sex in variable #31. We ignored self-reported events for cardiovascular outcomes as preliminary analyses suggested they were less precise than hospitalization and death records.

In association models, each SNP analyzed is coded by the number of non-reference alleles, G for rs1967309 and A for rs158477. SNP rs1967309 was also coded as a genotypic variable, to allow for non-additive effects. For continuous traits (Supplementary Table 2) in the UK Biobank, general two-sided linear models (GLM) were performed using SAS software (v.9.4). A GLM model was first performed using the covariates age, sex and PCs 1 to 10. The externally studentized residuals were used to determine the outliers, which were removed. The normality assumption was confirmed by visual inspection of residuals for most of the outcomes, except *birthwt* and *sleep*. For biomarkers and cardiovascular endpoints, regression analyses were done in R (v.3.6.1). Linear regression analyses were conducted on standardized outcomes and logistic regression was used for cardiovascular outcomes. Marginal effects were calculated using margins package in R. In both cases, models were adjusted for age at baseline and top 10 PCs, as well as sex when not stratified. In models assessing two-way (rs1967309 by rs158477) or three-way (rs1967309 by rs158477 by sex) interactions, we used a 2 d.f. likelihood ratio test for the genotypic dummy variables’ interaction terms (genotypic model) (Supplementary text).

### RNA-sequencing of ADCY9-knocked-down HepG2 cell line

The human liver hepatocellular HepG2 cell line was obtained from ATCC, a cell line derived from the liver tissue of a 15-year-old male donor (78). Cells were cultured in EMEM Minimum essential Medium Eagle’s, supplemented with 10% fetal bovine serum (Wisent Inc). 250 000 cells in 2 ml of medium in a six-well plate were transfected using 12.5 pmol of Silencer Select siRNA against human ADCY9 (Ambion cat # 4390826 ID 1039), Silencer Select siRNA against CETP (Ambion cat 4392420 ID 2933) or Negative Control siRNA (Ambion cat #4390844) with 5 μl of Lipofectamine RNAiMAX reagent (Invitrogen cat #13778) in 500 μl Opti-MEM I reduced serum medium (Invitrogen cat # 31985) for 72h (Supplementary table 3, Supplementary text). The experiment was repeated five times at different cell culture passages. Total RNA was extracted from transfected HepG2 cells using RNeasy Plus Mini Kit (Qiagen cat #74136) in accordance with the manufacturer’s recommendation. Preparation of sequencing library and sequencing was performed at the McGill University Innovation Center. Briefly, ribosomal RNA was depleted using NEBNext rRNA depletion kit. Sequencing was performed using Illumina NovaSeq 6000 S2 paired end 100 bp sequencing lanes. Basic QC analysis of the 10 samples was performed by the Canadian Centre for Computational Genomics (C3G). To process the RNA-seq samples, we first performed read trimming and quality clipping using TrimGalore! (79) (https://github.com/FelixKrueger/TrimGalore), we aligned the trimmed reads on the Hg38 reference genome using STAR (v.2.6.1a) and we ran RSEM (v.1.3.1) on the transcriptome aligned libraries. Prior to normalization with limma and voom, we filtered out genes which had less than 6 reads in more than 5 samples. For *ADCY9* and *CETP* gene-level differential expression analyses, we compared the mean of each group of replicates with a t-test for paired samples. The transcriptome-wide differential expression analysis was done using limma, on all genes having an average of at least 10 reads across samples from a condition. Samples were paired in the experiment design. The multiple testing was taken into account by correcting the p-values with the qvalue R package (v.4.0.0) (80), to obtain transcriptome-wide FDR values.

### Overexpression of *ADCY9* and *CETP* genes in HepG2 cell line

For *ADCY9* and *CETP* overexpression experiments, 500 000 cells in 2 ml of medium in a six-well plate were transfected using 1 ug of pEZ-M46-AC9 or pEZ-M50-CETP plasmids (GeneCopoeia^TM^) with 5 ul of Lipofectamine 2000 reagent (Invitrogen cat # 11668-019) for 72h. Total RNA was extracted from transfected HepG2 cells using RNeasy Plus Mini Kit (Qiagen cat #74136) in accordance with the manufacturer’s recommendation (Supplementary table 3, Supplementary text).

### Natural selection analyses

We used the integrated Haplotype Statistic (iHS) (22) and the population branch statistic (PBS) (74) to look for selective signatures. The iHS values were computed for the each 1000G population. An absolute value of iHS above 2 is considered to be a genome wide significant signal (22). Prior to iHS computation, ancestral alleles were retrieved from 6 primates using the EPO pipeline (version e59) (81) and the filtered 1000 Genomes vcf files were converted to change the reference allele as ancestral allele using bcftools (82) with the fixref plugin. The hapbin program (v.1.3.0) (83) was then used to compute iHS using per population-specific genetic maps computed by Adam Auton on the 1000G OMNI dataset (ftp://ftp.1000genomes.ebi.ac.uk/vol1/ftp/technical/working/20130507_omni_recombination_rates). When the genetic map was not available for a subpopulation, the genetic map from the closest sub-population was selected according to their global FST value computed on the phase 3 dataset. We scanned the *ADCY9* and *CETP* genes using the population branch statistic (PBS), using 1000G sub-populations data. PBS summarizes a three-way comparison of allele frequencies between two closely related populations, and an outgroup. The grouping we focused on was PEL/MXL/CHB, with PEL being the focal population to test if allele frequencies are especially differentiated from those in the other populations. The CHB population was chosen as an outgroup to represent a Eurasian population that share common ancestors in the past with the American populations, after the out-of-Africa event. Using PJL (South Asia) and CEU (Europe) as an outgroup, or CLM as a closely related population (instead of MXL) yield highly similar results. To calculate F_ST_ for each pair of population in our tree, we used vcftools (84) by subpopulation. We calculated normalized PBS values as in (21), which adjusts values for positions with large branches in all populations, for the whole genome. We use this distribution to define an empirical threshold for significance based on the 95^th^ percentile of all PBS values genome-wide for each of the three populations.

### Long-range linkage disequilibrium

Long-range linkage disequilibrium (LRLD) was calculated using the function geno-r2 of vcftools (v.0.1.17) which uses the genotype frequencies. LRLD was evaluated in all subpopulations from 1000 Genomes Project Phase III, in LIMAA and NAGD, for all biallelic SNPs in *ADCY9* (chr16:4,012,650-4,166,186 in Hg19 genome reference) and *CETP* (chr16:56,995,835-57,017,756 in Hg19 genome reference). We analyzed loci from the phased VCF files that had a MAF of at least 5% and a missing genotype of at most 10%, in order to retain a maximum of SNPs in NAGD which has higher missing rates than the others. We extracted the 99^th^ percentile of all pairs of comparisons between *ADCY9* and *CETP* genes to use as a threshold for empirical significance and we refer to these as *ADCY9/CETP* empirical p-values. In LIMAA, we also evaluated the genotypic association using a *χ*^2^ test with four degrees of freedom (*χ*^2^) using a permutation test, as reported in (25) (Supplementary text).

Furthermore, for both cohorts, we created a distribution of LRLD values for random pairs of SNPs across the genome to obtain a genome-wide null distribution of LRLD to evaluate how unusual the genotypic association between our candidate SNPs (rs1967309-rs158477) is while taking into account the cohort-specific background genomic noise/admixture and allele frequencies. We extracted 3,513 pairs of SNPs that match rs1967309 and rs158477 in terms of MAF, physical distance (in base pairs) and genetic distance (in centiMorgans (cM), based on the PEL genetic map) between them in both cohorts (Supplementary text), and report genome-wide empirical p-values based on this distribution. For the analyses of LRLD between *ADCY9* and *CETP* stratified by sex, we considered the same set of SNP pairs that we used for the full cohorts, but separated the dataset by sex before calculating the LRLD values. To evaluate how likely the differences observed in LRLD between sex are, we also performed permutations of the sex labels across individuals to create a null distribution of sex specific effects (Supplementary text).

### Local ancestry inference

To evaluate local ancestry in the PEL subpopulation and in the LIMAA cohort, we constructed a reference panel using the phased haplotypes from 1000 Genomes (YRI, CEU, CHB) and the phased haplotypes of NAGD (Northern American, Central American and Andean) (Supplementary text). We kept overlapping positions between all datasets, and when SNPs had the exact same genetic position, we kept the SNP with the highest variance in allele frequencies across all reference populations (Supplementary text). We ran RFMix (v.2.03) (85) (with the option ‘reanalyze-reference’ and for 25 iterations) on all phased chromosomes. We estimated the whole genome average proportion of each ancestry using a weighted mean of the chromosome specific proportions given by RFMix based on the chromosome size in cM. For comparing the overall Andean enrichment inferred by RFMix between rs1967309/rs158477 genotype categories, we used a two-sided Wilcoxon-t-test. To evaluate the Andean local ancestry enrichment specifically at *ADCY9* and *CETP*, we computed the genome-wide 95^th^ percentile for proportion of Andean attribution for all intervals given by RFMix.

### Code and source data

Numerical summary data represented as a graph in main figures, as well as the code to reproduce figures and analyses, can be found here: https://github.com/HussinLab/adcy9_cetp_Gamache_2021. Raw RNA sequencing data for knocked down experiments in hepatocyte HepG2 cells are deposited the data on NCBI Gene Expression Omnibus, accession number GSE174640.

## Supporting information

Supplementary figures, tables and informations

## Acknowledgements

We thank all members of the Hussin lab for their constructive comments and feedback throughout this project, as well as the insightful input from three reviewers and reviewing and senior editors at eLife. This work was completed thanks to computational resources provided by Compute Canada clusters Graham and Beluga. This work was funded by the *Institut de Valorisation des Données* (IVADO), Health Collaboration Acceleration Fund from the Ministère de l’Économie et de l’Innovation of the Government of Quebec and the Montreal Heart Institute (MHI) Foundation. J.G.H. is a *Fonds de la Recherche en Santé* (FRQS) Junior 1 fellow. I.G. receives a PhD scholarship from the MHI Foundation and M.A.L. holds a PhD scholarship from Canadian Institutes of Health Research. M.P.D. holds the Canada Research Chair in Precision Medicine Data Analysis. J.C.T. holds the Canada Research Chair in Personalized Medicine and the Université de Montréal endowed research chair in atherosclerosis.

## Author Contributions

Conceptualization: I.G., M.P.D and J.G.H.; Data curation: I.G., M.A.L., J.C.G.; Statistical and bioinformatic analyses: I.G., M.A.L., J.C.G., H.T, S.A, A.B. and Y.F.Z.; Data acquisition: J.C.G., J.G.H, Y.L., L.L., M.M. and S.R.; Wet lab experimentation: R.S. and E.R.; Writing – Original draft: I.G. and J.G.H.; Writing – Review & editing: I.G., M.A.L., J.C.G., R.S., E.R., S.A., H.T., Y.L., S.R., J.C.T., M.P.D. and J.G.H.; Supervision: J.C.T., M.P.D. and J.G.H.; Funding acquisition: J.C.T., M.P.D., J.G.H.

---

**Figure 3-figure supplement 1.**
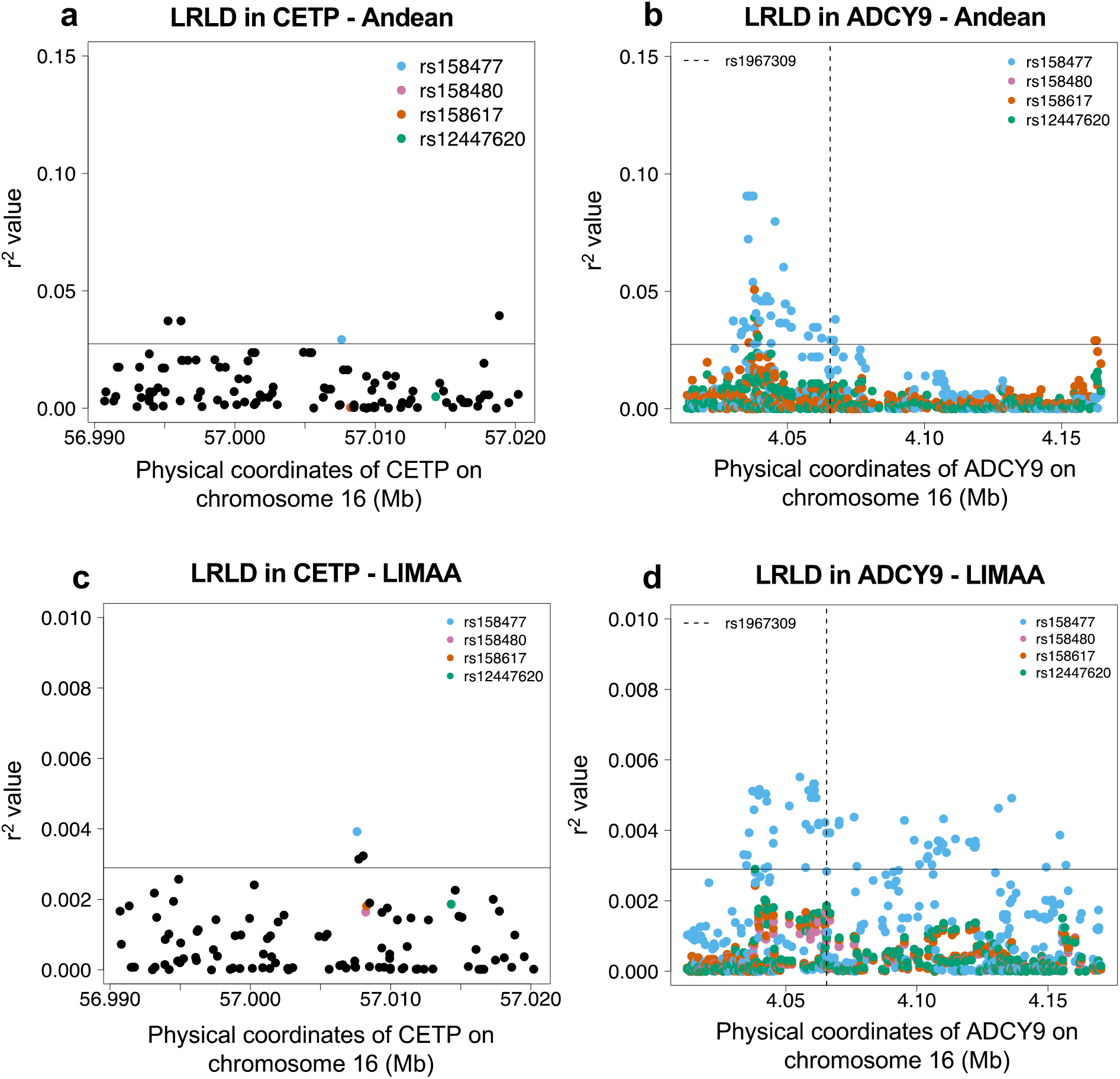
Long-range linkage disequilibrium in the Andean population from the Native Population (n=88) (a,b) and in the LIMAA cohort (n=3243) (c,d). (a,c) Genotype correlation (r^2^) between rs1967309 and all SNPs with MAF>5% in *CETP*. (b,d) Genotype correlation between the 3 loci identified in Figure 3a to be in the 99^th^ percentile and all SNPs with MAF>5% in *ADCY9*. The dotted line indicates the position of rs1967309. The horizontal lines represent the threshold for the 99^th^ percentile of all comparisons of SNPs (MAF>5%) between *ADCY9* and *CETP*.

**Figure 3-figure supplement 2.**
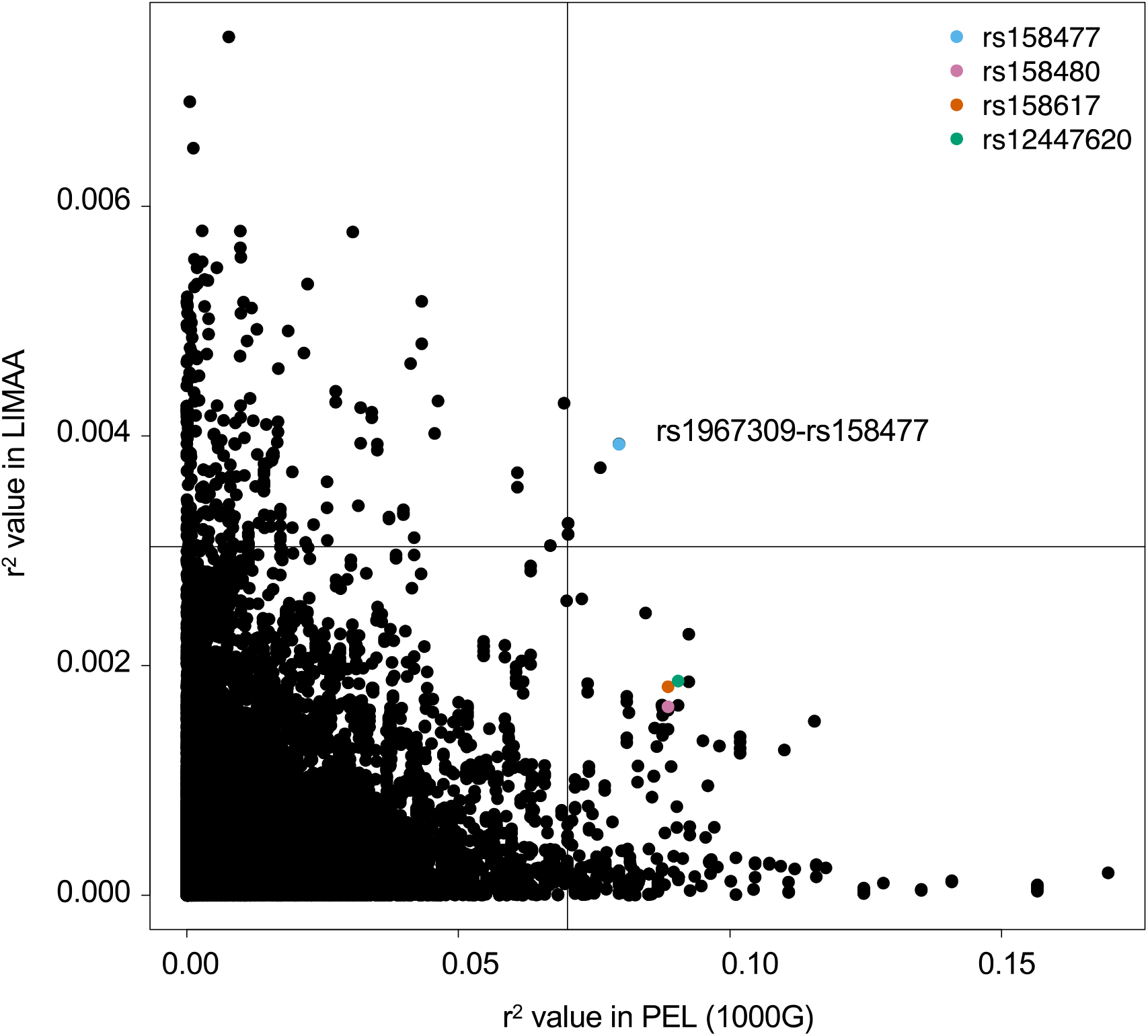
Comparison of genotype correlation between Peruvian from 1000G and from the LIMAA cohort. Comparison of genotype correlation (r^2^) between all SNPs in *ADCY9* and *CETP* with MAF>5% in the Peruvian population (PEL) in 1000G (x axis) and LIMAA cohort (y axis). Colored dots represent the value for SNPs higher than the 99^th^ percentile with rs1967309 in PEL identified in Figure 3a. Black lines represent the 99^th^ percentile in both populations.

**Figure 4-figure supplement 1.**
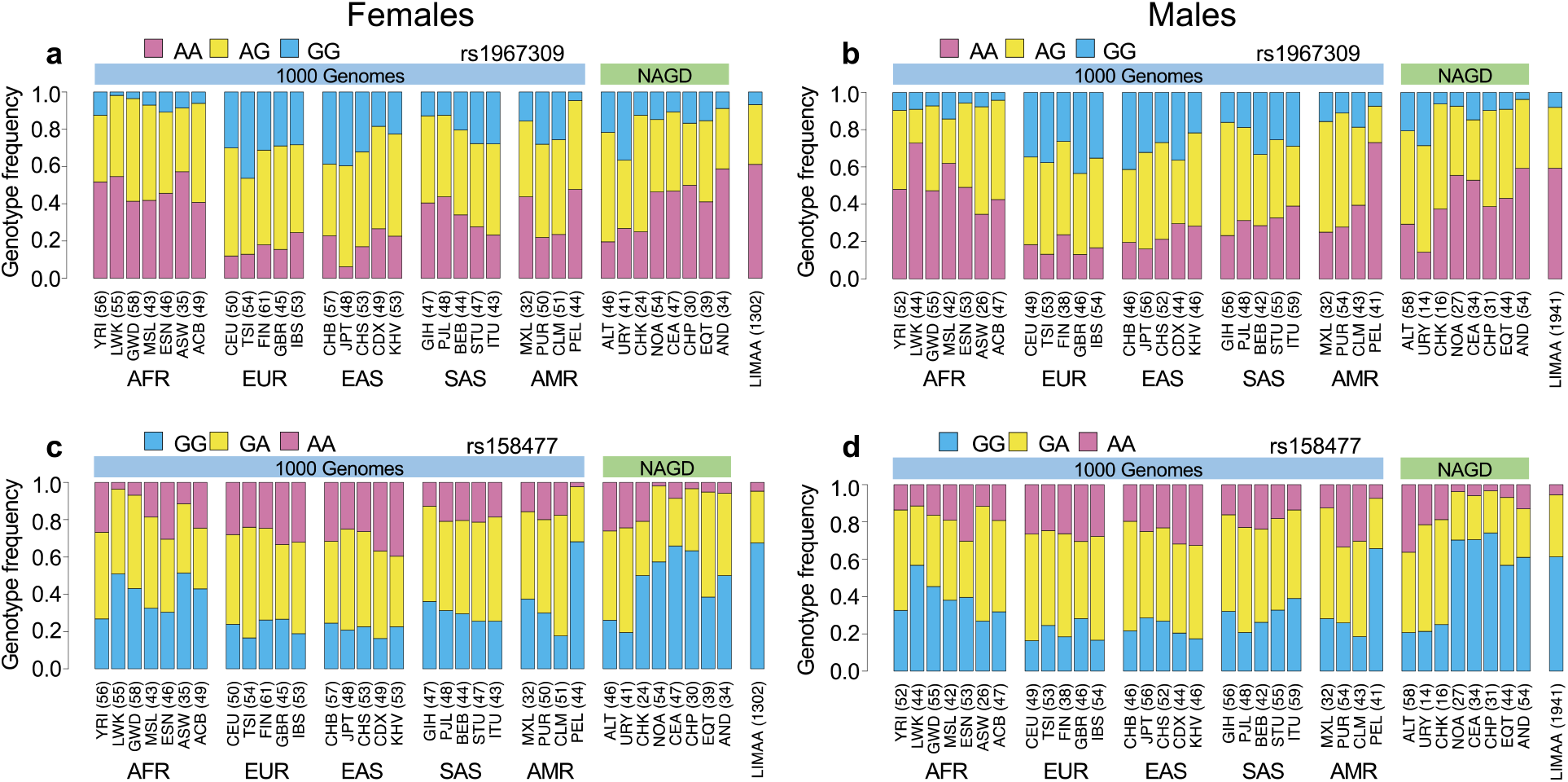
Genotype frequency distribution per sex. Genotype frequency distribution of rs1967309 in *ADCY9* (a,b) and rs158477 in *CETP* (c,d) in populations from the 1000 Genomes (1000G) Project, in Native Americans (NAGD) and LIMAA cohorts, in females (a,c) and males (b,d). Abbreviations: Altaic from Mongolia and Russia: ALT; Uralic Yukaghir from Russia: URY; Chukchi Kamchatkan from Russia: CHK; Northern American from Canada, Guatemala and Mexico: NOA; Central American from Costal Rica and Mexico: CEA; Chibchan Paezan from Argentina, Bolivia, Colombia, Costa Rica and Mexico: CHP; Equatorial Tucanoan from Argentina, Brazil, Colombia, Gualana and Paraguay: EQT; Andean from Bolivia, Chile, Colombia and Peru: AND. For 1000G populations, abbreviations can be found here https://www.internationalgenome.org/category/population/.

**Figure 4-figure supplement 2.**
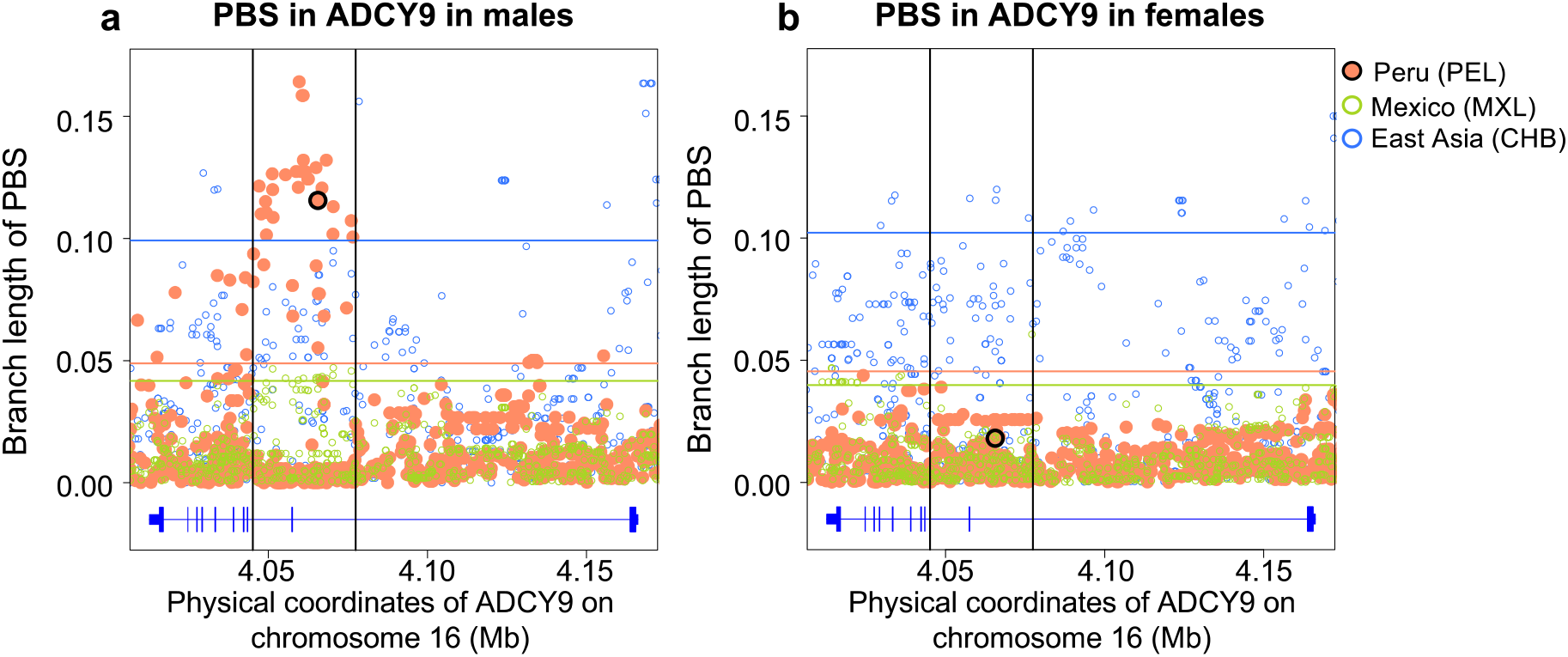
PBS values in the *ADCY9* per sex, comparing the CHB (outgroup), MXL and PEL. Horizontal lines represent the 95^th^ percentile PBS value of the chromosome 16 for each population for each sex. Vertical black lines represent the LD block around rs1967309 (shown as a black circle for PEL). Gene plots for *ADCY9* showing location of its exons are presented in blue below each plot.

**Figure 4-figure supplement 3.**
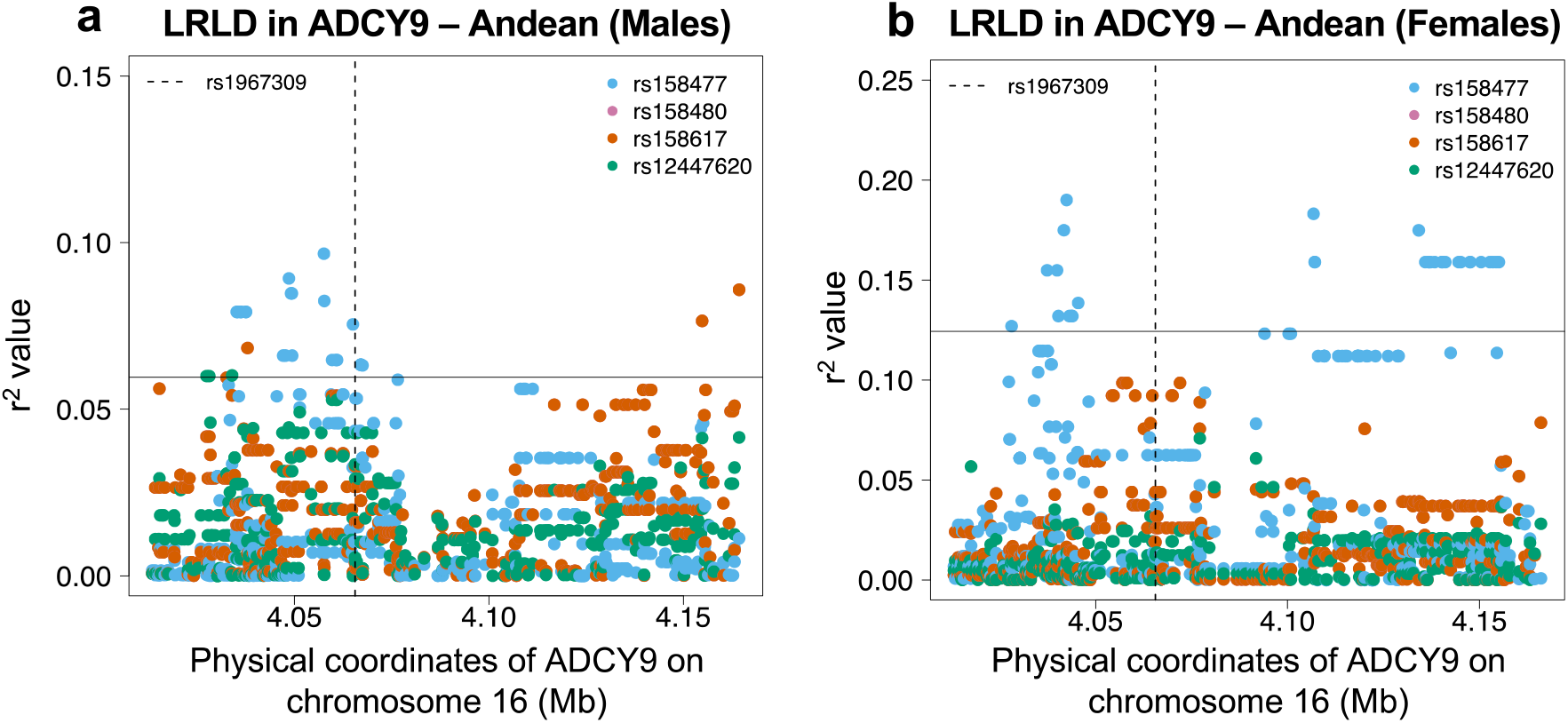
Sex-specific long-range linkage disequilibrium in the Andean population (NAGD). Genotype correlation between the loci identified in CETP in Figure 3a and all SNPs with MAF>5% in *ADCY9* for the Andean population, in males (N=54) and in females (N=34). The horizontal line shows the threshold for the 95^th^ percentile of all comparisons of SNPs (MAF>5%) between *ADCY9* and *CETP*. The vertical dotted line represents the position of rs1967309.

**Figure 5-figure supplement 1.**
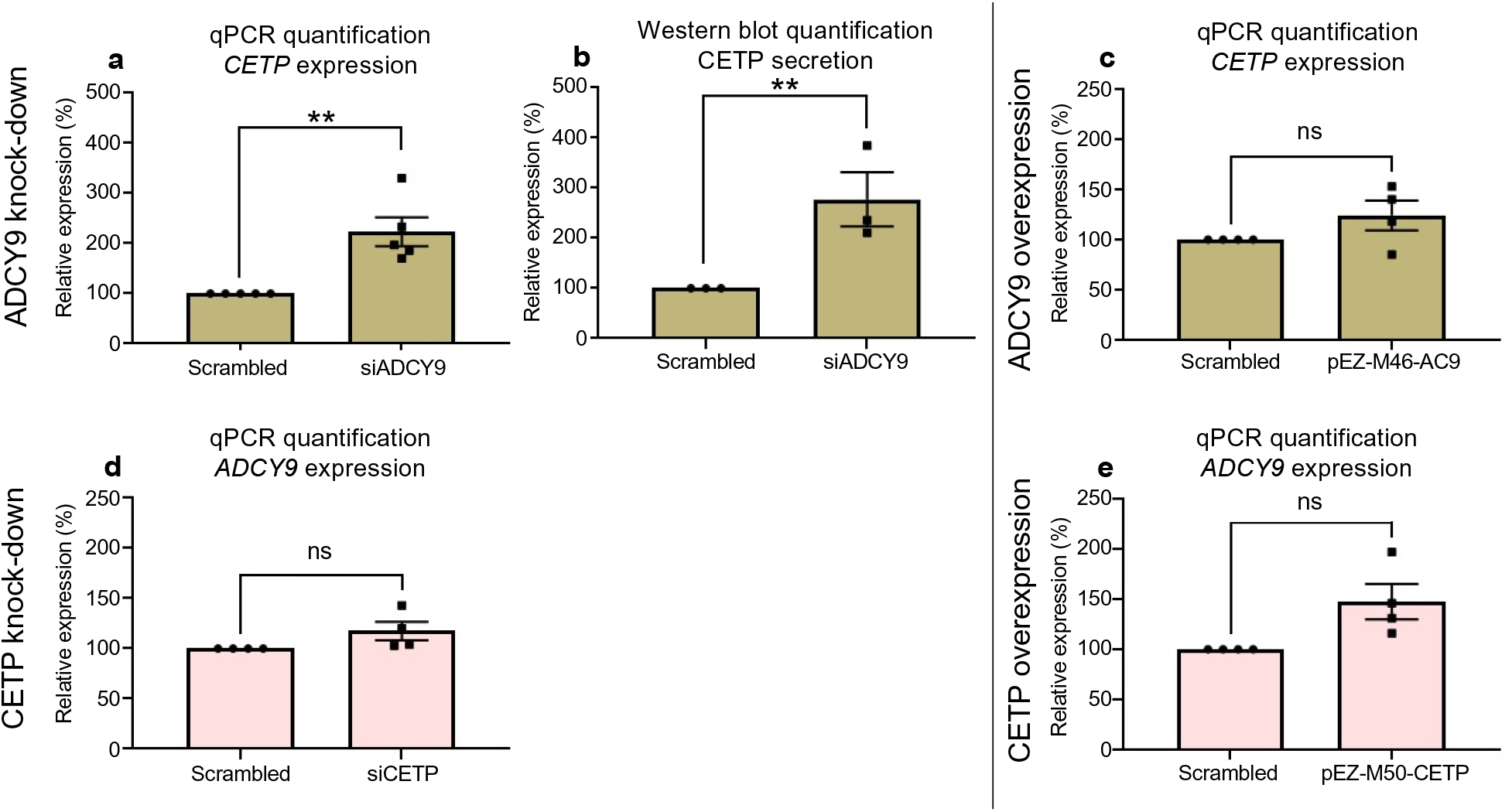
*ADCY9*/CETP interaction in HepG2 cells. (a) Relative mRNA expression of *CETP* of HepG2 cells 72h post-transfection with siRNA against human *ADCY9* (si1039). qPCR assay was normalized with PGK1 and HBS1L genes, n= 5 independent experiments, (p-value=0.0026 from t-test). (b) Quantification of CETP protein by Western blot assay, 200 ml of cell media (concentrated with Amicon ultra 0.5 ml 10 kDA units) from cells transfected with siRNA against human ADCY9 (si1039), were separated on 10% TGX-acrylamide gel and transferred to PVDF membrane. CETP protein expression was determined using a primary antibody rabbit monoclonal anti-CETP (Abcam, ab157183) 1:1000 (3% BSA, TBS, Tween 20 0.5%) O/N 4°C, followed by HRP-conjugated secondary antibody goat anti-rabbit 1:10 000 (3% BSA) 1h RT. Figure b represents densitometry analysis of n=3 experiments, p-value=0.0029 from t-test. (c,e) Relative mRNA expression of (c) *CETP* and (e) *ADCY9* genes in HepG2 cells post-transfection with pEZ-M50-CETP (overexpression of *CETP*) or pEZ-M46-ADCY9 (overexpression of *ADCY9*) plasmids. qPCR assay was normalized with PGK1 and HBS1L genes, n=4 independent experiments. (d) Relative mRNA expression of *ADCY9* of HepG2 cells 72h post-transfection with siRNA against human *CETP*. qPCR assay was normalized with PGK1 and HBS1L genes, n= 4 independent experiments.

**Figure 5-figure supplement 2.**
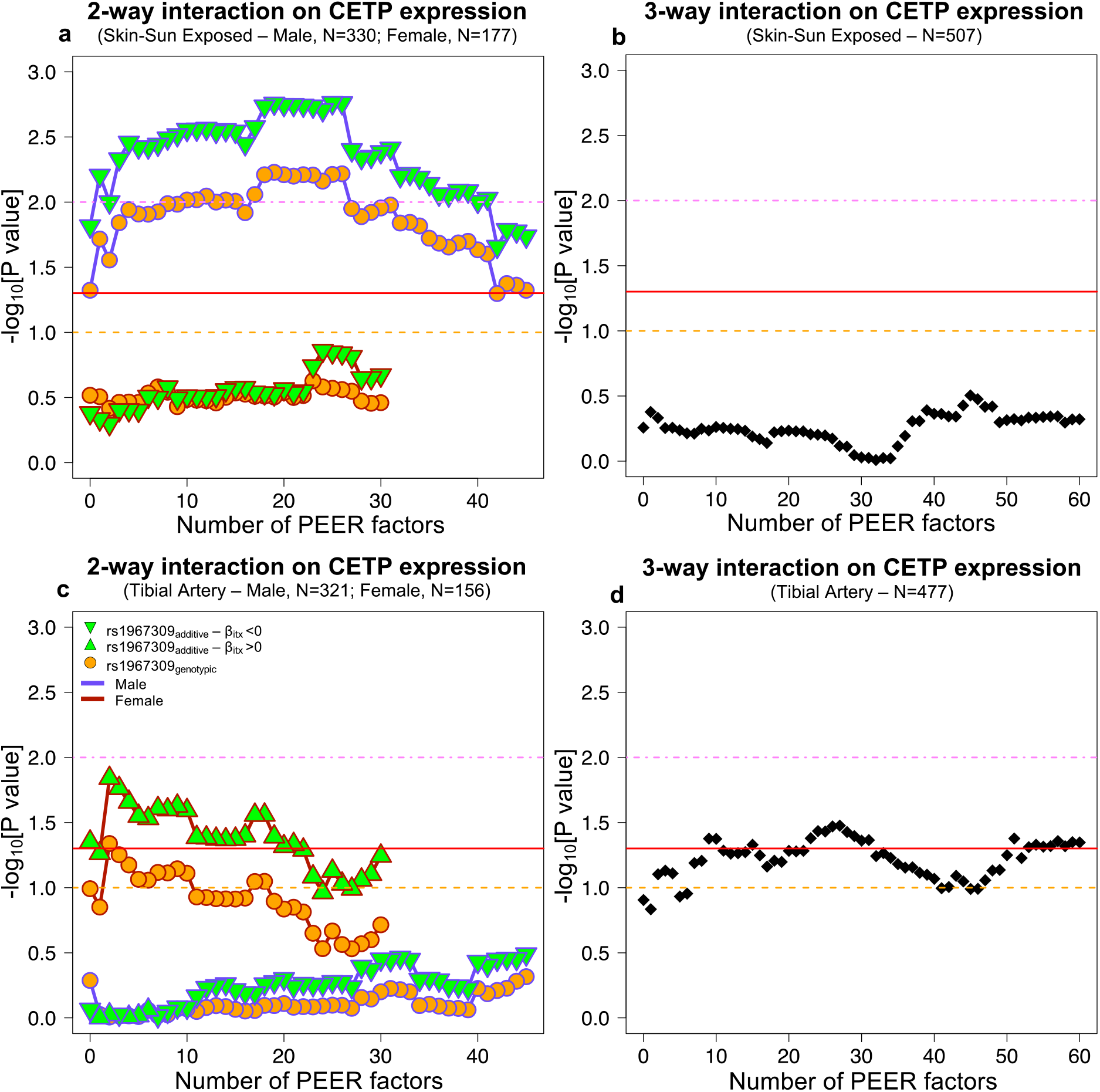
Interaction effect p-values on *CETP* expression depending by the number of PEER factors in Skin-sun exposed (a,b) and Tibial artery (c,d) in GTEx. For the two-way interaction (rs1967309*rs158477) (a,c), rs158477 is codded as additive (GG=0, GA=1, AA=2). In the additive model (green triangle), rs1967309 is codded as additive (AA=0, AG=1, GG=2). For the genotypic model (orange circle), rs1967309 was codded as a genotypic variable and p-values were obtained from a likelihood ratio test comparing models with and without the interaction term between the SNPs. The color of lines linking each value represents the sex. For the three-way interaction (rs1967309*rs158477*sex), both SNPs were codded as additive, and p-values were obtained from a linear regression model in R. P-values are presented on a -log_10_ scale. The orange, red and pink lines represent p-values of 0.1, 0.05 and 0.01 respectively.

**Figure 6-figure supplement 1.**
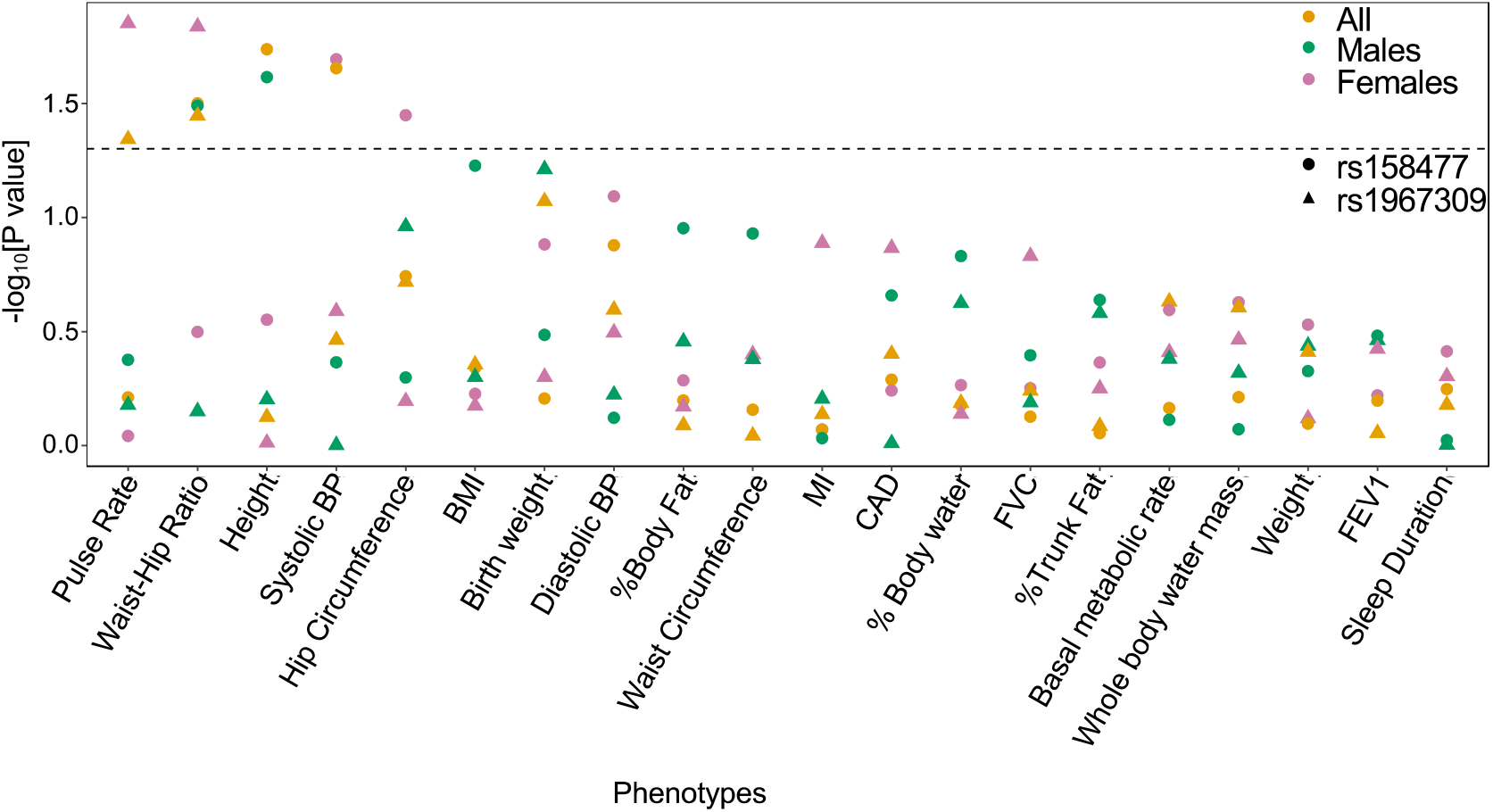
Single SNP effects of rs1967309 and rs158477 on phenotypes in the UK biobank. Significance of the marginal effect of rs1967309 and rs158477, both codded as additive, on several physiological traits, energy metabolism and cardiovascular outcomes, overall and stratified by sex in the UK biobank. The dotted line represents the p-value at 0.05. See Supplementary Table 2 for the list of abbreviations.

**Figure 6-figure supplement 2.**
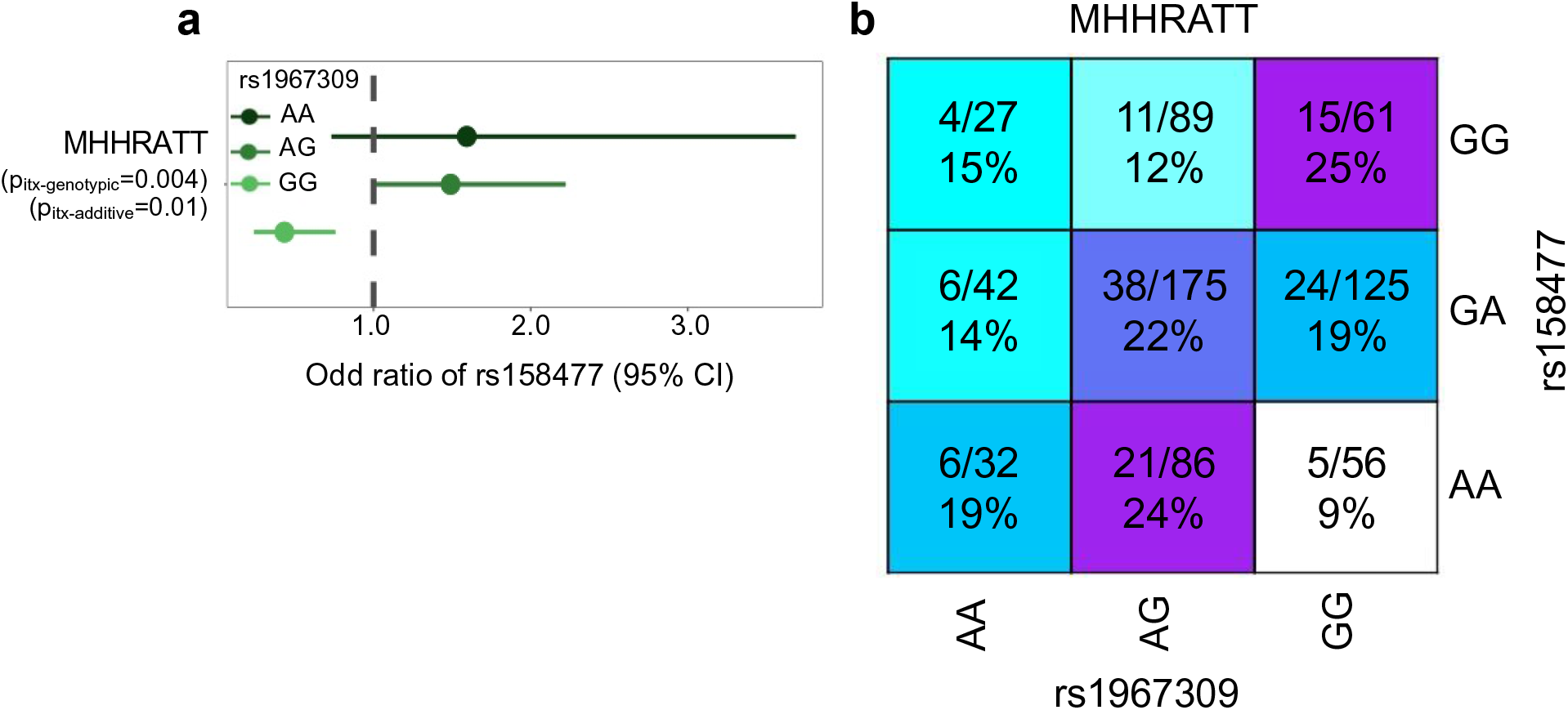
Epistatic association of rs1967309 and rs158477 on cardiovascular disease in GTEx. (a) Effect of the rs158477 SNP on the cardiovascular phenotype (n=693, cas=120, control=563) depending on the genotype of rs1967309 in GTEx. For both models, rs158477 was codded as additive (GG=0, GA=1, AA=2). For the additive model, rs1967309 was codded as additive (AA=0, AG=1, GG=2). P-value of the interaction (p_itx_) was obtained using a linear regression in R. For the genotypic model, rs1967309 was codded as a genotypic variable and p-values were obtained from a likelihood ratio test comparing models with and without the interaction term between the SNPs. (b) Proportion of cases for each genotype combinations between rs1967309 and rs158477. The numerator indicates the number of cases and the denominator the number total of individuals (cases+controls). Darker colors show higher proportions of cases.

