## Supplementary figures, tables and informations for "A sex-specific evolutionary interaction between *ADCY9* and *CETP*"

**Supplementary Material**

**Gamache et al. 2021**

### Supplementary Figures

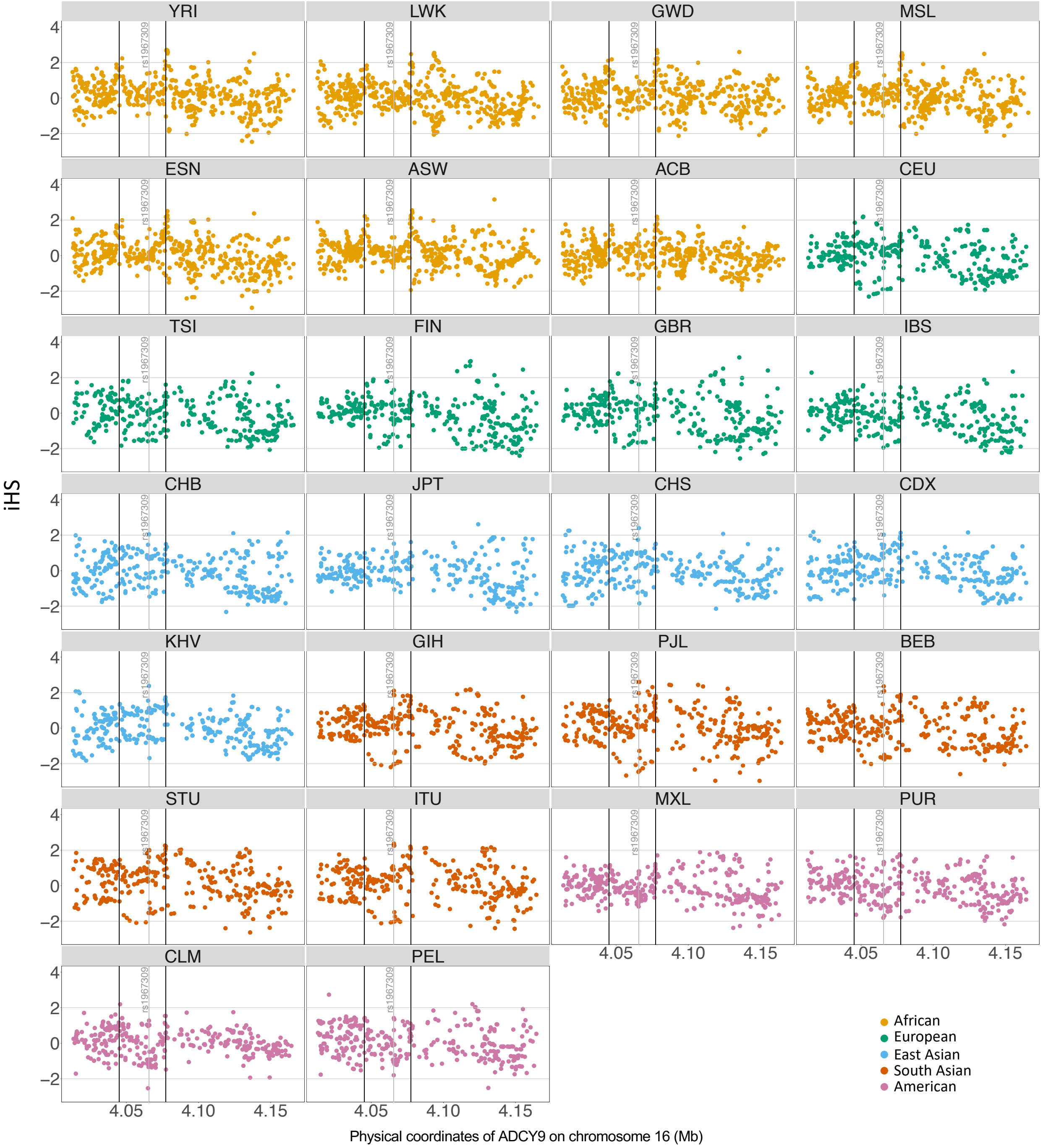

**Supplementary Figure 1. Selection signature in *ADCY9*.**
iHS values and recombination for all populations in the *ADCY9* gene. Vertical black lines represent the highest recombination rates around rs1967309 from 1000G population-specific genetic maps. Horizontal line represents the value at 2 and -2. Different colors represent one super population. In order of color: African, European, East Asia, South Asia and America. Abbreviations for the subpopulation of 1000G can be found here: https://www.internationalgenome.org/category/population/

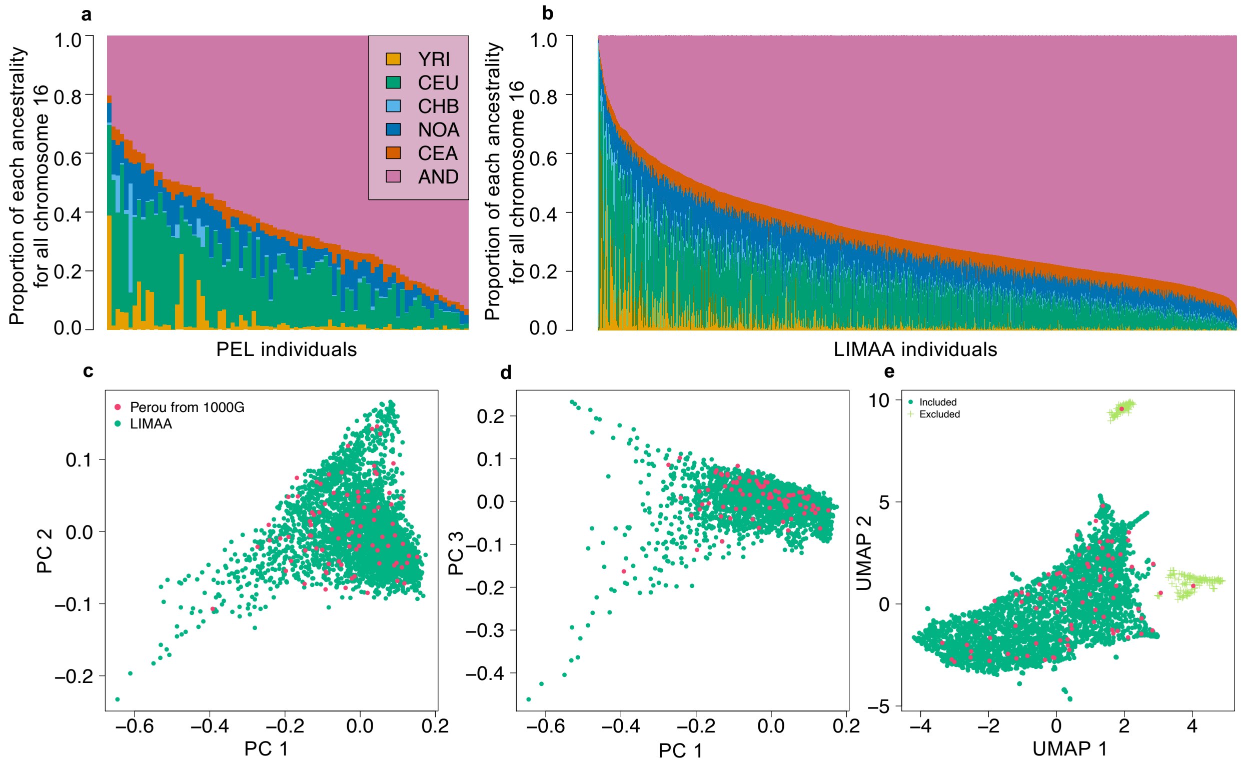

**Supplementary Figure 2. Population structure of Peruvian from LIMAA and Peruvian from 1000G.** Ancestry distribution on all chromosomes in the Peruvian from 1000G (a) and LIMAA cohort (b). Overall weighted proportion given by RFMix using reference populations from 1000G and Native American Genetic Dataset (NAGD) for the Peruvian population from 1000G (a) and from LIMAA cohort (b). 1000G populations YRI, CEU and CHB were chosen to represent African, European and Asian ancestry, respectively. (c,d) Principal Component Analysis using flashPCA on Peruvian from 1000G and LIMAA cohort. The top three PCs is shown. (e) UMAP analysis on the top 50 PCs. To limit confounders due to population structure, we excluded individuals in LIMAA coming from the two small groups identified by the UMAP (cross shaped light green symbols in (e)). Abbreviations for 1000G can be found here: https://www.internationalgenome.org/category/population/. Abbreviations for the Native American (NAGD): NOA: northern American; CEA: central American; AND: Andean.

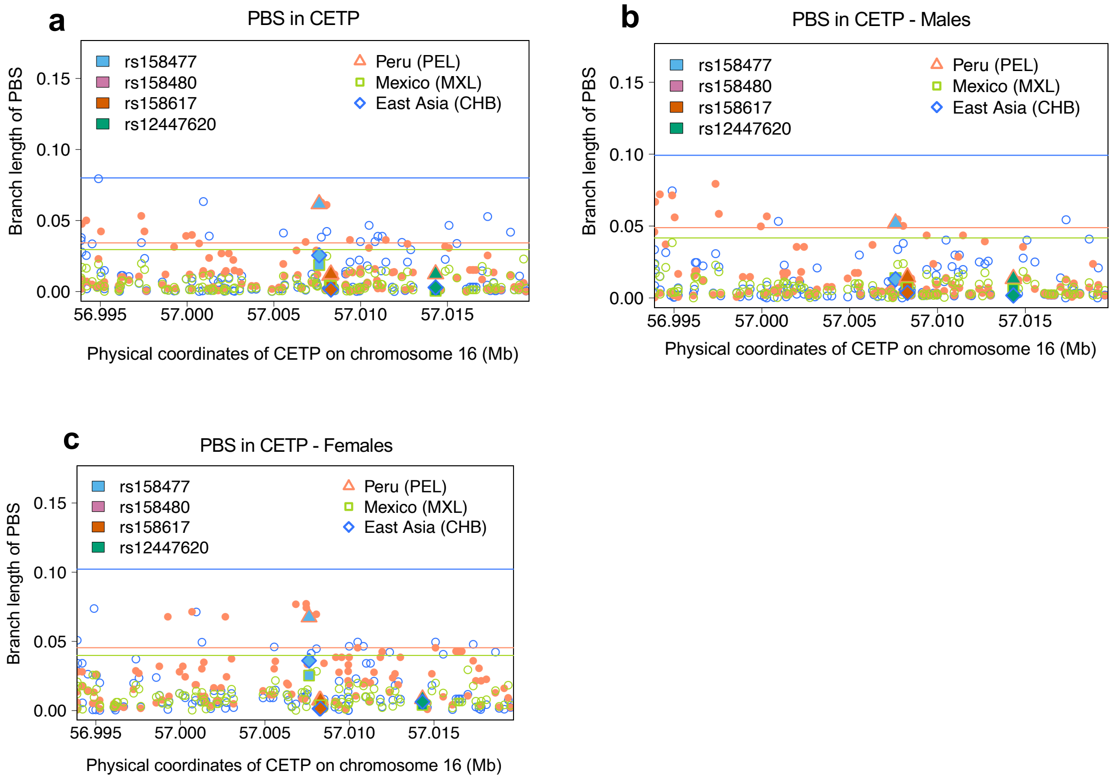

**Supplementary Figure 3. Populational differentiation of *CETP* gene using PBS statistic.** PBS values in the *CETP* gene, comparing the CHB (outgroup), MXL and PEL identified by different colors, overall (a), in males (b) and in females (c). Horizontal lines represent the 95$^{th}$ percentile PBS value genome-wide (a) or the chromosome 16 (b,c) for each population. Position with $r^2$ higher than the 99$^{th}$ percentile in the Peruvian population from the 1000G are represented by colored shape.

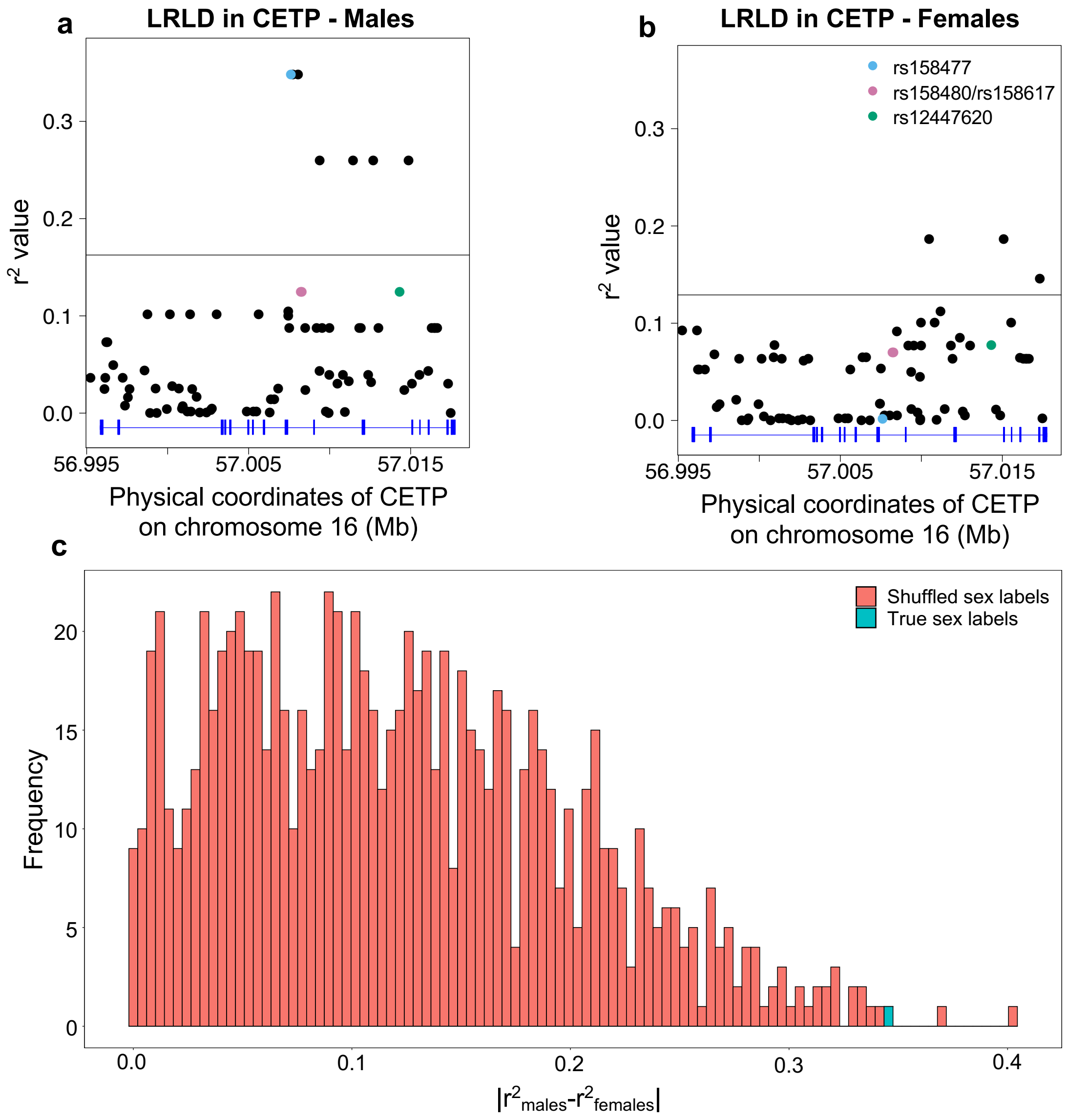

**Supplementary Figure 4. Long-range linkage disequilibrium shown in CETP for the PEL population from 1000G, stratified by sex.**

Genotype correlation ($r^2$) between the 3 loci identified in *CETP* (see Figure 2a) to be higher than the 99$^{th}$ percentile and all SNPs with MAF>5% in *ADCY9*, in males (a) and females (b). The horizontal black line is the 99$^{th}$ of all those comparisons between ADCY9 and CETP by sex. (c) Distribution of absolute difference of genotype correlation values obtained during the permutation analysis that shuffled the sex label for rs1967309 and rs158477 (red), compared to the value obtain with the real sex labels (blue).

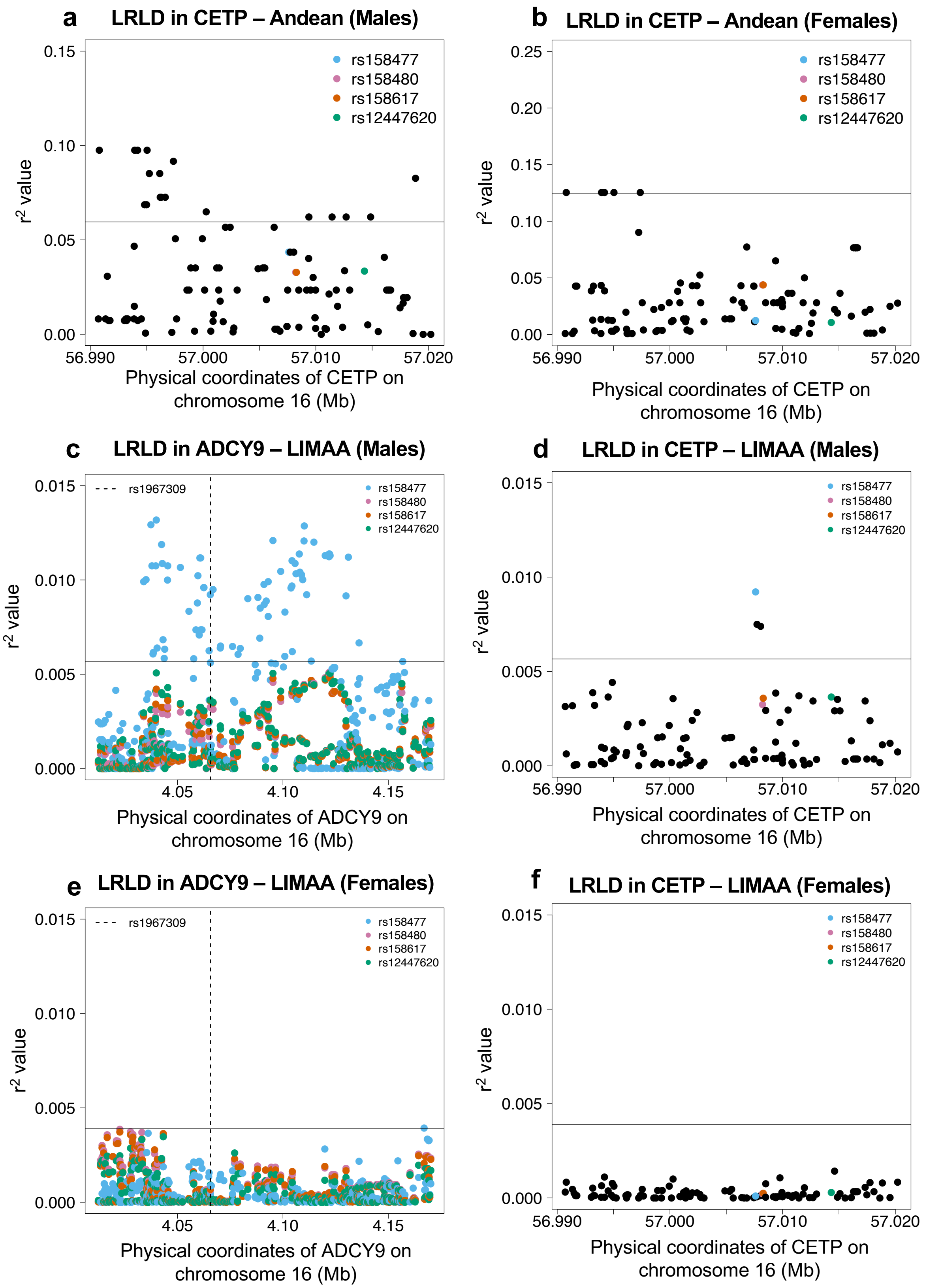
a
LRLD in CETP – Andean (Males)
rs158477
rs158480
rs158617
rs12447620
r2 value
0.15
0.10
0.05
0.00
56.990
57.000
57.010
57.020
Physical coordinates of CETP on chromosome 16 (Mb)
b
LRLD in CETP – Andean (Females)
rs158477
rs158480
rs158617
rs12447620
r2 value
0.25
0.20
0.15
0.10
0.05
0.00
56.990
57.000
57.010
57.020
Physical coordinates of CETP on chromosome 16 (Mb)
c
LRLD in ADCY9 – LIMAA (Males)
rs1967309
rs158477
rs158480
rs158617
rs12447620
r2 value
0.015
0.010
0.005
0.000
4.05
4.10
4.15
Physical coordinates of ADCY9 on chromosome 16 (Mb)
d
LRLD in CETP – LIMAA (Males)
rs158477
rs158480
rs158617
rs12447620
r2 value
0.015
0.010
0.005
0.000
56.990
57.000
57.010
57.020
Physical coordinates of CETP on chromosome 16 (Mb)
e
LRLD in ADCY9 – LIMAA (Females)
rs1967309
rs158477
rs158480
rs158617
rs12447620
r2 value
0.015
0.010
0.005
0.000
4.05
4.10
4.15
Physical coordinates of ADCY9 on chromosome 16 (Mb)
f
LRLD in CETP – LIMAA (Females)
rs158477
rs158480
rs158617
rs12447620
r2 value
0.015
0.010
0.005
0.000
56.990
57.000
57.010
57.020
Physical coordinates of CETP on chromosome 16 (Mb)

**Supplementary Figure 5. Long-range linkage disequilibrium in the Andean population from NAGD (a,b) and LIMAA cohort (c-f)**.

(a,b,d,f) Genotype correlation ($r^2$) between rs1967309 and all SNPs with MAF>5% in *CETP*, for the Andean population from NAGD (a,b) and the LIMAA cohort (d,f). (c,e) Genotype correlation between the 3 loci identified in Figure 3a to be higher than the $99^{th}$ percentile and all SNPs with MAF>5% in *ADCY9* in LIMAA. Males ($N_{Andean}$=54, $N_{LIMAA}$=1941) (a,c,d) and females ($N_{Andean}$=34, $N_{LIMAA}$=1302) (b,e,f) are shown separately. The horizontal line is the $95^{th}$ (a,b) and $99^{th}$ (c-f) percentile of all comparisons between *ADCY9* and *CETP* genes.

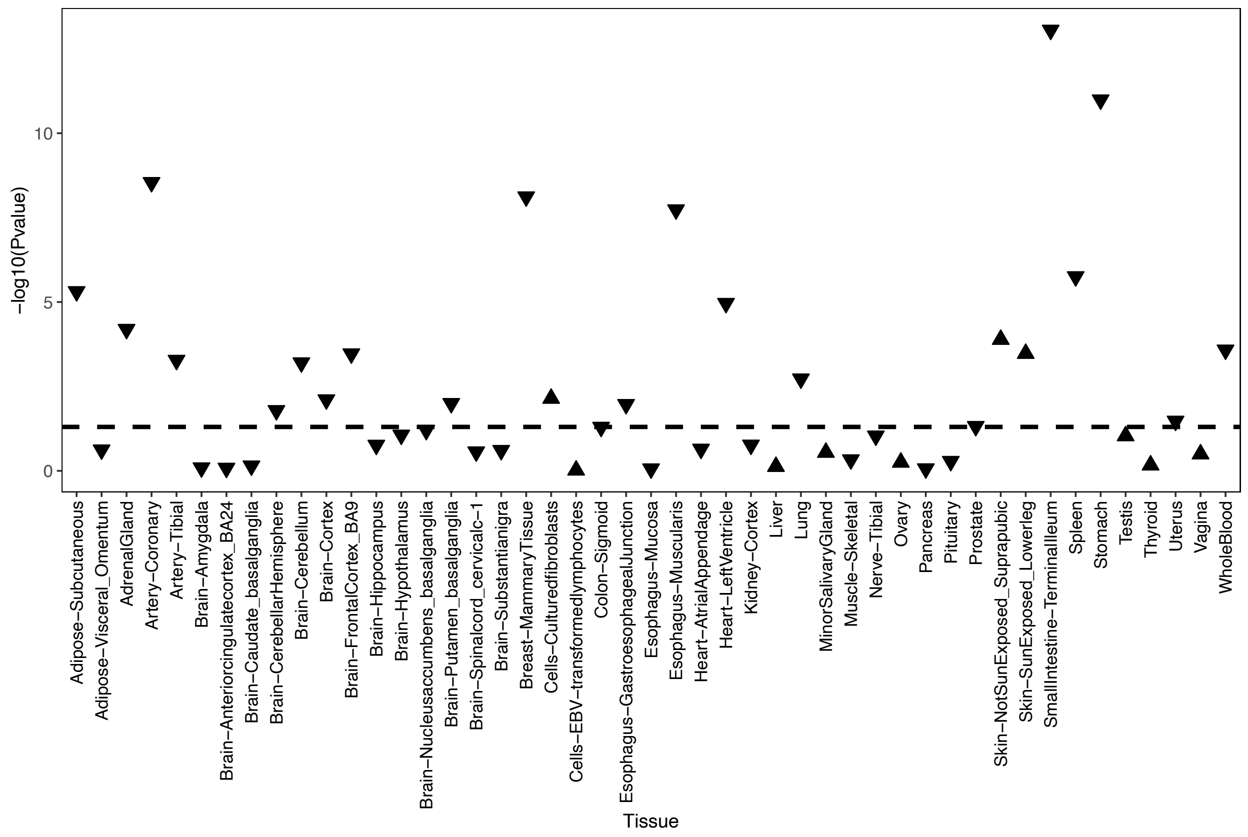

**Supplementary Figure 6. Significance of the correlation between *ADCY9* and *CETP* expression across GTEx tissues**. P-values are presented on a -$\log_{10}$ scale and are obtained from a linear regression on normalized expression with correction for age, sex, top 5 PCs, ischemic time death, sequencing platform, and sequencing center. Regular triangles mean that both gene expression levels are positively correlated, inverted triangles mean that both gene expression levels are inversely correlated. The dashed line represents the p-value at 0.05.

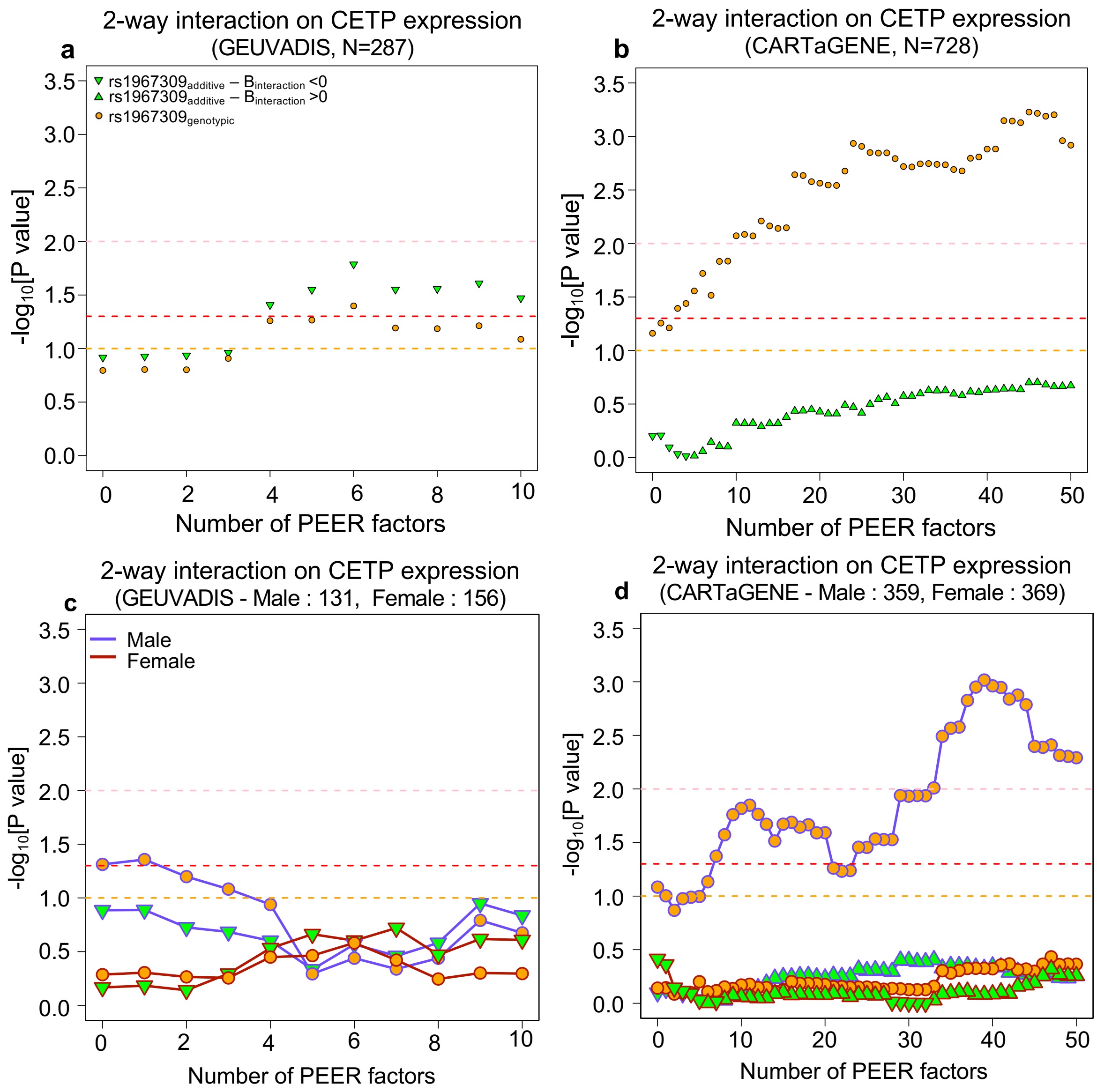

**Supplementary Figure 7. Epistatic effects between rs1967309 and rs158477 on CETP expression in GEUVADIS (LCL, N=287) and CARTaGENE (Whole blood samples, N=728).** P-values are presented on a $-\log_{10}$ scale and are reported in function of the number of PEER/sPEER factors in GEUVADIS (LCL) (a,c) and CARTaGENE (b,d) in sex-combined (a,b) and sex-stratified (c,d) analyses.

For all models, rs158477 is coded as additive (GG=0, GA=1, AA=2). In the additive model (green triangle), rs1967309 is coded as additive (AA=0, AG=1, GG=2), p-values are obtained using a linear regression in R. In the genotypic model (orange circle), rs1967309 is coded as a genotypic variable and p-values are obtained from a likelihood ratio test comparing models with and without the interaction term between the SNPs. The orange, red and pink lines represent p-values of 0.1,

0.05 and 0.01 respectively. The sample sizes reported are the number of individuals left after removing participants with missing genotypes for rs1967309 and/or rs158477. In (c,d), the color of the lines represents the sex label.

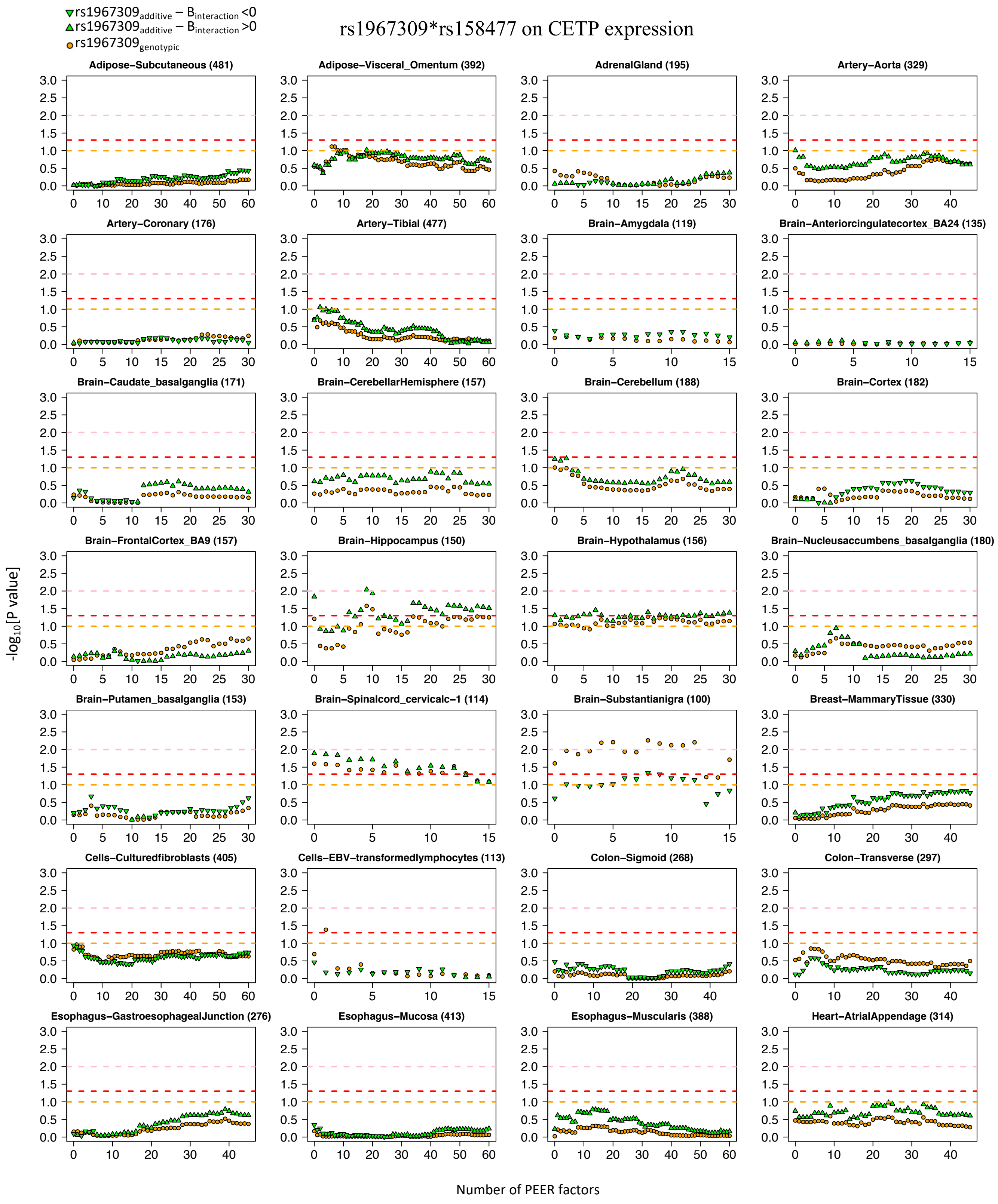
rs1967309*rs158477 on CETP expression
rs1967309$_{additive}$ − B$_{interaction}$ <0
rs1967309$_{additive}$ − B$_{interaction}$ >0
rs1967309$_{genotypic}$
Adipose−Subcutaneous (481)
Adipose−Visceral_Omentum (392)
AdrenalGland (195)
Artery−Aorta (329)
Artery−Coronary (176)
Artery−Tibial (477)
Brain−Amygdala (119)
Brain−Anteriorcingulatecortex_BA24 (135)
Brain−Caudate_basalganglia (171)
Brain−CerebellarHemisphere (157)
Brain−Cerebellum (188)
Brain−Cortex (182)
Brain−FrontalCortex_BA9 (157)
Brain−Hippocampus (150)
Brain−Hypothalamus (156)
Brain−Nucleusaccumbens_basalganglia (180)
Brain−Putamen_basalganglia (153)
Brain−Spinalcord_cervicalc-1 (114)
Brain−Substantianigra (100)
Breast−MammaryTissue (330)
Cells−Culturedfibroblasts (405)
Cells−EBV−transformedlymphocytes (113)
Colon−Sigmoid (268)
Colon−Transverse (297)
Esophagus−GastroesophagealJunction (276)
Esophagus−Mucosa (413)
Esophagus−Muscularis (388)
Heart−AtrialAppendage (314)
-log$_{10}$[P value]
Number of PEER factors

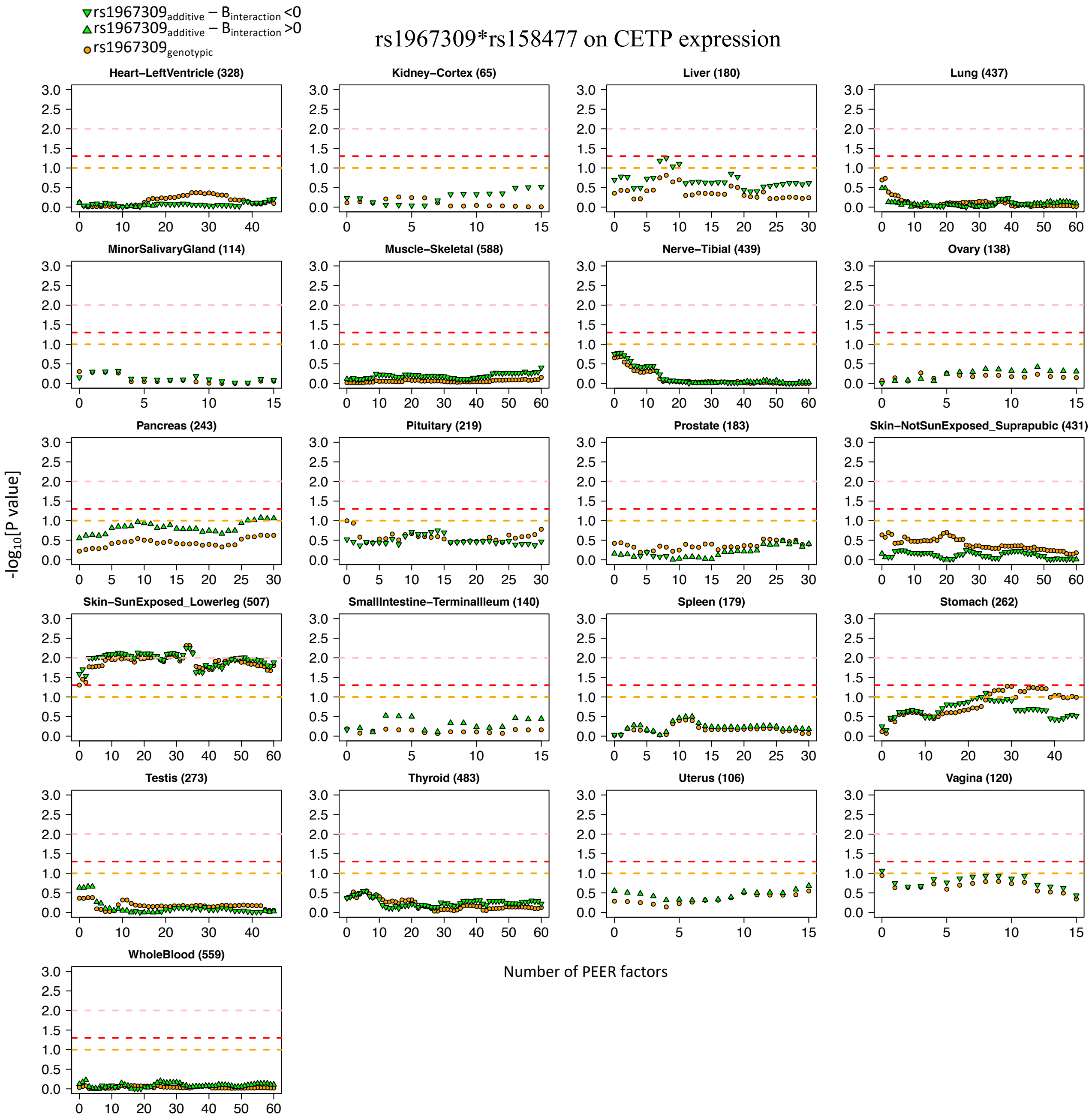

**Supplementary Figure 8. Sex-combined epistatic effect p-values for the interaction between rs1967309 and rs158477 on *CETP* expression depending on the number of PEER factors in GTEx by tissue**.

P-values are presented on a -$\log_{10}$ scale. For all models, rs158477 is coded as additive (GG=0, GA=1, AA=2). In the additive model (green triangle), rs1967309 is coded as additive (AA=0, AG=1, GG=2), p-values are obtained using a linear regression in R. In the genotypic model (orange circle), rs1967309 is coded as a genotypic variable and p-values are obtained from a likelihood ratio test comparing models with and without the interaction term between the SNPs. The orange, red and pink lines represent p-values of 0.1, 0.05 and 0.01 respectively. The tissue type and the number of samples for each, used in the analysis, are reported in the titles of the subgraphs.

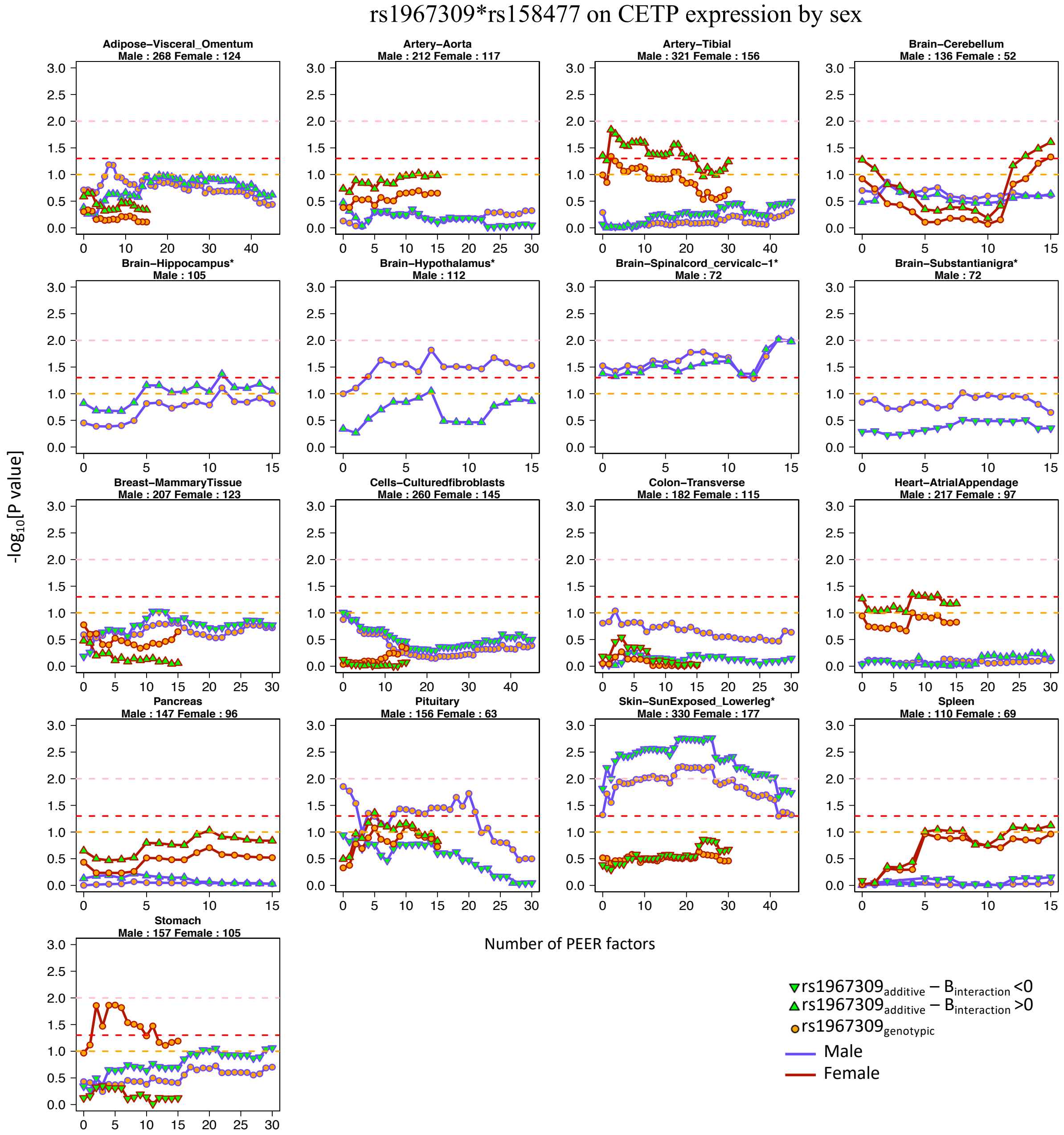

**Supplementary Figure 9. Sex-specific epistatic effects between rs1967309 and rs158477 on *CETP* expression depending on the number of sPEER factors in GTEx by tissue.**

P-values are presented on a $-\log_{10}$ scale. For all models, rs158477 is coded as additive (GG=0, GA=1, AA=2). In the additive model (green triangle), rs1967309 is coded as additive (AA=0, AG=1, GG=2), p-values are obtained using a linear regression in R. In the genotypic model (orange circle), rs1967309 is coded as a genotypic variable and p-values are obtained from a likelihood ratio test comparing models with and without the interaction term between the SNPs. The orange, red and pink lines represent p-values of 0.1, 0.05 and 0.01 respectively. The tissue type and the number of samples for each, used in the analysis, are reported in the titles of the subgraphs. The color of lines represents the sex label. Only tissues with at least one value under 0.10 are showed. Tissues with an asterisk (*) next to their title are tissues showing a the suggestive/significant effect in the sex-combined analysis.

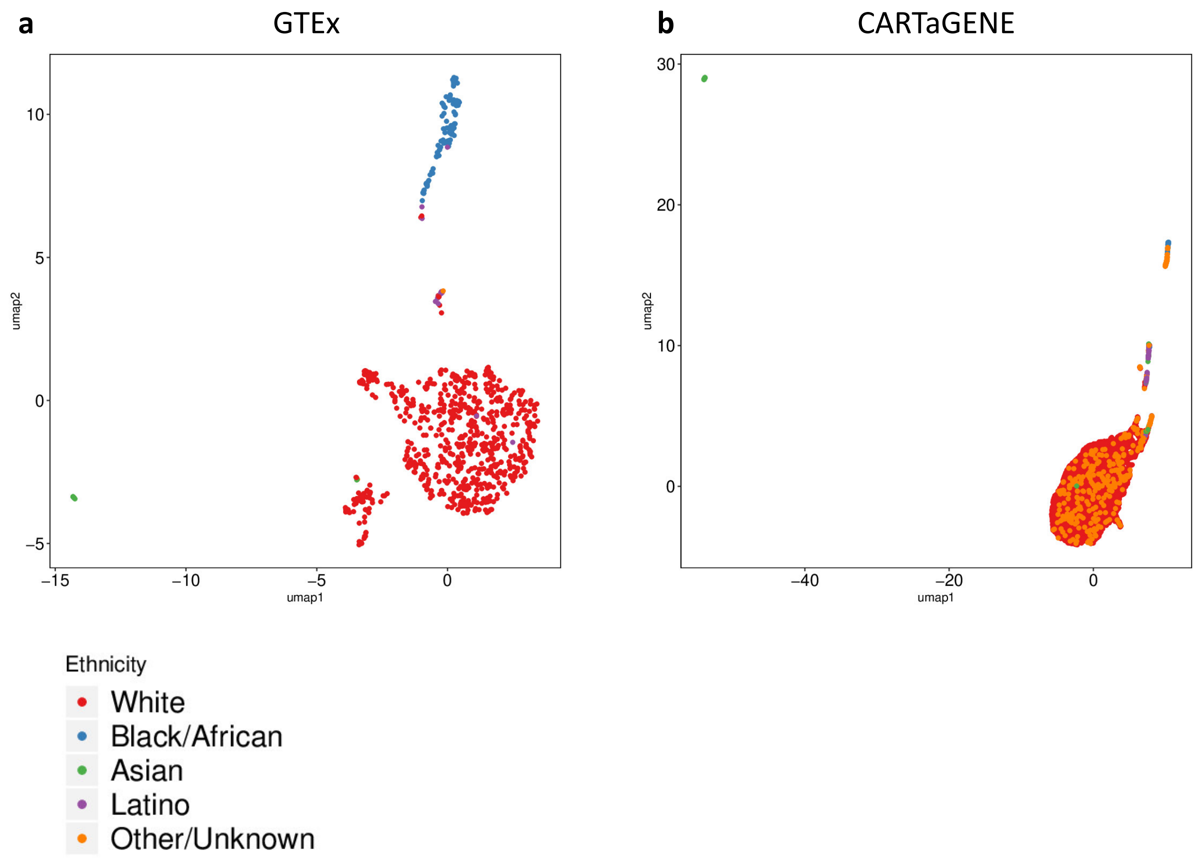

**Supplementary Figure 10. Population structure in datasets analysed.**
We estimate population structure using UMAP on the top 10 PCs generated with flashPCA2 on (a) GTEx (N=699) and (b) CARTaGENE (N=12,056) biobanks. The self-reported white non-Latino individuals were selected for further analyses.

### Supplementary Tables

Supplementary Table 1. Long-range linkage disequilibrium analysis in three datasets, and in subsets of the cohorts. Number of individuals (N) in each subset is reported. P-values correspond to the *ADCY9/CETP* empirical p-values computed as described in Section *Long-range linkage disequilibrium* in Methods. $r^2$ were obtained from the geno-r2 option of vcftools software. For 1000G populations, abbreviations can be found here https://www.internationalgenome.org/category/population/.

| Cohort | Population | Sex | Number | $r^2$ | p-value *ADCY9-CETP* |
|---|---|---|---|---|---|
| 1000G | YRI | All | 108 | 0.0236 | 0.11 |
| | CEU | All | 99 | 0.0003 | 0.86 |
| | GBR | All | 91 | 0.0117 | 0.28 |
| | CHB | All | 103 | 0.004 | 0.53 |
| | MXL | All | 64 | 0.0007 | 0.83 |
| | PEL* | All | 85 | 0.0796 | $5.42 \times 10^{-3}$ |
| | | Male | 41 | 0.3483 | $8.23 \times 10^{-5}$ |
| | | Female | 44 | 0.0016 | 0.78 |
| LIMAA | LIMAA | All | 3243 | 0.0046 | $3.24 \times 10^{-3}$ |
| | | Male | 1941 | 0.0097 | $3.71 \times 10^{-3}$ |
| | | Female | 1302 | 0.0003 | 0.52 |
| NAGD | Northern Amerind (NOA) | All | 81 | 0.0084 | 0.44 |
| | | Male | 27 | 0.0634 | 0.16 |
| | | Female | 54 | 0.0699 | 0.07 |
| | Central Amerind (CEA) | All | 81 | 0.0281 | 0.12 |
| | | Male | 34 | 0.0316 | 0.28 |
| | | Female | 47 | 0.0257 | 0.24 |
| | Andean (AND) | All | 88 | 0.0293 | 0.04 |
| | | Male | 54 | 0.0436 | 0.09 |
| | | Female | 34 | 0.0125 | 0.55 |

* Discovery cohort.

**Supplementary Table 2.** Details on metabolic and clinical variables extracted from the UK Biobank

| **Variable ID** | **UK Biobank variable location** | **Number of samples used for interaction** |
|---|---|---|
| *Category 100011 - Blood pressure - Physical measures - UK Biobank Assessment Centre* | | |
| **Pulse rate at baseline (Pulse rate)**<br>Units: bpm | Data-Field 102 (automatic entry) or Data-Field 95 (manual entry), to be derived as follows:<br>Pulse rate, automated reading (Data-Field 102) used mean of available measures for instance 0 (baseline) only. If a manual measure is available for an individual (Data Field 95 below) then do not use this automated reading (assumed to be abnormal).<br>Pulse rate (during blood-pressure measurement) (Data-Field 95), use Instance 0 (baseline). Use mean when there are multiple measures for a same individual. | All=395,319<br>Male=182,279<br>Female=213,040 |
| **Diastolic blood pressure at baseline (Diastolic BP)**<br>Units: mmHg | Data-Field 4079 (automatic entry) or Data-Field 94 (manual entry), as follow:<br>Diastolic blood pressure, automated reading: Data-Field 4079, use mean of available measures for instance 0 (baseline) only. If a manual measure is available for an individual (Data Field 94) then do not use this automated reading (assumed to be abnormal).<br>Diastolic blood pressure, manual reading: Data-Field 94, use mean of available measures for instance 0 (baseline) only. | All=395,384<br>Male=182,326<br>Female=213,058 |
| **Systolic blood pressure at baseline (Systolic BP)**<br>Units: mmHg | Data-Field 4080 (automatic entry) or Data-Field 93 (manual entry), as follow:<br>1) Systolic blood pressure, automated reading: Data-Field 4080, use mean of available measures for instance 0 (baseline) only. If a manual measure is available for an individual (Data Field 93) then do not use this automated reading (assumed to be abnormal).<br>2) Systolic blood pressure, manual reading: Data-Field 93, use mean of available measures for instance 0 (baseline) only. | All=395,353<br>Male=182,316<br>Female=213,037 |
| *Category 100010 - Body size measures - Anthropometry - Physical measures - UK Biobank Assessment Centre* | | |

| | | |
|---|---|---|
| **Waist circumference at baseline (Waist circumference)**<br>Units: cm | Data field 48, use mean of available measures for instance 0 (baseline) only. | All=395,006<br>Male=182,089<br>Female=212,917 |
| **Hip circumference at baseline (Hip circumference)**<br>Units: cm | Data field 49, use mean of available measures for instance 0 (baseline) only. | All=394,651<br>Male=181,988<br>Female=212,663 |
| **Waist-hip ratio** | Compute waist/hip | All=394,944<br>Male=182,056<br>Female=212,888 |
| **Weight**<br>Units: Kg | Data-Field 21002 (automatic entry) or Data-Field 3160 (manual entry), as follow:<br>3) Weight: Data-Field 21002, use mean of available measures for instance 0 (baseline) only.<br>Only if unavailable, then use:<br>4) Weight, manual reading: Data-Field 3160, use mean of available measures for instance 0 (baseline) only. | All=394,377<br>Male=181,732<br>Female=212,645 |
| **Height**<br>Units: cm | Data-Field 50 or 12144.<br>5) Standing height: Data Field 50, used mean of available measures for instance 0 (baseline) only.<br>Only if unavailable, then use:<br>6) Height: Data-Field 12144, used mean of available measures, as this is a singular instance field | All=394,871<br>Male=181,969<br>Female=212,902 |
| **UK Biobank BMI (BMI)**<br>Units: Kg/m2 | Data field 21001, used mean of available measures for instance 0 (baseline) only. | All=394,173<br>Male=181,705<br>Female=212,468 |
| *Category 100009 - Impedance measures - Anthropometry - Physical measures - UK Biobank Assessment Centre* | | |
| **Trunk fat percentage (% Trunk fat)**<br>Units: % | Data field 23127, use mean of available measures for instance 0 (baseline) only. | All=388,569<br>Male=178,837<br>Female=209,732 |
| **Body fat percentage (% Body fat)** | Data field 23099, use mean of available measures for instance 0 (baseline) only. | All=388,600<br>Male=178,752 |

| | | |
|---|---|---|
| Units: % | | Female=209,848 |
| **Basal metabolic rate**<br>Units: KJ | Data field 23105, use mean of available measures for instance 0 (baseline) only. | All=388,585<br>Male=178,758<br>Female=209,827 |
| **Whole body water mass**<br>Unites: Kg | Data field 23102, use mean of available measures for instance 0 (baseline) only. | All=388,719<br>Male=178,881<br>Female=209.838 |
| *Category 100020 - Spirometry - Physical measures - UK Biobank Assessment Centre* | | |
| **Forced vital capacity (FVC)**<br>Units: L | Data field 20151, use mean if more than one measure. | All=297,461<br>Male=138,909<br>Female=158,552 |
| **Forced expiratory volume in 1-second (FEV1)**<br>Units: L | Data field 20150, use mean if more than one measure. | All=297,499<br>Male=138,937<br>Female=158,562 |
| *Category 100057 - Sleep - Lifestyle and environment - Touchscreen - UK Biobank Assessment Centre* | | |
| **Sleep duration**<br>Units: hours/day | Data field 1160, use mean of available measures for instance 0 (baseline) only. | All=393,133<br>Male=181,452<br>Female=211,681 |
| *Category 100072 - Early life factors - Verbal interview - UK Biobank Assessment Centre* | | |
| **Birth weight**<br>Units: Kg | Data field 20022, use mean if more than one measure. | All=227,244<br>Male=89,715<br>Female=137,529 |
| *Category 717 - Biomarkers* | | |
| **Apolipoprotein A1 (ApoA)**<br>Units : g/L | Data field 30630, use mean of available measures for instance 0 (baseline) only.<br>Standardized using the mean : (x-mean)/sd | All=413,138<br>Male=190,454<br>Female=222,684 |
| **High Density Lipoprotein (HDL-c)**<br>Units : mmol/L | Data field 30760, use mean of available measures for instance 0 (baseline) only.<br>Standardized using the mean : (x-mean)/sd | |

| | | |
|---|---|---|
| **Lipoprotein (a) (Lp(a))**<br>Units : nmol/L | Data field 30780, use mean of available measures for instance 0 (baseline) only.<br>Standardized using the mean : (x-mean)/sd | |
| **C-Reactive Protein (CRP)**<br>Units : mmol/L | Data field 30710, use mean of available measures for instance 0 (baseline) only.<br>Ln transformation, then standardized using the mean: (x-mean)/sd | |
| **Low Density Lipoprotein (LDL-c)**<br>Units : mmol/L | Data field 30790, use mean of available measures for instance 0 (baseline) only.<br>Standardized using the mean : (x-mean)/sd | |
| **Apolipoprotein B (ApoB)**<br>Units : g/L | Data field 30640, use mean of available measures for instance 0 (baseline) only.<br>Standardized using the mean : (x-mean)/sd | |
| *Category of operation procedure codes (OPCS) and hospitalization or death record codes(ICD9/ICD10)* | | |
| **Coronary artery disease (CAD)** | Prevalent or incident | (cases/controls)<br>All=413,138 (44,713/368,425)<br>Male=190,454 (29,910/160,544)<br>Female=222,684 (14,803/207,881) |
| **Myocardial Infarction (MI)** | Prevalent or incident | (cases/controls)<br>All=413,138 (18,559/394,579)<br>Male=190,454 (13,812/176,642)<br>Female=222,684 (4,747/217,937) |

**Supplementary Table 3.** Primers sequence for real-time PCR quantification in HepG2 cells for the *KD-ADCY9* and KD-CETP experimentations

| Species | Gene | Strain | Sequence |
|---|---|---|---|
| Human | *ADCY9* | Forward | 5’ CTGAGGTTCAAGAACATCC 3’ |
| | | Reverse | 5’ TGATTAATGGGCGGCTTA 3’ |
| | *CETP* | Forward | 5’ CTACCTGTCTTTCCATAA 3’ |
| | | Reverse | 5’ CATGATGTTAGAGATGAC 3’ |
| | *HBS1L* | Forward | 5’ ACAAGAATGAGGCAACAG 3’ |
| | | Reverse | 5’ AGATACTCCAGGCACTTC 3’ |
| | *PGK1* | Forward | 5’ GTGGAGGAAGAAGGGAAG 3’ |
| | | Reverse | 5’ AAGCATCATTGACATAGACAT 3’ |

**Supplementary text**

### I. Data pre-processing

#### Pre-processing of Native American

The genetic data was obtained following correspondence with Reich et al. 2012 co-authors. The Native American Genetic Dataset (NAGD) dataset being quite sparse and samples coming from many different populations, no missing data threshold nor minor allele frequency or Hardy-Weinberg equilibrium filters were applied prior to the imputation. Harmonization to the hg19 reference genome has been done using GenotypeHarmonizer v.1.4.20 (1) and bcftools v.1.9 (2) with the fixref plugin (-m flip option). Imputation was done using the Sanger Imputation Server (3) using Haplotype Reference Consortium (r1.1) reference panel, with a pre-phasing using SHAPEIT2 r.837 (4) and imputation using PBWT (5). Post-imputation quality control was done by keeping sites with an INFO score over 0.8 and keeping genotypes having a posterior probability over 0.9. SHAPEIT2 was run to get phased genotypes (parameters: effective size of 10,000, burn of 10, prune of 10, main of 25, states of 400). The obtained VCF was used in the RFMix analysis (see below). SNPs with missing genotypes higher than 90% after imputation were removed for LRLD analysis.

#### Pre-processing of the LIMAA cohort

A pre-imputation step was conducted keeping only positions passing minor allele frequency (MAF) of 1%, 1% of missing data per site and HWE p-value > 1e-5 using PLINK v.1.9 (6). Harmonization to the hg19 reference genome has then been done using GenotypeHarmonizer and bcftools with the fixref plugin (-m flip option). Imputation was done using the Sanger Imputation

Server, using Haplotype Reference Consortium (r1.1) reference panel, with a pre-phasing using SHAPEIT2 and imputation using PBWT. Post-imputation quality control was done by keeping sites with an INFO score over 0.8 and keeping genotypes having a posterior probability over 0.9. Furthermore, positions having less than 5% missing rate after the genotyping recoding step were kept and duplicated positions were removed. SHAPEIT2 was run to get phased genotypes (parameters: effective size of 10,000, burn of 10, prune of 10, main of 25, states of 400). Another dataset was built to recover one of our SNPs of interest (rs1967309), which was excluded from our previous pipeline because of their INFO score (0.79). In this new dataset, the INFO score threshold was put to 0.7 and the post-imputation position missing data threshold was set to 35%, being less stringent, but recovering our positions. To make sure imputation quality did not impact our results because of incorrectly imputed genotypes, we redid the imputation of LIMAA with the TOPMED reference panel. The imputation $r^2$ score with TOPMED is higher than 0.9 for both, and only very limited differences in imputed genotypes are seen (only 5% and 2% of individual allele mismatches in LIMAA for rs1967309 and rs158477, respectively for the 3,243 individuals).

**Pre-processing of GTEx genetic data**

Starting from the imputed genotyping dataset, we kept bi-allelic SNPs and removed positions with more than 5% missing genotype, remaining 100,986 SNPs to calculate PCA using flashPCA2. To remove the Hispanic group, we reduced the dimensionality of the top 10 Principal Components (PCs) using the R package UMAP (7) (default parameters) to obtain a two dimensional representation of the genetic information contained within those PCs. We identified the largest homogeneous group (self-reported 'white') and excluded outlier groups (Supplementary Figure

10a), used only these individuals for the rest of the analyses. We did our all subsequent analyses with 699 individuals.

**Pre-processing of CARTaGENE**

CARTaGENE biobank (8) includes 40K individuals from Quebec (Canada) having between 36 and 72 years old. 12,056 individuals were genotyped and among these 911 had RNAseq performed on whole blood (9,10) The genotypes are coming from five different genotyping arrays on which imputation was processed independently. A pre-imputation step was conducted keeping only genotypes passing maf of 1%, 1% of missing data per site and HWE p-value > 1e-5 using PLINK. Harmonization to the Hg19 reference genome has then been done using GenotypeHarmonizer and bcftools with the fixref plugin (-m flip option). Imputation was done using the Sanger Imputation Server, using Haplotype Reference Consortium (r1.1) reference panel, with a pre-phasing using SHAPEIT2 and imputation using PBWT. Post-imputation quality control was done by keeping sites with an INFO score over 0.8 and keeping individual genotypes having a posterior probability over 0.9.

To extract only white European, we used the same filter as for GTEx, except that we removed SNPs having any missing genotypes which could create bias by different chips, then followed the recommendation from flashPCA2, remaining 8,869 SNPs to calculate PCA. We reduced the dimensionality of the top 10 PCs using the R package UMAP (default parameters) to obtain a two dimensional representation of the genetic information contained within those PCs. We identified the largest homogeneous group (Supplementary Figure 10b), which contains a majority of individuals from European descent (self-reported 'white'), and used only those individuals for the rest of the analysis. We kept 11,362 individuals at the end and among these, 911 individuals for which we had RNAseq. For these individuals, we merged samples from different batches, we removed samples who had less than 10 millions of reads, remaining 790 individuals with expression. After filtering out individuals missing the genotype of either rs1967309 or rs158477 SNPs, we did our interaction analysis on 728 individuals.

### II. <u>Population genetics</u>

#### iHS analyses

We computed the integrated haplotype score (iHS) (11) for each subpopulation in the 1000 Genomes project (Methods), a statistics that allows us to detect evidence for recent strong positive selection on derived alleles. The SNP rs1967309 is located in a region of high linkage disequilibrium (LD), delimited by recombination hotspots present in all populations. Several SNPs in this LD block exhibit absolute iHS values higher than 2 in non-African populations (Figure 2b, Supplementary Figure 1), specifically in CEU and GBR (highest signal is a 15 Kb away from rs1967309), CHB, CHS, CDX, KHV, and in all SAS sub-populations, all of which showing signals in several SNPs in less than 200 base pairs from rs1967309. Of note, however, rs1967309 itself does not show value over 2 in any population. In African populations, no signal is seen in this LD block (Supplementary Figure 1). Other SNPs in *ADCY9* are found to have absolute iHS values higher than 2, especially in the long intron 1 and around the last exon, but characterizing these signals is beyond the scope of this study.

**Sex-specific differentiation at rs1967309 in *ADCY9***

We first used $F_{ST}$ to evaluate differences in genotype frequencies between males and females. In the PEL from 1000G, we saw suggestive differences between males and females around rs1967309, but did not replicate in the LIMAA cohort, which suggests it was due to small sample size (12). Another approach we took was to investigate the impact of sex on our PBS results, by splitting the sample between males and females, and recomputing all PBS values using PEL, MXL and CHB for SNPs on chromosome 16 in each subsample. We report result on chromosome 16 that account for chromosome specific population history, as in our analyses of the full cohort, tests on chromosome 16 were more conservative than on the whole genome (ie. p-values were slightly larger with chromosome 16 alone). Although over the full chromosome, the distribution was not statistically different between males and females ($PBS_{95th\text{-}PEL,male}$ = 0.043; $PBS_{95th\text{-}PEL,female}$ = 0.040) as expected, curiously the PEL branch length for all SNPs around rs1967309 increases for males compared to the full-sample results : at rs1967309, the PBS value became 0.096 in males (chromosome 16 empirical p-value = 0.004). On the other hand, for females the value dropped to 0.017 (chromosome 16 empirical p-value = 0.20). No such male-female difference is seen in *CETP*, with the PEL PBS value for rs158477 remaining significantly elevated in both sexes (chromosome 16 empirical $\text{p-value}_{rs158477,male}$ = 0.04, chromosome 16 empirical $\text{p-value}_{rs158477,female}$ = 0.01, Supplementary Figure 3b,c). This suggests that the LD block around rs1967309 is differentiated between males and females in the Peruvians from 1000G. However, we note that the null model for the $F_{ST}$ statistic underlying PBS assumes no difference in genotype frequencies between sex (ie. may not be the appropriate tool to address this specific question), and we cannot exclude the possibility of random sampling noise.

### Admixture analyses

Recent admixture and migration events can influence LRLD. If segments of the genome are particularly enriched for a specific ancestry, this could lead to inflated LRLD between these segments. Given that the Peruvian is an admixed population between individuals of Native American ancestry (mainly Andean) as well as of European ancestry (Supplementary Figure 2), we ran several analyses to establish whether our results at *ADCY9/CETP* can be explained by admixture patterns.

#### 1. Local ancestry inference pre-processing

The reference populations used to run RFMix were YRI for the African ancestry, CEU for the European, CHB for the Asian from 1000G, subpopulations in NAGD (Northern American, Central American and Andean) for the Native American ancestry. We estimated local ancestry with RFMix on PEL from 1000G and LIMAA individuals.

For all 1000G populations (YRI, CEU, CHB, PEL), NAGD (Northern American, Central American and Andean) and LIMAA cohort, from the pre-processed datasets (see above) we kept only biallelic SNPs positions, removed SNPs with a MAF under 1% for each subpopulation, with more than 1% of missing individuals, with Hardy-Weinberg equilibrium p-value $< 10^{-4}$ with mid-adjustment using PLINK. We kept overlapping positions between all datasets and extracted the minor allele frequencies for each reference group. To avoid overlapping positions on the genetic maps, when SNPs had the exact same genetic position, we selected the SNP with the higher variance in allele frequencies (using var in R) between the reference groups (all subpopulations except PEL and LIMAA), keeping between 6,742 and 57,238 SNPs per chromosome for RFMix analysis.

### 2. Assessing proportions of global Andean ancestry

To see if there could be a potential enrichment or depletion of Andean ancestry at *CETP* and *ADCY9* loci compare to the rest of the genome, we looked at the proportion of attribution of Andean at those loci compared to the overall distribution of all chromosomes. From the 584,797 positions used for RFMix on all chromosomes, 4,476 position intervals were given, and we calculated the proportion of Andean attribution for each interval, then calculated the 95% confidence interval (CI) for all chromosomes which is [0.43 - 0.75]. The proportion at *ADCY9* and *CETP* loci were 0.58 and 0.66 respectively, which suggests that the correlation between *ADCY9* and *CETP* loci is unlikely to be due to an enrichment or depletion of Andean ancestry at both loci. Results are similar when only considering chromosome 16 to calculate the 95% CI.

### 3. LRLD in the Andean population from NAGD

Another question is to assess if the association was already present in the non-admixed ancestral Andean population. If this is the case, the association cannot be explained by the random distribution of Andean segments across the Peruvian genome. We computed LRLD as described in Peruvians in the Andean population from NAGD and we found that the association between rs1967309 and rs158477 is also significant (*ADCY9/CETP* empirical p-value=0.04, Figure 3-figure supplement 1a,b, Supplementary Table 1). We note that, in this population, strong association signals with rs158477 are also seen at other SNPs in the *ADCY9* LD block region. This result provides convincing evidence that the results in PEL and LIMAA are not due to random distribution of admixed segments but rather might have been inherited from the Andean population, where it was already present, and is maintained since then by selection.

In the Andean population, the association between rs1967309 and rs158477 is not significant when we stratified by sex (Supplementary Table 1), but we still see significant association signals with rs158477 at other SNPs in *ADCY9* LD block in both sexes (Figure 4-figure supplement 3)

**Comparison between Peruvian cohorts**

To evaluate the genetic difference between Peruvian from 1000G and LIMAA, we performed a PCA starting from the phased data files. We kept only biallelic SNPs with a MAF higher than 5% in each cohort and kept only positions with no missing genotype. We followed the suggestion given by flashPCA2 (https://github.com/gabraham/flashpca) (13), remaining 18,345 SNPs for the PCA. We then did a UMAP on 50 PCs given by flashPCA2 using the UMAP package on R (default parameters) (Supplementary Figure 5). As seen in the UMAP analysis, population structure exists in LIMAA, and PEL samples are mainly part of the largest subgroup observed in Supplementary Figure 2e, which was kept for LIMAA analyses to remove any confounders linked to population subdivision (see below). Also, the LIMAA cohort was initially recruited as part of a tuberculosis study (14), but our PCA and UMAP analysis showed no separation according to disease state.

**Null distributions of LRLD**

To evaluate how likely it is to observe, specifically in the admixed Peruvian population, a genotype correlation of $r^2 = 0.08$ between SNPs that are approximately 53 Mb apart on the same chromosome like between rs1967309 and rs158477, we have used two approaches. The first one was specific to the two genes under study, *ADCY9* and *CETP,* and therefore controls for all genomic factors specific to these regions. We selected all SNPs with MAF>0.05 in the two genes, and computed $r^2$ values for all 37,802 pairs (461 SNPs in *ADCY9* and 82 SNPs in *CETP*), yielding a null distribution for the expected genetic correlation between these genes. We then compared our $r^2$ value for rs1967309 and rs158477 to this distribution, with its rank being reported as an empirical p-value. This is referred to in the Results section as "*ADCY9/CETP* empirical p-value".

This approach is appropriate to correct for the genomic context specific to our genes of interest, but does not account neither for allele frequencies (most SNPs in the null will be at lower frequencies than our two SNPs) nor for overall admixture levels in the genome of this sample, thus we used a second empirical approach to account for these important confounders. For this genome-wide null distribution of the LRLD matching our SNPs, we generated one set of pairs of SNPs and evaluated LRLD between these random pairs in both LIMAA cohort and PEL from 1000G. Since frequencies in the LIMAA cohort are likely better estimates of allele frequencies in the Peruvian population because of the size of the sample, we started our selection based on SNPs' characteristic in this cohort: we extracted pairs of biallelic SNPs from chromosome 1 to 18, (the other chromosomes being too small) with a MAF between 15% and 30%, separated by between 50-60 Mb and 61±10 cM based on the PEL genetic map from 1000G. If SNPs in a pair shared coordinates on the genetic map (in cM) with another SNP from another pair, we kept only one of these pairs. We ended up with 3,576 non-overlapping SNP pairs for calculating the LRLD null distribution matching our rs1967309-rs158477 pair obtained from the LIMAA cohort. For analysis in PEL from 1000G, we added an extra frequency filtering step to remove pairs for which one or both SNPs had a MAF below 5% in PEL, leaving 3,513 pairs for analysis for PEL of 1000G. To calculate an empirical p-value in PEL, we evaluated the number of pairs which had a LRLD value larger to the observed value for rs1967309-rs158477 and divided this number by the total number pairs (n=3,513). This is referred to in the Results section as "genome-wide empirical p-value".

From the 3,513 pairs of SNPs sampled to create the genome-wide null distribution in both sexes in PEL, we stratified by sex and recomputed null distributions for males and females in the same way as for the full cohorts, also with a MAF filter at 5%, leaving 3,505 pairs in males and 3,512 in females in PEL. In males, the $r^2$ value between rs1967309 and rs158477 was the highest of the distribution (genome-wide empirical p-value < $2.85 \times 10^{-4}$), but for females, it was in the 20th percentile (genome-wide empirical p-value = 0.80).

**Permutation analysis of sex-specific LRLD at the positions rs1967309 and rs158477**

A second null distribution was derived for evaluating if the LRLD difference between sex for the rs1967309-rs158477 pair was significant, given the significant LRLD observed at these loci. We permuted the sex labels within the cohort and split them into two random groups of 42 pseudo-males and pseudo-females, while making sure an equal number of real males and females (21 of each) are found in each random group, yielding a total of 919 unique random splits that respected these conditions for the 85 PEL individuals. For each iteration, we calculated LRLD between rs1967309-rs158477 for each group and computed the absolute difference between them. To calculate a p-value, we evaluated the number of iterations that had a LRLD difference of more than or equal to the observed difference for the rs1967309-rs158477 pair between true males and females. The true absolute difference in $r^2$ values between rs1967309 and rs158477 (0.346) is the third highest value in this null distribution (p-value=0.002) (Supplementary Figure 4c).

**Genotype association between rs1967309 and rs158477 in LIMAA**

In the LIMAA cohort, we performed a genotype association test using a $\chi^2$ test with four degrees of freedom ($\chi^2_4$) with a permutation scheme to obtain the p-values, as reported in (15), to control for the marginal one-locus genotype counts. To avoid the potential effects of population subdivision on LRLD (16,17), we only kept individuals in the largest, likely more homogeneous, group seen in the UMAP performed on the first 50 PCs with PEL from 1000G (Supplementary Figure 2e). Two smaller distinct groups were identified in the UMAP analysis and these individuals were excluded from our analysis (cross shaped individuals in Supplementary Figure 2e), leaving 3,243 individuals for analysis. The permutation scheme consists in permuting the rs1967309 values 1,000 times and computing the number $\chi_4^2$ values obtained by permutation that are higher than the observed value for the rs1967309/rs158477 pair. For LIMAA, the $\chi_4^2$ value is 82.0 (permutation p-value < 0.001). We then performed the same analysis by stratifying by sex, and obtained a $\chi_4^2$ value of 56.6 (permutation p-value = 0.001) in males and a $\chi_4^2$ value of 37.0 (permutation p-value = 0.017) in females. We note that performing this analysis in the full cohort of 3,509 individuals (without excluding individuals from subpopulations shown in Supplementary Figure 2e) yield very similar results (full cohort $\chi_4^2$=77.6, male $\chi_4^2$=56.5, female $\chi_4^2$=34.5). To assess which combination is driving the effect, we used an empirical combination-specific test: the p-value is obtained by breaking the real rs1967309-rs158477 genotype combinations by permuting rs1967309 genotypes and evaluating how many permutated samples show an enrichment of a specific genotype combination as large as in the real data. Interestingly, the combination driving the highly significant male effect is an excess of rs1967309-AA + rs158477-GG (combination-specific permutation p-value < 0.001), whereas in female, the result seems to be driven by rs1967309-AA + rs158477-AA (combination-specific permutation p-value = 0.014). These sex-specific genotypic effects could not be captured by a linear model and can explain why the $r^2$ value in LIMAA is smaller than in PEL. Additionally, we note that in both sexes (but mainly in males), the low-frequency rs1967309-GG + rs158477-GA combination is enriched in LIMAA (observed counts is 112 whereas expected according to allele frequencies at both loci is 59.7).

Finally, to evaluate the effect genome-wide, we calculated the $\chi^2_4$ for all 3,576 pairs from the above described genome-wide null distribution for LRLD, then compared these with the value obtained for the rs1967309/rs158477 pair, in all individuals, males and females of LIMAA. In all groups, the rs1967309/rs158477 $\chi^2_4$ values were in the top values (genome-wide empirical p-value$_{all}$ = 0.0003; genome-wide empirical p-value$_{males}$ = 0.002; genome-wide empirical p-value$_{females}$ = 0.001), meaning that the association is significant genome-wide, as found in PEL using $r^2$ (Figure 3d).

Despite lower power in 1000G PEL sample, we replicated the rs1967309-AA + rs158477-GG enrichment in males in PEL using a 2x2 $\chi^2$ test, comparing specifically the rs1967309-AA + rs158477-GG to the three other combinations (rs1967309-nonAA+rs158477-GG; rs1967309-AA+rs158477-nonGG; rs1967309-nonAA+rs158477-nonGG, permutation p-value = 0.018). The rs1967309-AA + rs158477-AA association seen in females does not replicate (permutation p-value = 0.51) possibly due to low sample size (observed counts is 1, expected counts is 0.66).

Age was available in the LIMAA cohort, enabling us to test whether the LRLD pattern is associated with age, which could suggest a survival benefit if the association is not seen at younger ages. No correlation was seen between genotype and age for rs1967309 and rs158477, and age distributions between males rs1967309-AA+rs158477-GG and females rs1967309-AA+rs158477-GG were not significantly different. Because sample size was large enough in this cohort to perform a stratified analysis, we further split the cohort into nearly balanced age categories in males (0-19 years old: 435; 20-25: 464, 26-35: 523; over 35: 519) to establish if the LRLD is present in a specific sub-group. To test if the enrichment of rs1967309-AA + rs158477-

GG in males varies between age group, we calculated the expected frequencies using the frequencies in all age combined in males only (using the whole sample allele frequencies did not change the results). First, we generated a 2x2 contingency table comparing rs1967309-AA + rs158477-GG versus the three others (see above), then calculated a $\chi^2$, then we used a permutation test permuting rs1967309 genotypes 1,000 times to assess statistical significance. The empirical p-values suggest differences between age groups (permutation p-value$_{0\text{-}19}$ = 0.12; permutation p-value$_{20\text{-}25}$ = 0.21; permutation p-value$_{26\text{-}35}$ = 0.01; permutation p-value$_{>35}$ = 0.04), with significant p-values in older groups only. We thus more formally tested if the association differs between age groups between 0 and 25 years old (n=899) and above 25 (n=1042): we performed a $\chi^2$ test based on a 2x2 contingency table using the chisq.test() function in R, comparing the rs1967309-AA + rs158477-GG versus all other genotypes for the 2 age groups, permutating age values 1,000 times to estimate an empirical p-value. There was no significant difference between age group ($\chi^2$ = 0.02, permutation p-value=0.88). Similar results were obtained when the number of individuals were balanced across the two groups (cut off of 26 years old) or when the four initial age groups were used.

Finally, we also considered how the imputation quality in LIMAA could affect the main result of LRLD, because of imputation from non-representative reference panel populations is known to be problematic. We recomputed the genotype correlation ($r^2$) in LIMAA with our two SNPs imputed with the TOPMED panel, a more representative panel than the Haplotype Reference Consortium initially used. The value obtained is 0.0047 compared to 0.0046 before, showing that imputation quality is unlikely to have affected our results.

### III. Expression data

#### *ADCY9* and *CETP* expression quantification from RNAseq data

By analysing more in depth the *ADCY9* gene and its isoforms, we noticed that a considerable proportion of reads were assigned to a specific isoform, *ADCY9-205* (ENST00000574721.1), a 2.4 Kb long retained intron that does not have a validated status. It was removed from the gene definition file (GTF) to remove any noise from spurious transcription. All GTEx data was therefore reprocessed for *ADCY9* and *CETP* by the same pipeline (see below) to obtain transcription levels per sample at the gene level, consistent for the two genes across cohorts.

For each eQTL dataset (GEUVADIS, GTEx, CARTaGENE), we recalculated the top 5 PCs using genotype data with flashPCA2. For duplicated samples, we kept the sample with the highest read count in the library and removed samples which had a total of less than 10 million of reads. We trimmed the sequencing reads from Illumina adaptors and bad quality ends (BQ>20) using TrimGalore!. We mapped the alignment files on Hg38 human genome reference using STAR v2.6.1a (18) with the Ensembl 87 genome annotation, then estimated count for each gene using RSEM v1.3.1 (19). For GTEx, we separated each tissue at this step, then removed tissues with less than 50 samples, leaving samples from 49 different tissues to avoid over-interpretation due to low sample size while maximizing the number of tissues to be tested. We kept the genes which had more than 6 reads in at least 20% of the sample. We then normalized expression data using limma (TMM normalization) (20) and voom (21). We calculated PEER factors (22) on the normalized expressions. For all sex-stratified analysis, we kept sex-stratified tissues that had at least 50 samples, and recomputed PEER factors with samples from only one sex (which we term sPEER factors).

To test if *ADCY9* and *CETP* expression is correlated across tissues in humans, we used data GTEx and performed a linear regression correcting for the first 5 PCs, age, sex, the collection site (SMCENTER), the sequencing platform (SMGEBTCHT) and total ischemic time (TRISCHD). We find that *ADCY9* and *CETP* gene expression levels are negatively correlated (and significantly($p<0.05$) so for Adipose-Subcutaneous, Adrenal Gland, Artery Coronary, Artery Tibial, Brain-Cerebellar Hemisphere/Cerebellum/Cortex/Frontal Cortex/Putamen (basal ganglia), Breast-Mammary Tissue, Esophagus-Gastro esophageal Junction/Muscularis, Heart-Left Ventricle, Lung, Prostate, Small Intestine-Terminal, Spleen, Stomach, Uterus, Whole blood), except in skin tissues and cells cultured fibroblast, for which a significant positive correlation is found (Supplementary Figure 6).

**Expression Quantitative Trait Loci (eQTL) analysis for rs1967309 and rs158477**

We first looked at the effects of the SNPs independently on their respective genes. The covariates include the first 5 PCs, age (except for GEUVADIS, information not available), sex, as well as PEER factors, calculated to take into account hidden factors. In GTEx, we added additional covariates: the collection site (SMCENTER), the sequencing platform (SMGEBTCHT) and total ischemic time (TRISCHD). One limitation is there is no standardized way of deciding how many PEER factors to include. We tested the robustness of results to the inclusion of different numbers of PEER factors in the models and we report them all for GEUVADIS, CARTaGENE (CaG) and GTEx for transparency (Supplementary Figure 7-9). The maximum number of PEER factors considered follows recommendation by GTEx based on sample size for each tissue.

SNP rs1967309 is a *cis* eQTL of *ADCY9* in whole blood in CARTaGENE (p-value=4.46 x $10^{-13}$, ß= -0.10, N=728, 10 PEER factors) with AA individuals having increased *ADCY9* expression compared to GG individuals. This effect is replicated in whole blood samples from GTEx (N=559), and several other tissues in GTEx (with esophagus being the most significant), but some tissues show an inversion of the direction of the effect, such as lung and thyroid. These results may differ from GTEx reported eQTL results, because of the removal of ADCY9-205 isoform (see above), the different expression normalization method and filters by ethnicity applied here. SNP rs158477 is found as a *cis* eQTL of *CETP* in GEUVADIS (p-value =1.69 x $10^{-4}$, ß=0.26) lymphoblastoid cell lines, replicated in GTEx (EBV transformed lymphocytes) as well as in many other GTEx tissues. Tissues with p-value below 0.05 across most PEER factors values (if not all) are: adipose tissues, hippocampus, liver, lung, small intestine, muscle, stomach, thyroid, with GG genotype having consistently less *CETP* expression than AA.

We next tested whether the SNPs are *trans* eQTL for the genes. We found nominally significant associations between rs1967309 and *CETP* expression in the ovary (p-value=0.0017, N=138, max PEER factors = 15) and hippocampus (p-value=0.049, N=150, max PEER factors = 30), results that are stable across PEER factors values, two tissues for which rs1967309 is not significantly associated with *ADCY9* expression (p-value>0.05). We found nominally significant associations between rs158477 and *ADCY9* expression in the brain-cerebellar hemisphere and liver.

We next evaluated the interaction effect between rs1967309 and rs158477 on gene expression levels, despite somewhat low statistical power, especially given the small number of samples by tissue. Because the appropriate value of the number of PEER factors to be included on the model is not obvious, we report all values of PEER until the maximum suggested by GTEx for the GTEx tissues (15 for N<150, 30 for 150≤ N<250, 45 for 250≤ N<350, and 60 for N≥350). We also required that the interaction term rs1967309*rs158477 for a tissue had p-values under 0.1 for a majority of values of the number of PEER factors included, to qualify a result to be suggestive of an interaction effect. In the GEUVADIS dataset, which has 287 samples, the interaction was significant on *CETP* expression and stable across PEER factors (Supplementary Figure 7a), which could mean that the effect of a SNP could be modulated by the other SNP. To evaluate this effect further and make sure this is not due to outlier effects or other statistical flukes, we stratified by genotypes of each SNP and investigated the effect of the other SNP on *CETP* expression. We used a linear regression with the same covariates mentioned above. We first stratified by the genotype of rs1967309, then evaluated the effect of rs158477 on *CETP* expression. SNP rs158477 is significant in the AA of rs1967309 (p-value=0.03, ß=0.45, OR=[1.05-2.36], n=46) and AG (p-value=0.009, ß=0.24, OR=[1.07-1.53], n=143), but not for GG (p-value=0.58, ß =0.07, OR=[0.83-1.40], n=96) (Figure 5b), potentially showing a mitigation of the eQTL effect of rs158477 for each alternative allele of rs1967309 on *CETP* expression. We also evaluated the effect of rs1967309 when we stratified by rs158477 on *CETP* expression. SNP rs1967309 is significant only for GG of rs158477 (p-value=0.05, ß=0.3, OR=[1.00-1.83], n=72), and not for GA (p-value=0.51, ß=-0.06, OR=[0.77-1.14], n=139) nor AA (p-value=0.68, ß=-0.07, OR=[0.66-1.30], n=74). The second dataset that we used was the GTEx, in which we evaluated 49 tissues. Since the effects across tissues are likely not independent, we did not correct for multiple testing, keeping a suggestive threshold at 0.10 and a significant threshold at 0.05, but those values need to be reached for a majority values of the number of PEER factors included to be convincing. Among the 49 tissues (Supplementary Figure 8), those with p-values under 0.10 for several numbers of PEER factors are hippocampus (N=150), hypothalamus (N=156), brain spinal cord (cervical c-1)

(N=114), substantia nigra (N=100) and skin sun-exposed (N=507). Among those tissues, rs1967309 is only a cis-eQTL of *ADCY9* in the substantia nigra.

Since the selective pressure differ between sexes, we stratified our expression analysis by sex. For *CETP* expression analysis, there are no consistent signals for GEUVADIS, possibly reflecting lack of power or that there is no sex-specific effect in lymphoblastoid cell lines (Supplementary Figure 7c). However, in CaG, the significant interaction found is present only in male (again for the genotypic coding, Supplementary Figure 7d). In GTEx, we see significant interaction effects in males in tissues that had signals with sex-combined, such as brain hippocampus ($N_{male}$=105), hypothalamus ($N_{male}$=112) and spinal cord cervical ($N_{male}$=72), skin sun-exposed for CETP ($N_{male}$=330) (Figure 5d, Supplementary Figure 9). We note that for most brain tissues, the low sample size in females does not allow to conclude on the presence of an interaction effect in that sex. In these tissues, the direction of the effect in males is reversed compared to what is observed in GEUVADIS with sexes combined (Figure 5b), whereas the highly significant result in skin shows an effect consistent with the sex-combined GEUVADIS result (p-value=0.0017, ß=-0.32). More specifically, in the sun-exposed skin samples, in rs1967309 AA males, copies of the rs158477 A allele increase CETP expression by 0.49 (95% CI 0.12-0.87) on average. In rs1967309 AG males, the effect of rs158477 is null ($p\text{-value}_{AG}$=0.33) and the effect of the rs158477 A allele is suggestive in rs1967309 GG individual ($p\text{-value}_{GG}$=0.10) with a decrease of the CETP expression. Conversely, if we look at the effect of rs1967309 on *CETP* expression in skin sun-exposed when we stratified by rs158477 in males, SNP rs1967309 is neither a significant eQTL of *CETP* in GG of rs158477 in males (p=0.11, n=89) nor for GA (p=0.65, n=164), but is a significant eQTL for AA males (p=0.026, ß=-0.46, OR=[-0.87, -0.06], n=76).

We also identified new tissues where the interaction is either suggestive or significant in females, in artery tibial ($N_{female}$=156), heart atrial appendage ($N_{female}$=97), spleen ($N_{female}$=69) and stomach ($N_{female}$=105) (Supplementary Figure 9), with an effect reversed compared to the initial GEUVADIS result (Supplementary Figure 7a). For the pituitary tissue, it is significant in both sex (additive coding in female and genotypic coding in male, $N_{male}$=156, $N_{female}$=63), but the direction of the additive coding is reversed between sexes, possibly explaining why the sex-combined analysis did not show any signal. We note that the newly discovered signals are mainly for females, indicating that the signal was hidden by the male effects (or absence of effects), likely because of higher sample sizes.

### IV. Experiments

#### Real-time PCR quantification

Reverse transcription was performed from 500 ng total RNA in a 20 ml reaction using High-Capacity cDNA Reverse Transcription Kit (Applied Biosystems cat #4368814). RNA quantification was assessed using Agilent RNA 6000 Nano Kit for Bioanalyzer 2100 System (Agilent Technologies). Primers were designed using the Beacon designer software v.8 (Premier Biosoft) (Supplementary Table 3). The real-time PCR was carried out with SYBR-Green reaction mix (BioRad cat #1725274). The thermal cycling program was 3 min at 95$^{o}$C for initial denaturation followed by 40 cycles of denaturation for 10 sec at 95$^{o}$C, 30 sec annealing at 60$^{o}$C and 30 sec extension at 72$^{o}$C. qPCR assay was normalized with PGK1 and HBS1L genes.

**Western Blot analysis.**

200 ml of cell media from HepG2 transfected cells were concentrated using Amicon Ultra 0.5 ml 10 kDa cutoff units (cat #UFC501096) to 25 ml. Proteins were separated on 10% TGX-acrylamide gel. After O/N electrotransfer at 10 volts to PVDF membranes, CETP protein was determined using a primary anti-CETP rabbit monoclonal antibody (Abcam cat #ab157183) 1:1000 in 3% BSA, TBS, tween 20 0.5%, O/N 4$^{o}$C, followed by HRP-conjugated secondary antibody goat anti-rabbit 1:10 000 in 3% BSA 1h at room temperature. Detection was performed using Western Lightning ECL Pro (Perkin Elmer cat #NEL122001EA). Proteins levels were normalized with total proteins loaded.

### V. Phenotype associations

**Two-way and three-way interaction models in UK Biobank**

With rs1967309 coded under the genotypic model, allowing to capture non-additive effects, we tested if the effect of the interaction term was significant for a phenotype $Y$ using a likelihood ratio test (LRT) by comparing the following models:

$$Y \sim rs158477 + rs1967309 + sex + age + PC(1-5) \quad (m1)$$

$$Y \sim rs158477 * rs1967309 + sex + age + PC(1-5) \quad (m2)$$

$$Y \sim rs158477 * rs1967309 * sex + age + PC(1-5) \quad (m3)$$

We used the R function glm with family = "binomial" and compared models using the following: anova(a, b, test = "LRT"), with a=*m1* and b=*m2* to test for two-way interaction effects, and a=*m2* and b=*m3* to test for three-way interaction effects.

Individually, SNP rs1967309 ($f_A$=39%) is nominally associated with heart rate, and rs158477 ($f_G$=47%) with the systolic blood pressure (Figure 6-figure supplement 1), both results being mainly driven by association in females. Both SNPs are nominally associated with waist-hip ratio, rs1967309 in females, rs158477 in males. None of these effects are genome-wide significant.

**Phenotype associations in GTEx**

In GTEx, we had the variable MHHRTATT (phv00169162.v8.p2) for cardiovascular disease. This variable is defined as *Heart attack, acute myocardial infarction, acute coronary syndrome*. GTEx also has phenotypes, including cardiovascular traits. The variable DTHFUCOD (First Underlying Cause Of Death) was used to identify individuals whose cause of death included *Heart Attack/Stroke, Heart Disease, Acute Myocardial Infarction, Possible MI*, who were considered as cases. From the 699 samples kept, we excluded 6 for which the phenotype was missing or unknown for MHHRTATT variable and for which the cause of death was unrelated to heart disease, yielding a total of 130 cases and 563 controls. We added as covariates: sex, age and top 5 PCs. For this phenotype, neither rs1967309 nor rs158477 are associated with those phenotypes when taken alone. However, the interaction and both SNPs in the equation are significant (p-value≤0.05) or close to be significant (p-value≤0.10) for the phenotype ($p\text{-value}_{MHHRATT}$=0.01, $Estimate_{MHHRATT}$=-0.54) (Figure 6-figure supplement 1). Like for *CETP* expression, this means that for each G allele for rs1967309, there is a decrease of the effect of rs158477 on cardiovascular outcome. A difference with *CETP*'s expression is that there is an inversion of the direction of effect. In other word, for AA of rs1967309, directions of the effect of rs158477 are positive, with GG having less probability to have an event than AA

($Estimate_{MHHRATT}$=0.47), but for GG of rs1967309, estimates of rs158477 are negative ($Estimate_{MHHRATT}$=-0.79). Those results are consistent with the direction of dalcetrapib pharmacogenomic analysis. Considering that the GG genotype of rs158477, with less *CETP*'s expression, is a proxy for dalcetrapib, which is an inhibition of CETP, the same gradient is present for rs1967309. In AA of rs1967309, there is less heart disease with dalcetrapib (23). In GG of rs1967309, there is more heart disease with dalcetrapib. More study of this interaction is needed to understand the mechanism. However, this could lead to new insights into the potential biological mechanism behind the pharmacogenomic association involving the gene *ADCY9* with cardiovascular outcome of dalcetrapib.
